# Metabolic Symbiosis between Oxygenated and Hypoxic Tumour Cells: An Agent-based Modelling Study

**DOI:** 10.1101/2023.07.24.550289

**Authors:** PG Jayathilake, P Victori, CE Pavillet, D Voukantsis, A Miar, A Arora, AL Harris, KJ Morten, FM Buffa

## Abstract

Deregulated metabolism is one of the hallmarks of cancer. It is well-known that tumour cells tend to metabolize glucose via glycolysis even when oxygen is available and mitochondrial respiration is functional. However, the lower energy efficiency of aerobic glycolysis with respect to mitochondrial respiration makes this behaviour, namely the Warburg effect, counter-intuitive, although it has now been recognized as a source of anabolic precursors. On the other hand, there is evidence that oxygenated tumour cells could be fuelled by exogenous lactate produced from glycolysis, and that glycolytic tumour cells export lactate to the environment. Using a multi-scale approach combining multi-agent modelling, diffusion-reaction and stoichiometric equations, we simulated these cell populations and their environment, and studied the metabolic co-operation between cells exposed to different oxygen concentration and nutrient abundance. The results show that the metabolic cooperation between cells reduces the depletion of environmental glucose, resulting in an overall advantage of using aerobic glycolysis. In addition, the environmental oxygen level was found to be decreased by the symbiosis, promoting a further shift towards anaerobic glycolysis. However, the oxygenated and hypoxic cell populations may gradually reach quasi-equilibrium. A sensitivity analysis based on Latin hypercube sampling and partial rank correlation shows that the symbiotic dynamics depends on properties of the specific cell such as the minimum glucose level needed for glycolysis. Our results suggest that strategies that block glucose transporters may be more effective than those blocking lactate intake transporters to reduce tumour growth by blocking lactate production.

**AUTHOR SUMMARY:** Metabolic alteration is one of the hallmarks of cancer and the well-known metabolic alteration of tumour cells is that cells prefer to do glycolysis over mitochondrial respiration even under well-oxygenated and functional mitochondrial conditions. On the other hand, there is evidence that oxygenated tumour cells could be fuelled by exogenous lactate produced from hypoxic glycolytic cells in which it can create a metabolic co-operation between oxygenated and hypoxic cell populations. This metabolic co-operation could allow tumour cells to economically share oxygen and glucose and promote tumour survival. Using a multi-scale approach combining multi-agent modelling, diffusion-reaction and stoichiometric equations, we studied this metabolic co-operation between different populations of cells, exposed to a changing microenvironment. We predict that the tumour environmental glucose depletion is decreased while the oxygen depletion is increased by this metabolic symbiosis, promoting a further shift towards glycolysis. Our results also show that blocking glucose transporters could be more effective than blocking lactate intake transporters, because the former would disrupt both glycolysis and lactate production, drastically reducing tumour growth.

## 1. INTRODUCTION

Emergence of metabolic pathways played a vital role in cell evolution **[1]**.Among metabolic pathways, glucose metabolism is one of the basic survival metabolic pathways of human cells. Glucose is imported from the extracellular environment through glucose transporters (GLUT) of which GLUT1, GLUT2, GLUT3 and GLUT4 are best characterized. In the presence of oxygen, healthy cells usually convert glucose into pyruvate and then pyruvate is converted to acetyl-CoA. The acetyl-CoA is oxidized in the mitochondria in the tricarboxylic acid (TCA) cycle **[2]**. This aerobic respiration or oxidative phosphorylation (OXPHOS) can produce about 28-36 ATP molecules per glucose molecule **[3, 4]**. Under hypoxic/anoxic conditions (lack or absence of oxygen), as it often occurs in cancer, cells are not able to produce mitochondrial ATP and instead, they may use glycolytic ATP production. This only yields two ATP molecules per every glucose and lactate molecules. However, this respiration pathway is also used in presence of oxygen **[3]**. This use of glycolysis in aerobic conditions is known as the Warburg effect and it is one of the hallmarks of cancer cells **[5]**. The pyruvate produced by glycolysis is converted to lactate and then lactate and protons (H+ ion) are exported to the extracellular environment through membrane proteins called monocarboxylate transporters (MCT) which help to maintain the alkaline pH level inside tumour cells **[6]**. Glycolysis is an inefficient way to produce ATP and therefore more glucose is needed to maintain a sufficient ATP production rate for cell proliferation. When there is a low glucose concentration in the medium, cancer cells use lactate instead of glucose as their energy source **[6, 7]**. Lactate is imported through MCT transporters (in addition to their function as lactate exporters) and the imported lactate is then converted back to pyruvate that can be oxidized in the mitochondria producing far more ATP molecules. This is the so-called “reverse Warburg effect”, one of the adaptive metabolic mechanisms of tumour cells **[8, 9]**. Vascular tumours tend to have oxygenated cells in the tumour boundary, near vessels, and hypoxic cells in distant regions, thus this metabolic reprogramming could induce a metabolic symbiosis between lactate-fuelled oxygenated OXPHOS cells and glucose-fuelled hypoxic glycolytic cells **[6, 10]**, which, in turn, could result in a beneficial metabolic cooperation **[7, 10–14]** (**Fig 1**). Therefore, inhibition of lactate consumption by cancer cells could be an effective therapeutic strategy **[10, 15]**. On the other hand, lactate accumulation could cause higher intracellular acidity and lactate can also inhibit pyruvate dehydrogenase (PDH) and activate HIF.

**Fig 1.**
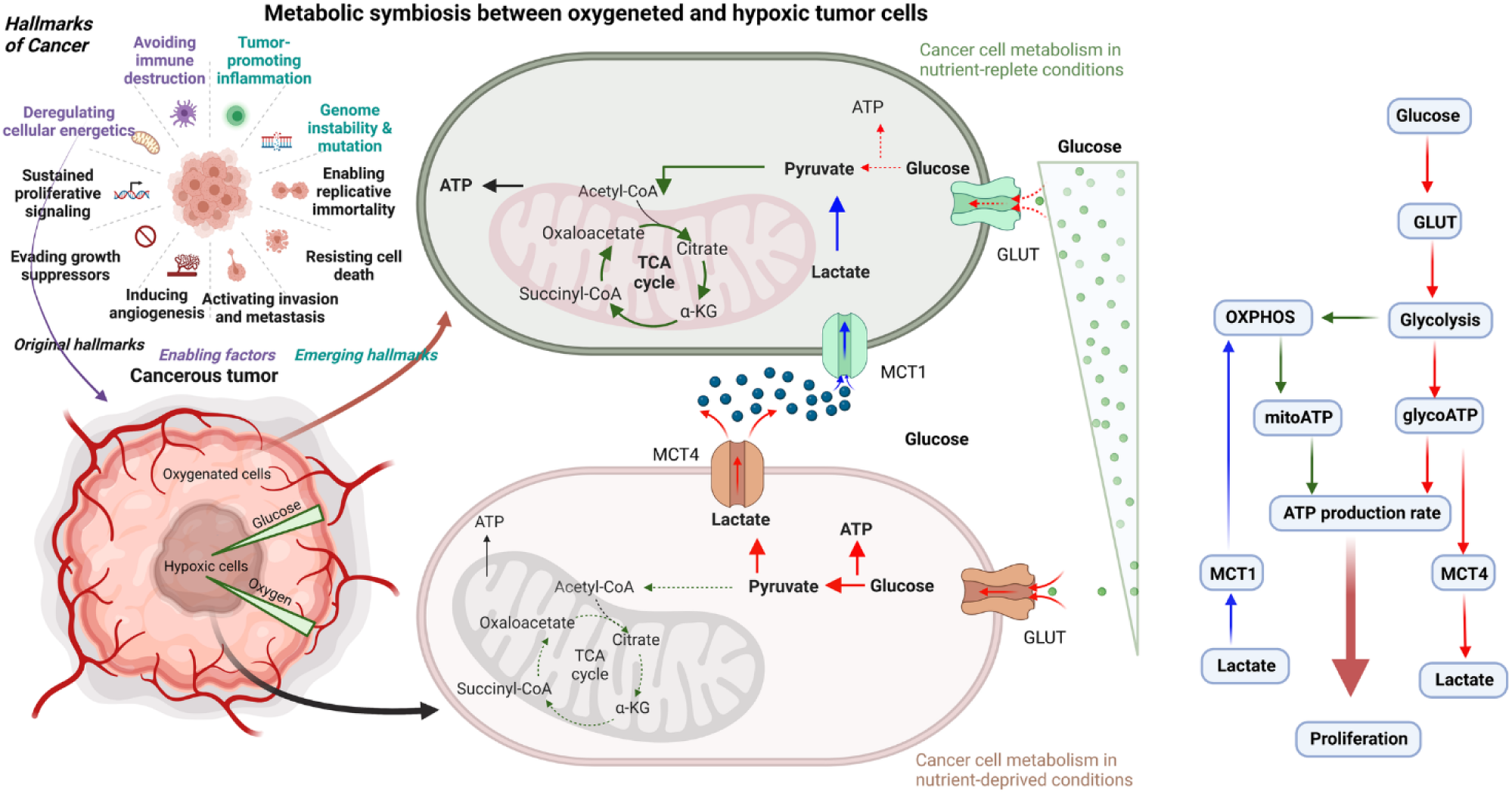
Metabolic symbiosis in tumour: Schematic diagram and flow chart illustrating the metabolic symbiosis mechanism between oxygenated and hypoxic tumour cells. Oxygenated cells at the tumour boundary (shown as green cells) consume exogenous lactate via MCT1 transporters and undergo lactate metabolism through OXPHOS. Inner hypoxic cells (shown as brown cells) consume glucose through GLUT transporters and undergo glycolysis and release lactate into the tumour microenvironment using MCT4 transporters. Metabolic symbiosis between the two cell populations (green cells and brown cells) helps hypoxic cells increase their glucose uptake, thereby helping tumour cells survive under low glucose conditions (created with BioRender.com).

Building on this knowledge, we investigated the possible beneficial effects of this potential synergy on the growth of the whole tumour and what factors influence the metabolic cooperation. Specifically, we asked:

i. How would the metabolic cooperation between hypoxic and oxygenated cancer cells affect nutrient levels in the microenvironment?
ii. Does this metabolic cooperation lead to increased tumour growth, overall and for specific cell populations?
iii. How is this cooperation affected by changes in environmental conditions (levels of oxygen, glucose, lactate etc.), cell spatial competition, heterogeneous somatic mutations, and continuous fluctuation versus steady state of nutrient levels in the microenvironment?
iv. What are the best strategies to disrupt this metabolic symbiosis?

## 2. MULTI-SCALE MODELLING APPROACH COMBINING MULTI-AGENT MODELS, DIFFUSION-REACTION & STOICHIOMETRIC EQUATIONS

We needed a modelling framework suitable to study the interactions between heterogeneous cell populations, including the effect of perturbing sub-cellular gene networks (i.e., cell regulatory networks) and the cellular microenvironment. Agent-based modelling (ABM) is a useful methodology to study ecological problems of this kind and identify emerging behaviours of heterogeneous populations **[16–20]**. Indeed, ABM models have been employed by us and others to study how tumours respond to drugs **[19, 21–23]**, the impact of environmental conditions **[22, 24]**, cell metabolism **[3, 4, 25, 26]** and cell competition **[27, 28]**. However, most of these models have assumed homogeneous environments and populations and have none or extremely simplified sub-cellular molecular interactions. To investigate emerging behaviours from heterogeneous utilization of cellular pathways, we adopted a recently proposed multi-scale agent-based framework that enabled us to model cancer cells, the microenvironment surrounding them (including nutrients and oxygen), and their respective gene regulatory networks **[16]**. The model was developed building on the widely adopted modelling platform NetLogo **[29]**, with previous **[16]** and new gene network and spatial functionalities developed by our laboratory as described below. **Fig 2A** shows the basic components of our model, and **Fig 2B** demonstrates how different spatial scales communicate with each other. More detailed descriptions of the model are given in **S1 Text**. The intra-cellular scale represents the cell’s gene regulatory network, which is a mitogen-activated protein kinase (MAPK) network **[30]**, driving the growth of cancer cells (**Fig S2**). Additionally, we modelled metabolic pathways including cellular respiration, glucose and lactate metabolism, and resulting ATP production. Specifically, glucose is imported through GLUT1 transporters (Note that not all GLUT isoforms are included here and GLUT1 is highly expressed in breast cancer samples as shown in **Fig S4**) and is converted to pyruvate at the end of the glycolysis process (**Figs 1** and **S2)**. The pyruvate can go through OXPHOS, which produces mitochondrial ATP (mitoATP) or can be converted into lactate resulting in only glycolytic ATP (glycoATP). Both mitoATP and glycoATP will determine the ATP production rate, and if the ATP production rate is above a certain threshold, that is 80% of the maximum possible ATP production rate **[4]**, the cell will be able to proliferate – subject to inhibition by other parts of the regulatory network. When cells do not use mitochondrial respiration, the pyruvate will be converted to lactate and then it will be exported to the environment through MCT4 membrane proteins. When there is enough lactic acid and oxygen in the medium, cells will import lactic acid through MCT1 membrane proteins. The lactate will be converted back into pyruvate and pyruvate will go through OXPHOS producing mitochondrial ATP **[12, 31, 32]**. To cover cellular functions including proliferation, apoptosis, growth arrest, cellular response to growth factors and the hypoxic microenvironment, we modelled the links between glucose/lactate metabolism and a previously developed MAPK-HIF network **[16, 30]**. To model Boolean logical conditions for the metabolic network nodes and the nodes linking with the existing MAPK-HIF network, supporting evidence was used from the literature (**Table S1)**.

**Fig 2.**
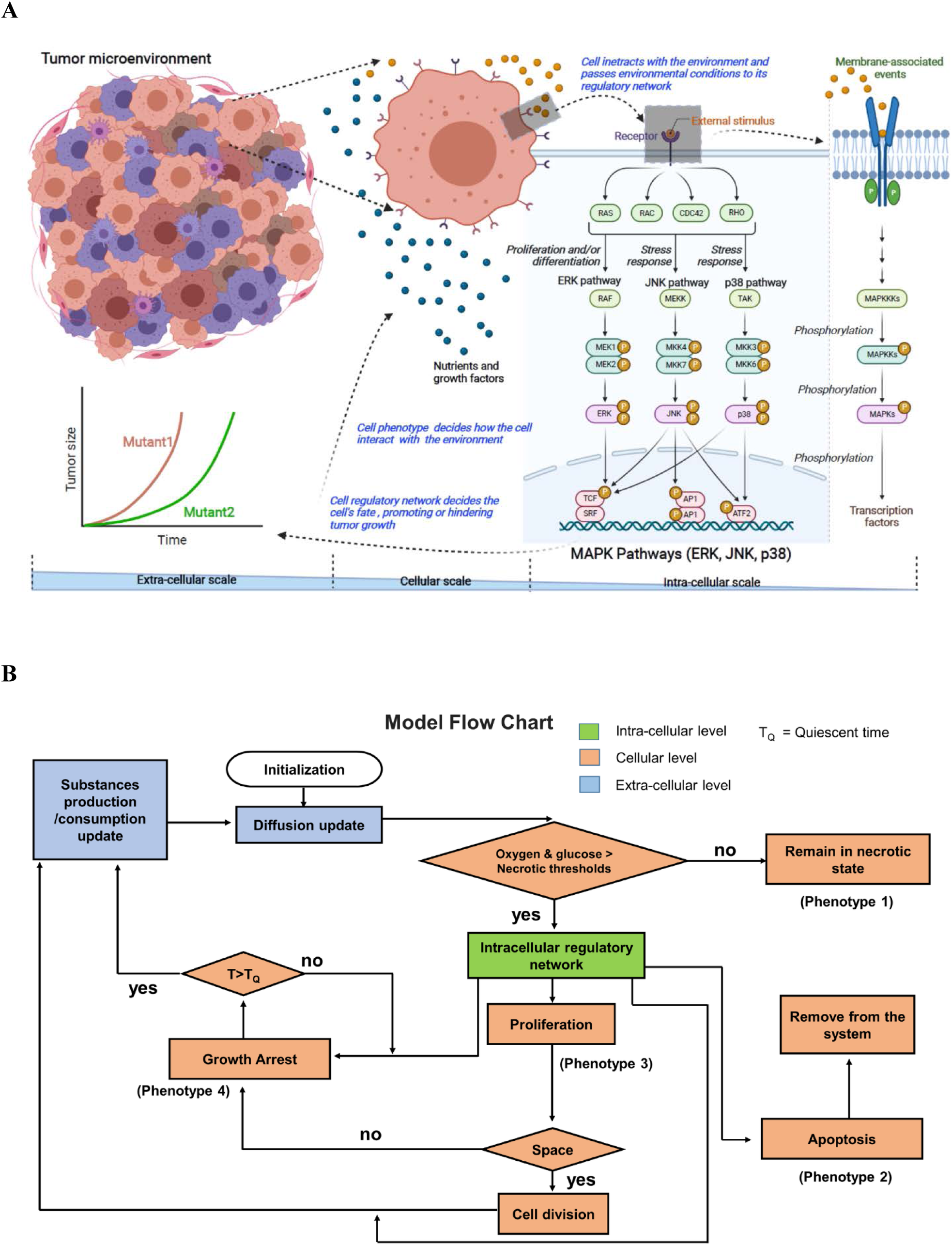
Agent-based mathematical model: (A). The model is a multi-scale agent-based model implemented using the NetLogo platform. The extra-cellular scale models gradients of substances such as oxygen, glucose and lactate by using partial differential equations. The cellular scale models cell-cell interactions using a cellular automaton approach. The smallest scale, the intra-cellular scale, handles subcellular molecular interactions using a Boolean gene regulatory network. All scales communicate with each other and therefore the tumour growth is an emerging property of the sub-cellular molecular interactions. The model can be used to study tumour growth under different environmental conditions, gene alterations and heterogeneous cell populations (created with BioRender.com). (B). The model flow chart shows how the three different scales are connected. If the local oxygen and glucose levels are below their respective threshold values, the cell becomes necrotic. If the cell is not in the necrotic state, the intra-cellular regulatory network determines the cell fate which is Proliferation or Apoptosis or Growth Arrest. An apoptotic cell is removed from the simulation immediately. A proliferative cell can divide if there is empty space nearby, otherwise it can switch to growth arrest state and waits a TQ time before checking environmental conditions again to find its new phenotype. The cell phenotype can influence the gradients of the diffusible substances on the microenvironment through producing/consuming diffusible substances based on its phenotype. The altered environmental properties are fed to the intra-cellular network of the cell again through input nodes and then the network can decide its new phenotype according to altered microenvironmental conditions.

In the cellular scale, each cell is represented by entities in a lattice. Gradients of diffusible substances in the microenvironment (glucose, oxygen, lactate etc.) are modelled in the extra-cellular scale. The soluble substances: oxygen, glucose, lactate and growth factors (GFs), are described by the diffusion-reaction equation as given below.

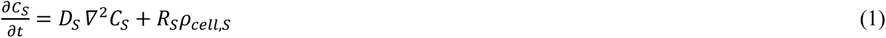

where *C*, *D* and *R* are the substance concentration, its diffusion coefficient and its rate of consumption or production, respectively. *ρ_cell,S_* represents the density of cells which consume or produce substance *S*. Typically, the time scale of substance diffusion (order of seconds) is much smaller than that of cell phenotype changes (order of hours) and therefore the diffusion is assumed to be in a steady state when the cell phenotype is updated. Cell necrosis occurs when both oxygen and glucose levels are lower than their respective critical values, as previously used **[20]**. If proliferative cells do not have enough neighbouring space to divide, those cells will switch to a quiescent state until they have some empty space to proliferate, thus modelling contact inhibition **[33, 34]**. We used the model in two-dimensional space (2D) for the present study.

The sink term of Eqn (1) (R) is described in accordance with the stoichiometry of oxygen, glucose, and lactate in their respective reaction equations **[3, 4, 27]**. Therefore, the oxygen consumption rate *R_o_*_2_is modelled as 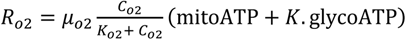, *K* = 0.5, in which *μ_o_*_2_ is the maximum oxygen consumption rate and *K_o_*_2_ is the half-saturation coefficient. mitoATP and glycoATP are dynamic variables and their status are provided by the executable cell regulatory network encapsulated inside each cell (**Figs S1** and **S2**), at each time step of the simulation. Specifically, when the cell uses glycolytic ATP production, glycoATP is 1 and otherwise it is 0. Similarly, if the cell uses mitochondrial ATP production, then mitoATP is 1 and otherwise it is 0. The value of *K.*glycoATP reflects the amount of oxygen consumption by a cell engaged in the glycolysis process. Here, we assume that even under glycolysis the tumour cells can consume a certain amount of oxygen for some other cellular processes, such as for example macromolecule synthesis **[35]**.

The glucose consumption rate can be calculated using the stoichiometry of the glucose oxidation equation given below.

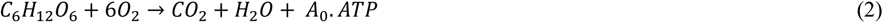

where, *A*_0_ is the ATP yield at the relevant oxygen and glucose abundance conditions, which is about 28 to 36 ATP molecules. The proportion of ATP production by glycolysis and OXPHOS depends on the cell type **[36]** and therefore, we assume that glycolytic and OXPHOS cells have similar ATP production rates for their proper functioning. The glucose consumption rate is then modelled as

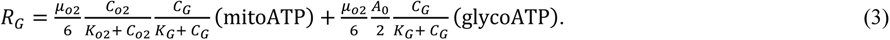

Taking different ATP production efficiencies of OXPHOS and glycolysis, the ATP production rate of a cell is then described as

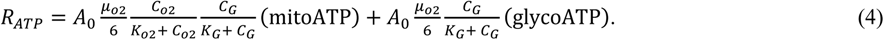

The glucose conversion into pyruvate in the glycolysis process can be written as

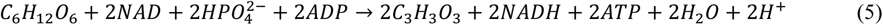

and then pyruvate conversion to lactate can be written as

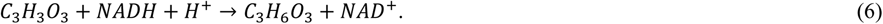

Eqn (5) and (6) show that one glucose molecule can produce two lactate molecules through the Warburg effect, and therefore the lactate production rate can be modelled as

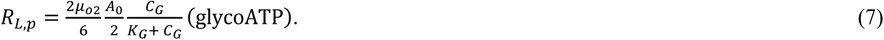

Considering that stoichiometry between glucose and oxygen is 1:6 for mitochondrial respiration (Eqn. 2), and glucose to pyruvate is 1:2 for glycolysis, it is assumed that lactate to oxygen ratio for reverse Warburg effect is 1:3. Therefore, the lactate consumption rate for the reverse Warburg effect is estimated as

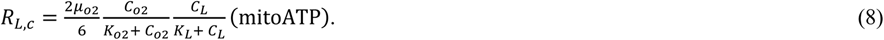

The glycolysis pathway can produce two protons (H+) per glucose molecule and hence the proton production rate is modelled as

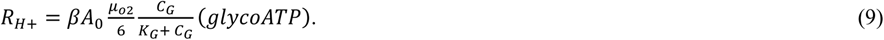

Here, β < 1 is the proton buffering coefficient of the tumour microenvironment **[4]** and it is chosen as 0.001 for this model because it results in a realistic pH level in the tumour microenvironment. The local extracellular pH due to exported H+ ion is calculated as *pH* = −*log*10([*H*+]).

The consumption rate of growth factors and inhibitors are modelled as *R_S_* = *γ_S,C_C_S_* and the production rate of any growth factor is described as *R_S_* = *γ_S,P_*.

The diffusible substance (oxygen, glucose, growth factors etc.) concentrations are kept constant at the boundary of the computational domain. These boundary values and all other model parameters are given in **Table S3**. Not all the parameters are based on evidence from previous studies, and therefore some parameter values are assumed based on other relevant data. However, we ran ten replicates of each simulation to assess the effect of different initial conditions, and a sensitivity analysis based on Latin hypercube sampling and partial rank correlation **[37, 38]** was performed to ensure the correctness and robustness of all our conclusions.

The level of symbiosis is quantified by the metabolic symbiosis index (*MSI*) and its definition is motivated by **[31]** as below:

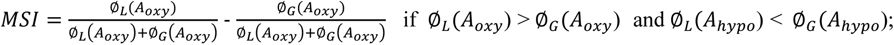

And *MSI* = 0 otherwise. Here, Ø*_L_*(*A*) and Ø*_G_*(*A*) are lactate metabolic and glycolytic cell fractions of the tumour region A, respectively. *A_oxy_* and *A_hypo_* are oxygenated and hypoxic regions of the tumour, respectively. The symbiosis index would vary from 0 to 1 depending on the strength of the symbiosis.

## 3. RESULTS & DISCUSSION

### 3.1. A gene network model of cellular metabolism and respiration

First, we used RNAseq data for multiple breast cancer cell lines grown under normoxic and hypoxic conditions (see **Fig S4** for results, **S1Text** for experimental conditions) to evaluate the coherence of our modelling assumptions. We observed that for cells under hypoxic conditions, the expression levels of SLC2A1 (or GLUT1), SLC16A3 (or MCT4) and LDHA increased, in agreement with our model of the Warburg effect and previous reports **[39–41]**. The increased expression of these genes supports our modelling hypothesis that hypoxic tumour cells transport more glucose, then to convert it into pyruvate, and finally the pyruvate would be metabolized to lactate and exported to the tumour microenvironment through MCT4 transporters as we have modelled in our regulatory network.

We then used the model with parameters setting in **Table S3** to investigate metabolic symbiosis between oxygenated and hypoxic cell populations. Our simulations showed metabolic symbiosis between hypoxic and oxygenated cells of the tumour when MCT1 was in wild type status (MCT1wt), while this symbiosis was lost when MCT1 was knocked out in our model. Of note, we model knockouts by setting a constrain on the status of the corresponding node in the gene regulatory network (MCT1) to inactive, for the duration of the current simulation. When a tumour grows with MCT1wt, we observe symbiosis, whereby the well-oxygenated tumour cells on the boundary of the simulated organoid switch to OXPHOS while hypoxic inner cells switch to glycolysis (**Fig 3A**). Importantly, glycolytic and OXPHOS cell populations co-exist under symbiosis, while the tumour growth mainly depends on the aerobic glycolytic population when symbiosis stops (**Figs 3B-C**). In the symbiotic tumour, the size of both cell populations is initially comparable, and the OXPHOS cell population dominates over the glycolytic cell population later when the glycolytic cells produce more lactate for OXPHOS cells **(Fig 3B)**. However, when MCT1 is inactive the symbiosis is lost and the OXPHOS cell population is about only 4.5% of the glycolytic cell population at the end of the simulation because OXPHOS is not driven by lactate **(Fig 3C)**. When symbiosis exists in the tumour, we observe that the cells at the boundary consume more oxygen for lactate metabolism, and therefore oxygen is depleted inside the tumour pushing to a more hypoxic environment (**Fig 3D**). However, when symbiosis stops, the emergence of a hypoxic cell population is delayed (**Fig 3E**). As the hypoxic population emerges when symbiosis exists in the tumour, the symbiosis index rapidly increases from zero to 1 indicating that metabolic cooperation between oxygenated and hypoxic tumour cells establishes at that time (**Fig 3F**). These simulation results clearly show that a tumour would gain a growth advantage due to metabolic symbiosis because symbiotic tumour cells can proliferate through either glycolysis (i.e., glucose to lactate) or OXPHOS (i.e., glucose/lactate to OXPHOS), or both pathways (**Fig 3G**). In support of the results from our simulations, in-vitro experiments have also shown that lactate metabolism can increase growth of human cancer cells in glucose-limited mediums of breast **[15]**, glioma **[31]** and colon **[10]** cancer. In-vitro studies of breast cancer cells have also shown that the presence of lactate would enable cells to withstand glucose-limited conditions **[15]**.

**Fig 3.**
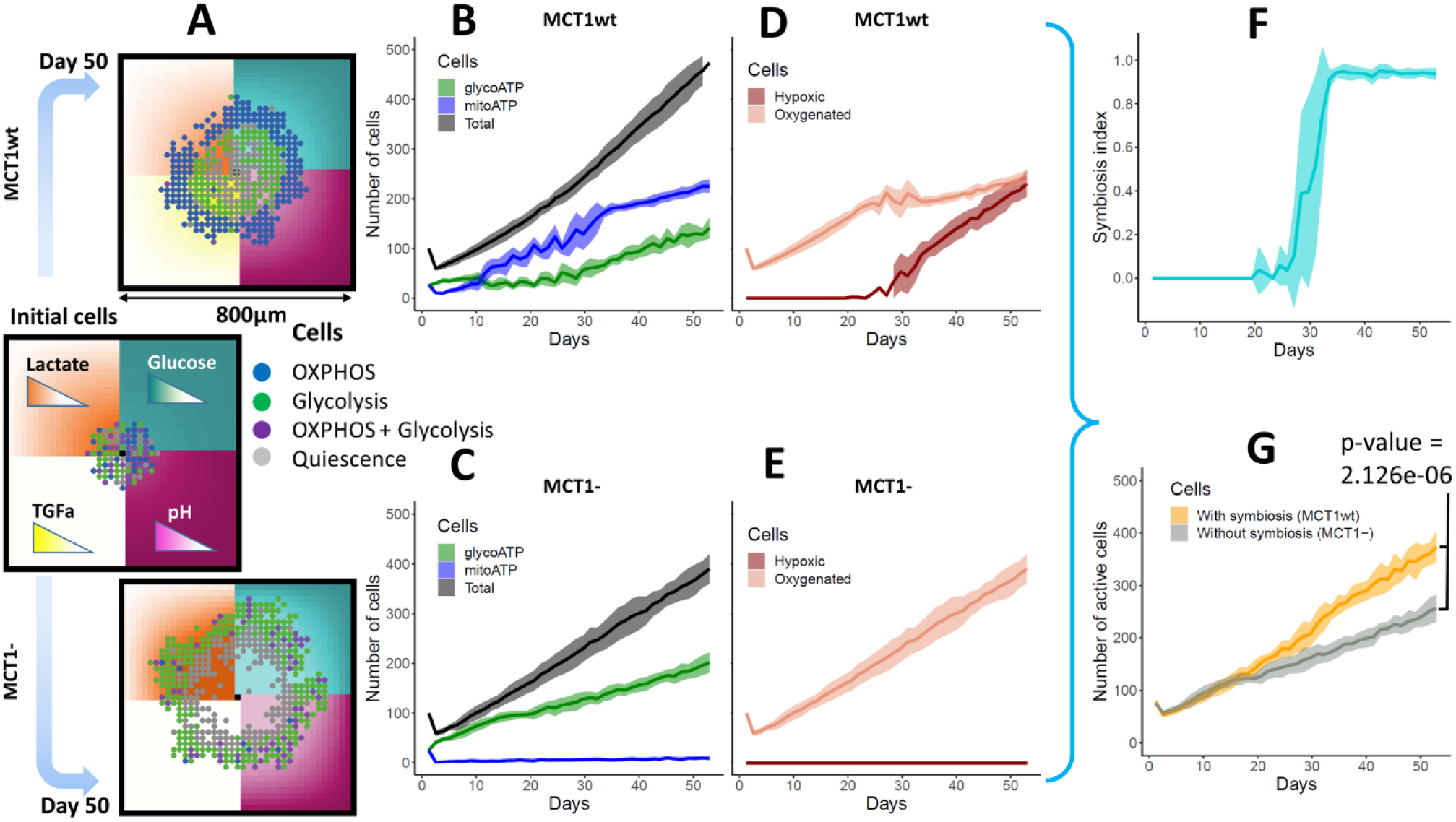
Tumour growth and symbiosis: **(A)**. If MCT1 is not mutated (MCT1wt), symbiosis is observed, while when MCT1 is mutated with loss of function (MCT1-), symbiosis is not observed. Different cell populations are shown: OXPHOS (blue cells, glucose/lactate--> pyruvate -->OXPHOS), Glycolysis (green cells, glucose-->pyruvate-->lactate), both OXPHOS and glycolysis (purple cells), not metabolically active or quiescence (gray cells). Gradients of glucose, lactate, tumour growth factor alpha (TGFA) and pH are shown on the microenvironment. **(B, C)**. Number of total, glycoATP and mitoATP cells are shown. The total number of cells contains all the cells in the tumour including dead cells. The mitoATP and glycoATP are the number of cells rely on OXPHOS and glycolysis for ATP production, respectively. **(D, E)**. Number of hypoxic and oxygenated tumour cells with MCT1 wild type and mutated condition are shown. The hypoxic cell population is not seen when symbiosis is lost (MCT1-) because oxygen cannot be depleted fast enough without lactate oxidation. **(F, G)**. Metabolic symbiosis index and number of active (viable) cells are shown when tumour grows with MCT1wt and MCT1-, so with and without symbiosis. The metabolic symbiosis index quantifies the strength of symbiosis between hypoxic and oxygenated tumour cells. The index can range from 0 to 1 depending on the symbiosis strength. The active cells are the cells with their metabolic pathways are active (i.e., blue, green and purple cells shown in A). The shaded area of curves shows respective standard deviation.

### 3.2. Metabolic symbiosis between oxygenated and hypoxic tumour cells depends on the status of genes that are commonly altered in cancer

We then asked if a similar scenario would be observed under a number of gene alterations that are commonly observed in cancer. Therefore, we examined gene aberration and expression patterns observed in the clinical setting for breast cancer samples taken from The Cancer Genome Atlas (TCGA) (see supporting materials for details). We specifically asked how our network genes are likely to either over or under-express in clinical samples. We took a conservative approach and considered the gene as over-expressed when the standard deviation was above +3 with respect to normal samples and under-expressed as when it was below −3. **Fig S5** shows the probability of aberrant gene expression (i.e., the percentage of samples that gene is over (**Fig S5A**) or under expressed (**Fig S5B**)). Of note, typically a greater number of genes would be considered as differentially expressed with respect to normal samples, but for the current study, we restricted the set of genes to the genes associated with our network. As can be seen in our results, in our network more genes tend to be under-expressed than over-expressed. Using a published protocol-sigQC **[42]** we also looked for a correlation among the absolute expression of these genes taken from TCGA (**Fig S6A**) and Cancer Cell Line Encyclopedia (CCLE) (**Fig S6B**) databases. Although some correlations could be observed, this was overall not strong with correlation coefficients found between −0.5 and +0.5, and many around 0. Thus, for the purpose of our simulations we assumed in first instance that the aberrant over- or under-expression of these genes occurred independently of each other (**Fig S6C**). While this might be improved in future studies, it allowed us to simplify the problem in first instance, and simulate each gene knockout or enrichment in our model as a single event rather than as linked coordinated events. Furthermore, TCGA and CCLE datasets (**Fig S6C**) showed similar distributions of sigQC metrics of variability and expression, suggesting that this genes set would be applicable to both cell lines and clinical samples. More details about these metrics can be found in **[42]** and are not repeated here.

Next, we perturbed the status of each of the network genes in our model and the model was run with MCT1wt and MCT1-conditions to investigate symbiosis and possible interaction between symbiosis and gene alterations commonly observed in cancer. We asked whether each of these gene alterations would affect the metabolic symbiosis-induced growth of the tumour **(Figs 4A-B),** symbiosis index **(Figs S7A-B)** and if there was a difference between the number of active (viable) cells in symbiotic and non-symbiotic tumours **(Figs 4C-D** and **Figs S7C-E)**. As we can see in **Figs 4A-B** and **Figs S7A-B**, some gene alterations give a growth advantage when there is symbiosis between hypoxic and normoxic cells (MDM2+, ERK+, MSK+, p38-, p53-, p14-etc.; here + and − indicate enriched and knockout status, respectively), while some other gene alterations reduce tumour growth under symbiosis (p14+, FOXO3+, GRB2+, RAS-, PLCG-, MYC- etc.). Symbiosis-induced growth advantage is seen at the later stage of tumour growth because it takes some time for the emergence of hypoxic cells in the tumour and then establish a cooperative interaction between two cell populations, approximately after 17 days (**Figs S7A-B**).

**Fig 4.**
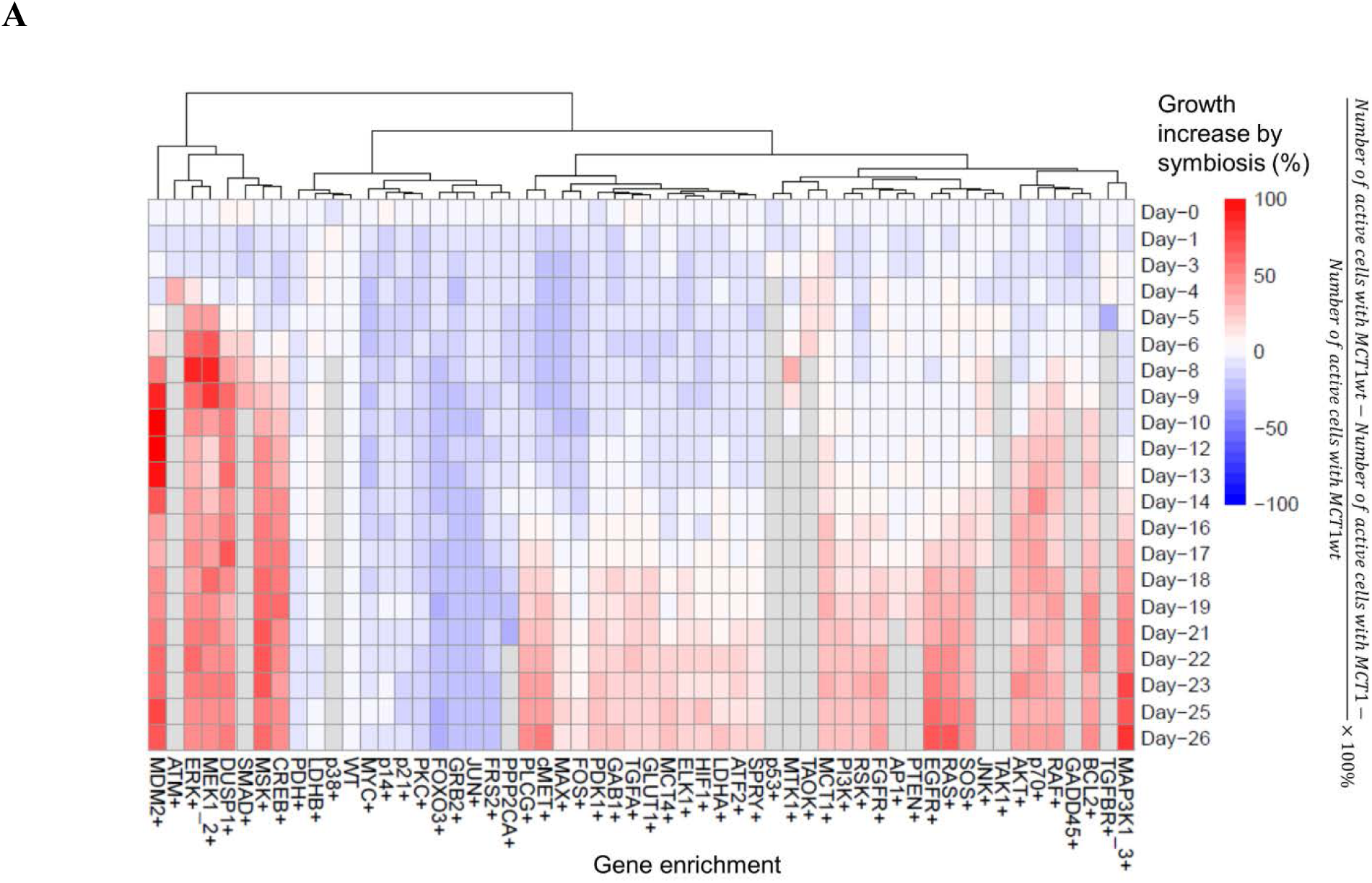
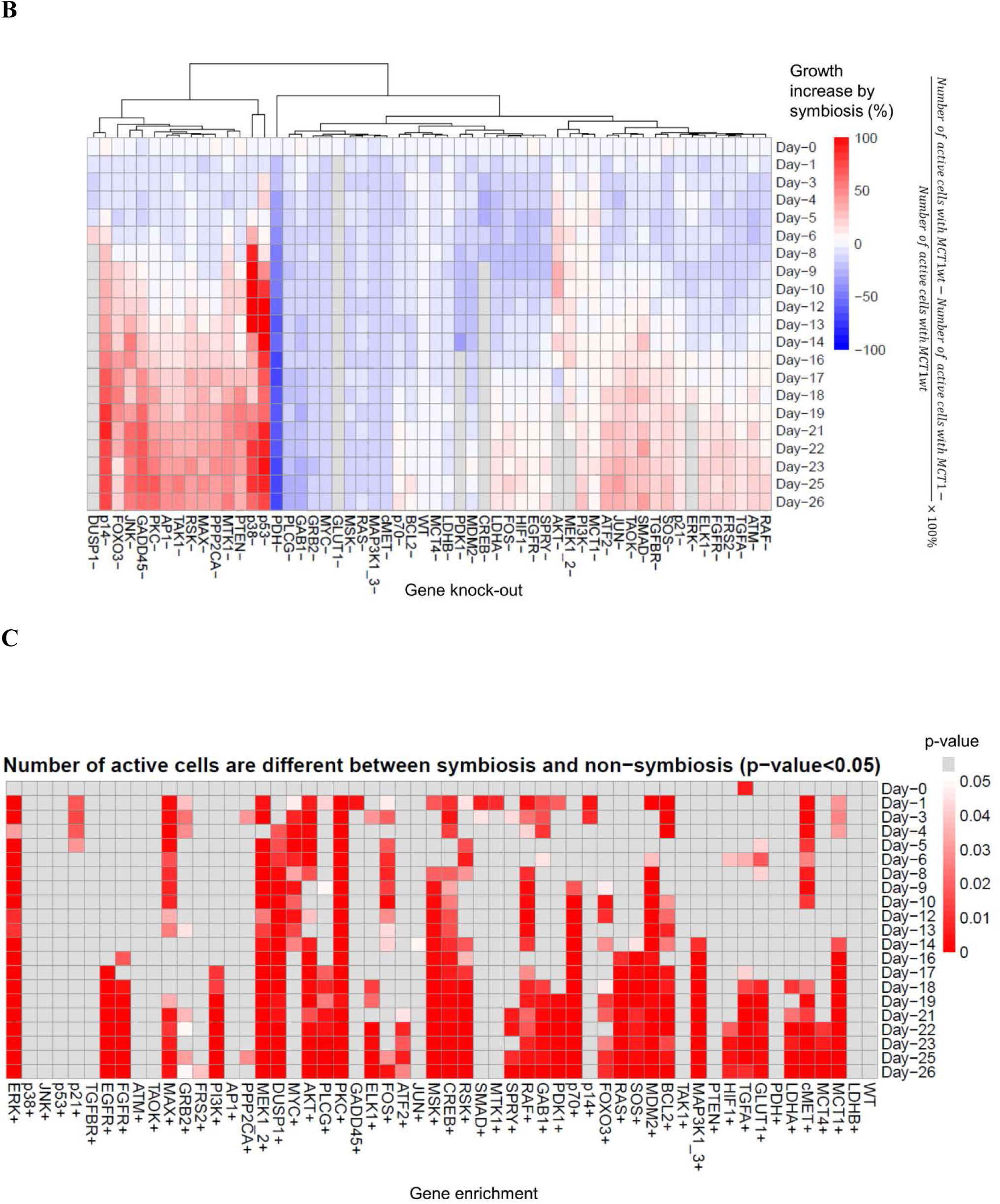
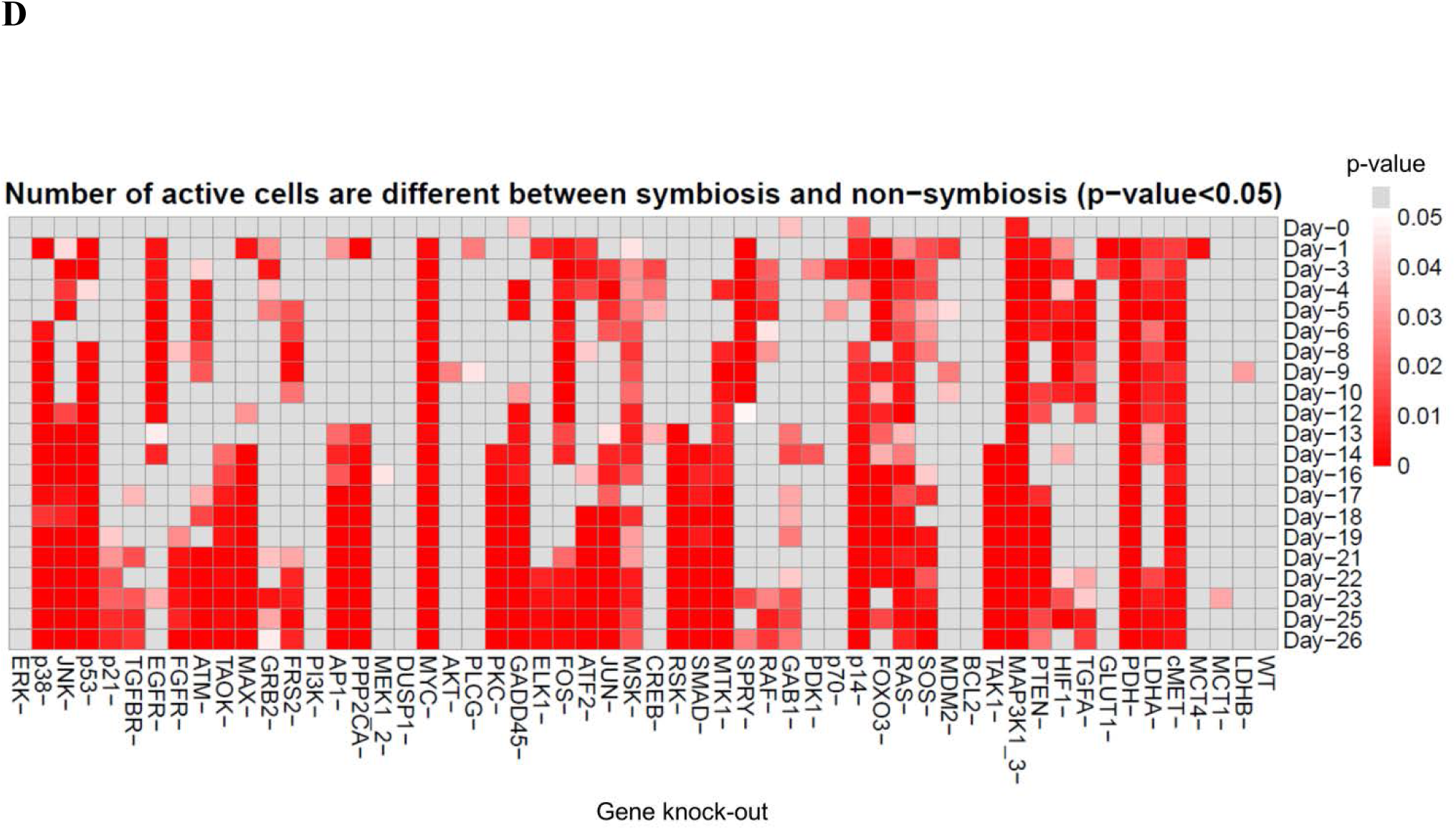
Metabolic symbiosis simulations with network gene alterations: The gene enriched (+) and knockout (−) status were simulated by setting the respective node of the regulatory network to 1 and 0, respectively. The gene wild type (WT) status was simulated without setting the respective node to either 0 or 1. Each gene was altered individually. **(A)**. Percentage tumour growth increase due to symbiosis is shown for each gene enrichment status. **(B)**. Percentage tumour growth increase due to symbiosis is shown for each gene knockout status. **(C)**. Whether tumour growth is significantly different (p-value < 0.05) between symbiosis and non-symbiosis for each gene enrichment status is shown. **(D)**. Whether tumour growth is significantly different (p-value < 0.05) between symbiosis and non-symbiosis for each gene knockout status is shown. Clusters of gene alterations can be identified, which enhance tumour growth due to symbiosis while some other gene alterations together with symbiosis adversely affect tumour growth (A, B). Colours indicate percentage growth increase by symbiosis (A, B) and p values (C, D). p values from 0 to 0.05 are shown in red to white colour scale and p values ≥ 0.05 are shown in grey colour.

Gene enrichments such as EGFR+, ERK+ and DUSP1+ and gene knockouts such as p53-, p38-, PDH- significantly change the number of active cells obtained between symbiotic and non-symbiotic tumours **(Figs 4C-D** and **Fig S7E)**. Similarly, gene enrichments such as MDM2+, ERK+ and DUSP1+ and gene knockouts such as p53-, p38-, p14- significantly increase the number of active cells obtained under symbiosis compared to non-symbiosis conditions **(Figs S7C-E)**.

It is important to note that under MYC+ condition, the tumour growth is increased under both MCT1wt and MCT1-conditions. However, the tumour growth increase in the presence of symbiosis (MCT1wt) is slightly less than the tumour growth increase in the absence of symbiosis (MCT1-), and therefore the percentage change in tumour growth due to symbiosis is negative (i.e., decrease) as seen in **Fig 4A**. Interestingly, for some genes, including for example GRB2, MYC, and LDHB, the tumour growth in the presence of symbiosis is reduced regardless of the nature of gene alteration (enrichment or knockout). This is somewhat counterintuitive; however, it does not mean that tumour growth rate is exactly the same under both types of gene alterations. For example, GRB2-tumour cells grow faster than GRB2+ cells, though both alterations reduce tumour growth at similar proportions under symbiosis.

These results taken together introduce the important concept that the impact of symbiosis on the tumour, whether beneficial or detrimental, will also depend on the type of gene aberrations present in the tumour cells. Thus, this supports the need for more advanced genomics and metabolic combined classifications of patients, which consider not only the gene expression observed at a given time before treatment, but also the predicted interaction of that specific gene expression profile with a given perturbation. In the present model, knockout of MCT1 transporter blocks lactate consumption and hence metabolic symbiosis is disrupted, but these concepts could be extended to other transporters and receptors.

### 3.3. p53 status affects the metabolic symbiosis between oxygenated and hypoxic cells

The most commonly mutated gene in cancer, and also in our samples, is the tumour suppressor p53. TP53 is mutated in 65.5% of the CCLE breast cancer samples and 32.6% of TCGA breast cancer samples (TCGA, PanCancer Atlas) **[43]**. Not only p53 is well-known to be involved in apoptosis and growth arrest of cells **[44]**, but it is also a regulator of the glycolysis pathway with mutations in p53 deregulating cell metabolism in a number of ways including increased GLUT expression **[45, 46]**. Therefore, we studied the effect of p53 alterations on metabolic symbiosis of oxygenated and hypoxic cells in cancer by growing p53wt and p53-cells in isolated populations in our model. **Fig 5** shows variations of cell numbers, cell metabolic pathway utilization and ATP production rates for p53wt and p53 knockout (p53-) simulations. Tumour growth is predicted to be accelerated under p53-, which is expected because cell apoptosis is stopped. However, our simulation proposes an additional mechanism for this because in p53- cells, the glucose transporter GLUT1 expression is increased, and therefore the higher glucose uptake contributes to the higher growth rate. In our results, the number of cells producing mitoATP (i.e mitochondrial ATP production through OXPHOS) is greater than the number of cells producing glycoATP (i.e glycolytic ATP production through glycolysis alone). This is a consequence of lactic acid excretion by the glycolytic cells and its utilization by oxygenated OXPHOS tumour cells (**Figs 5A-B**). By comparing OXPHOS/Glycolysis ratio, it is clear that p53-enhances glycolysis over time, partly because inhibiting p53 causes increased glucose uptake as mentioned above (**Fig 5C**). As expected, the cell ATP production rate is found to be independent from both p53 status and metabolic pathway because it was assumed that cells can adjust their nutrient consumption rates to maintain a similar ATP production rate for all metabolic pathways (**Figs 5D-E**). For example, ATP production rates of glycolysis and OXPHOS are similar while glucose consumption by glycolytic cells is much higher than OXPHOS cells.

**Fig 5.**
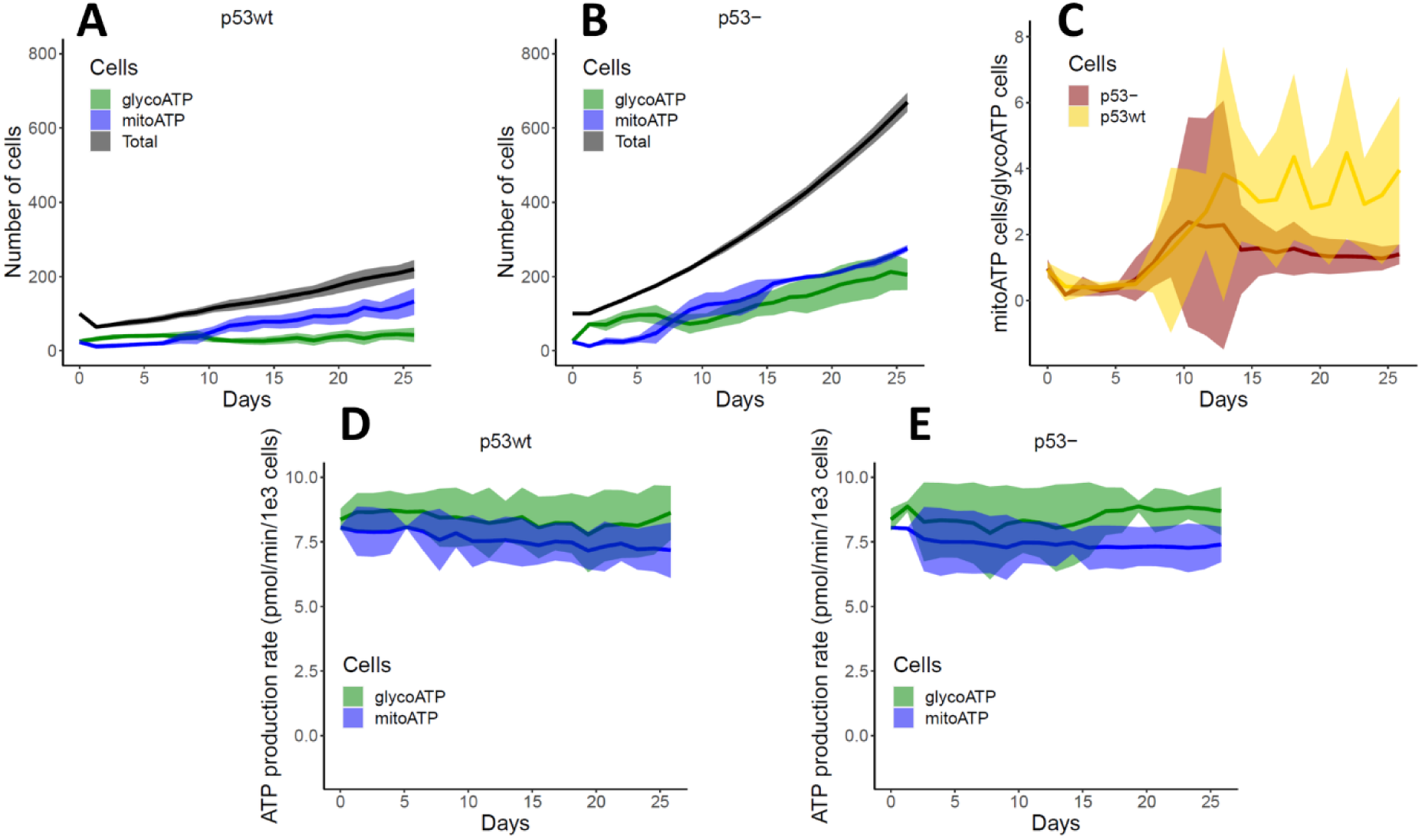
Tumour growth under wild type and mutated p53 status: Effect of p53 status on OXPHOS (shown as mitoATP), glycolysis (shown as glycoATP) and ATP production rate are shown. p53wt and p53-cells were grown in isolation. The starting number of cells was 100. **(A, B)**. Growth of p53wt (A) and p53- (B) cells over time. The OXPHOS cell population is marginally dominant over glycolytic population regardless of the p53 status. **(C)**. Evolution of OXPHOS/Glycolysis cell ratio with p53wt and p53-status. **(D, E)**. Variation of OXPHOS (mitoATP) and glycolytic (mitoATP) ATP production rate of p53wt (D) and p53- (E) cells as tumour grows. The tumour cells with p53 knockout were found to grow faster than p53wt cells because they lacked p53 to suppress tumour growth.

**Figs 6A-D** compare the oxygen, glucose, lactate and pH levels in the medium over time for p53wt and p53-simulations. It shows that oxygen and glucose are depleted at the center of the tumour because when tumour size increases less nutrients would diffuse to the center of the tumour, or they would be consumed at higher rates by the cells at the tumour rim. Lactate accumulates at the center of the tumour making it more acidic. For the p53-tumour, we can see that the tumour microenvironment becomes more acidic compared to the p53wt tumour. **Figs 7A-B** show percentage change of glucose and oxygen in the medium due to metabolic symbiosis for both p53wt and p53-tumours. In the p53wt case, the glucose at the center is increased by symbiosis, while in the p53-simulation, the glucose level at the tumour center is increased by symbiosis only at the initial phase of the tumour growth. As already shown in **Fig 4B**, the tumour gets larger when p53-cells grow with, rather than without, symbiosis, and hence more glucose is consumed by the symbiotic tumour (MCT1wt) than by the non-symbiotic tumour (MCT1-). This would be the reason we see a lower glucose concentration in the center of the tumour at the later stages of symbiotic p53-tumour growth. However, it is vital to note that the glucose level at the tumour boundary where most of the proliferative cells reside is always higher due to symbiosis (**Fig 7A**). With symbiosis occurring between oxygenated and hypoxic tumour cells, oxygenated cells would consume more oxygen for mitochondrial lactate metabolism and therefore symbiosis would result in less oxygen throughout the tumour compared to a non-symbiotic tumour (**Fig 7B**).

**Fig 6.**
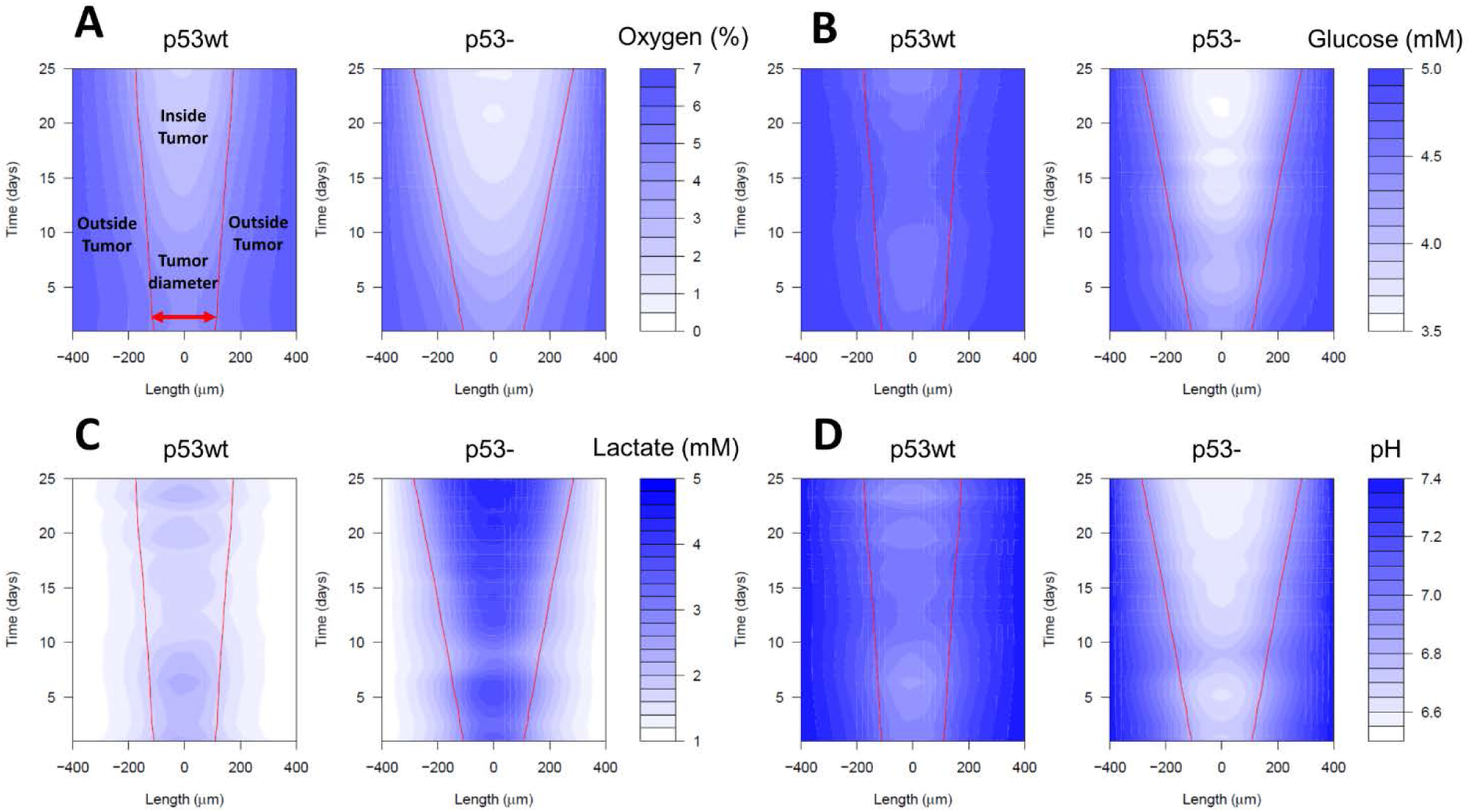
Distribution of tumour microenvironmental substances under wild type and mutated p53 status: **(A-D)**. Heat maps show spatial-temporal variation of microenvironmental (A) oxygen, (B) glucose, (C) lactate, and (D) pH levels for p53wt and p53-tumour growth (here, the Length is the cross section of the tumour microenvironment through the center of the tumour). Red lines show the development of tumour boundary over time. The results show that p53 loss of function could make the tumour microenvironment more acidic during the period of time (i.e., 25 days) considered here. This is mainly because the p53-tumour grows faster and therefore more waste products are accumulated inside the tumour making it more acidic.

**Fig 7.**
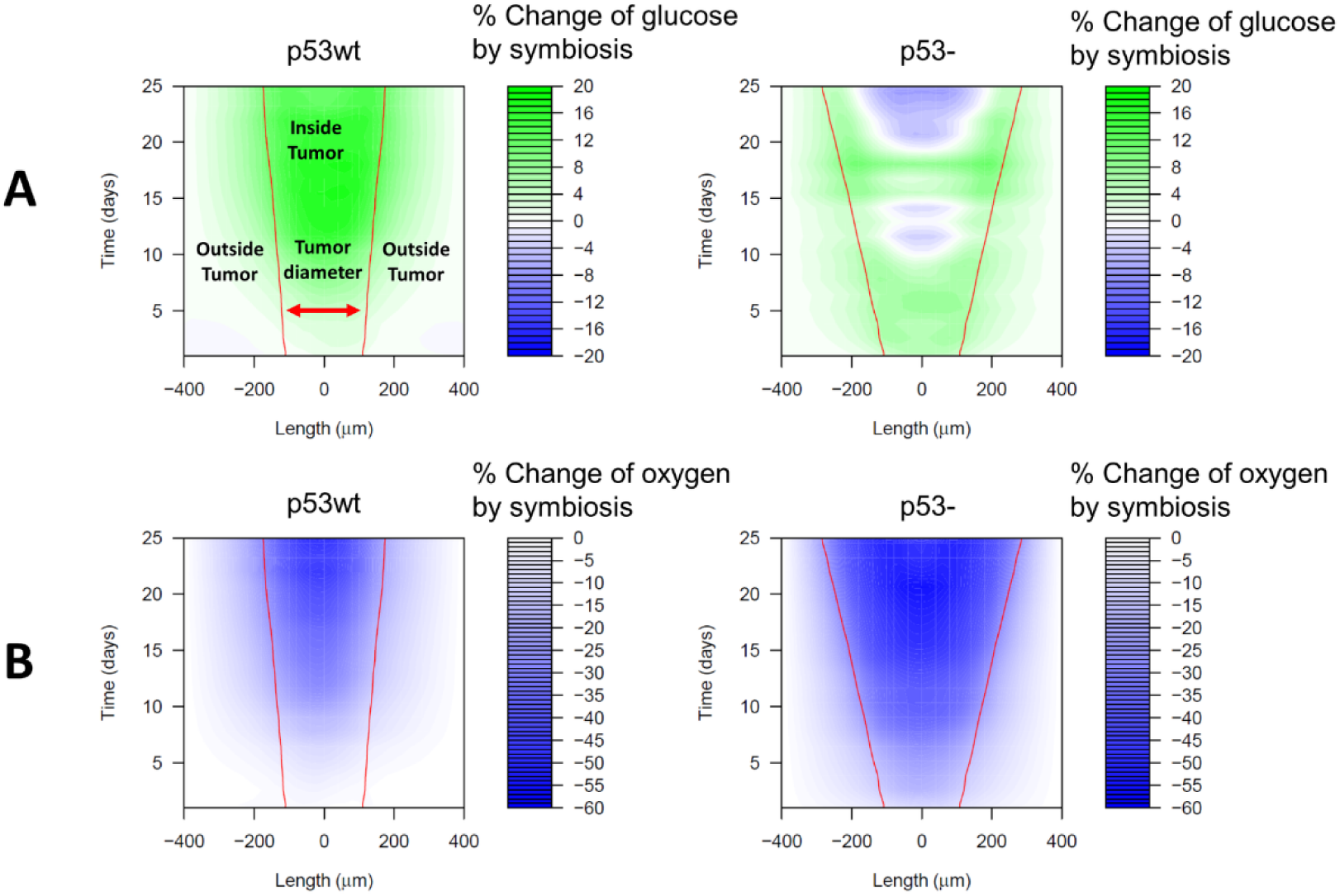
Symbiosis-induced percentage change of glucose and oxygen in the tumour microenvironment: **(A-B)**. Percentage change of (A) glucose and (B) oxygen in the tumour microenvironment over time for tumour growth with p53wt and p53-status. Length and red lines are as described in Fig 6. The model was run with MCT1wt and MCT1-status. The percentage change of glucose and oxygen in the tumour microenvironment was calculated over time. The heat maps suggest that symbiosis increases glucose level while decreasing oxygen level at the tumour boundary. More supplementary results are shown in Fig S8.

To get a clearer insight into how symbiosis affects the levels of extracellular environmental glucose and oxygen, it is necessary to compare symbiotic and non-symbiotic tumours of equal sizes. As tumours grow differently in the two conditions, this would be possible only at the initial stage. Therefore, we performed a sensitivity analysis by varying the initial sizes of the tumour, and compared the nutrient levels between symbiosis and non-symbiosis only at the initial stage of the tumour growth. This allowed us to simulate the contribution of symbiosis on nutrient levels for different sizes of the tumour. **Fig S8A** shows that both p53wt and p53-tumours have an increase in glucose level in the environment due to symbiosis. We observe that the glucose level increment for the largest p53-tumour (i.e., with 500 initial cells) is particularly strong, indicating that the glucose level at the center of the tumour would be increased by about 100% due to symbiosis. **Fig S8B** indicates that metabolic symbiosis would result in a low concentration of oxygen in the tumour microenvironment, and thus a higher degree of hypoxia in the tumour, as already shown in **Fig 3**.

In summary, our simulations illustrate the interaction between p53 status and metabolic symbiosis, and its effect on tumour growth and environmental oxygen and glucose levels. Specifically, symbiosis would increase glucose level (i.e., prevent glucose depletion due to glycolysis) while it would decrease oxygen level at the tumour boundary. These symbiosis-induced nutrient conditions (i.e., increased glucose and hypoxic conditions) of the microenvironment may contribute to further p53-selection or the modulation of other metabolic pathways **[47]**. It should be noted that although microenvironmental glucose level is increased by symbiosis, it is not increased beyond the glucose level set at the boundary of the simulation domain (i.e., 5 mM) in which this behaviour is similar to that of actual tumour glucose levels, which do not increase beyond blood glucose level.

### 3.4. Heterogeneity in the local microenvironment affects the extent of metabolic symbiosis

We asked to what extent changes in the gradient of diffusible substances and cell heterogeneity could impact on the observed metabolic symbiosis. To investigate this, the model was run with different nutrient conditions, with p53 wt/-and MCT1 wt/-, simulating heterogeneous environmental conditions. First, we varied the levels of glucose, oxygen and lactate, for p53wt or p53-cells. When the glucose and lactate levels are kept at a low value (1 mM), the p53wt tumour shrinks over time because dead cells are not replaced by new cells due to lack of nutrients (**Figs S9A-B**). However, in a p53-tumour, cell apoptosis is stopped and those cells can also use the initially provided lactate (5mM) and hence they can proliferate through lactate metabolism when there is enough oxygen. When lactate decreases as it is consumed by tumour cells, the cells are starved and hence OXPHOS cell population starts to decrease after 25 days (**Figs S9C-D**). When we maintain a constant lactate level (5mM) at low glucose level (1 mM), we can see that the tumour continues its growth through lactate metabolism when enough oxygen is available (**Figs S9A, C**). Then, when we maintain low oxygen (3% O_2_) and enough glucose (5 mM) levels at the boundary of the medium, we can see that cells use glycolysis for both p53wt and p53-tumours and the tumour growth continues through pure glycolysis (**Figs S9B, D**). **Fig S10** further shows how the tumour responds to different combinations of oxygen and glucose levels. At low oxygen level (3% O_2_), glycolysis is the dominant pathway and at intermediate oxygen level (6% O2) we can see that the glycolysis and OXPHOS cell populations are comparable. Then, at the high oxygen level (9% O_2_), the dominant cell population is OXPHOS because there is sufficient oxygen for mitochondrial metabolism of either glucose or lactate.

The half-saturation coefficients (*i. e.*, *K_O_*_2_, *K_G_ etc*.) quantify the sensitivity of nutrient uptake by tumour cells to the variation of environmental nutrient levels. **Fig S11** shows that cell’s metabolic pathway would be more sensitive to the half-saturation coefficient of oxygen than that of glucose for the model.

Next, we simulated two p53-cell populations in direct competition; one population had metabolic symbiosis capacity (MCT1wt) and the other population did not (MCT1-).

Initially, two cell populations were randomly mixed at a 1:1 ratio. The glucose and lactate levels were set to either 1 mM or 5 mM. When both glucose and lactate are at the lowest levels, we can see that there is no tumour growth (**Fig 8A**). However, when we increase lactate level to 5 mM at the lowest glucose level, MCT1wt cell population grows while MCT1-cell population is eliminated from the tumour because there is not sufficient glucose for the survival of MCT1-cells (**Fig 8B**). The MCT1wt cell population gets a smaller growth advantage over MCT1-population at 5 mM glucose and 1 mM lactate levels (**Fig 8C**). When both nutrients are at 5 mM, there is no clear difference between two cell populations because there would be enough glucose for MCT1-cells and enough lactate and glucose for MCT1wt cells for their growth (**Fig 8D**).

**Fig 8.**
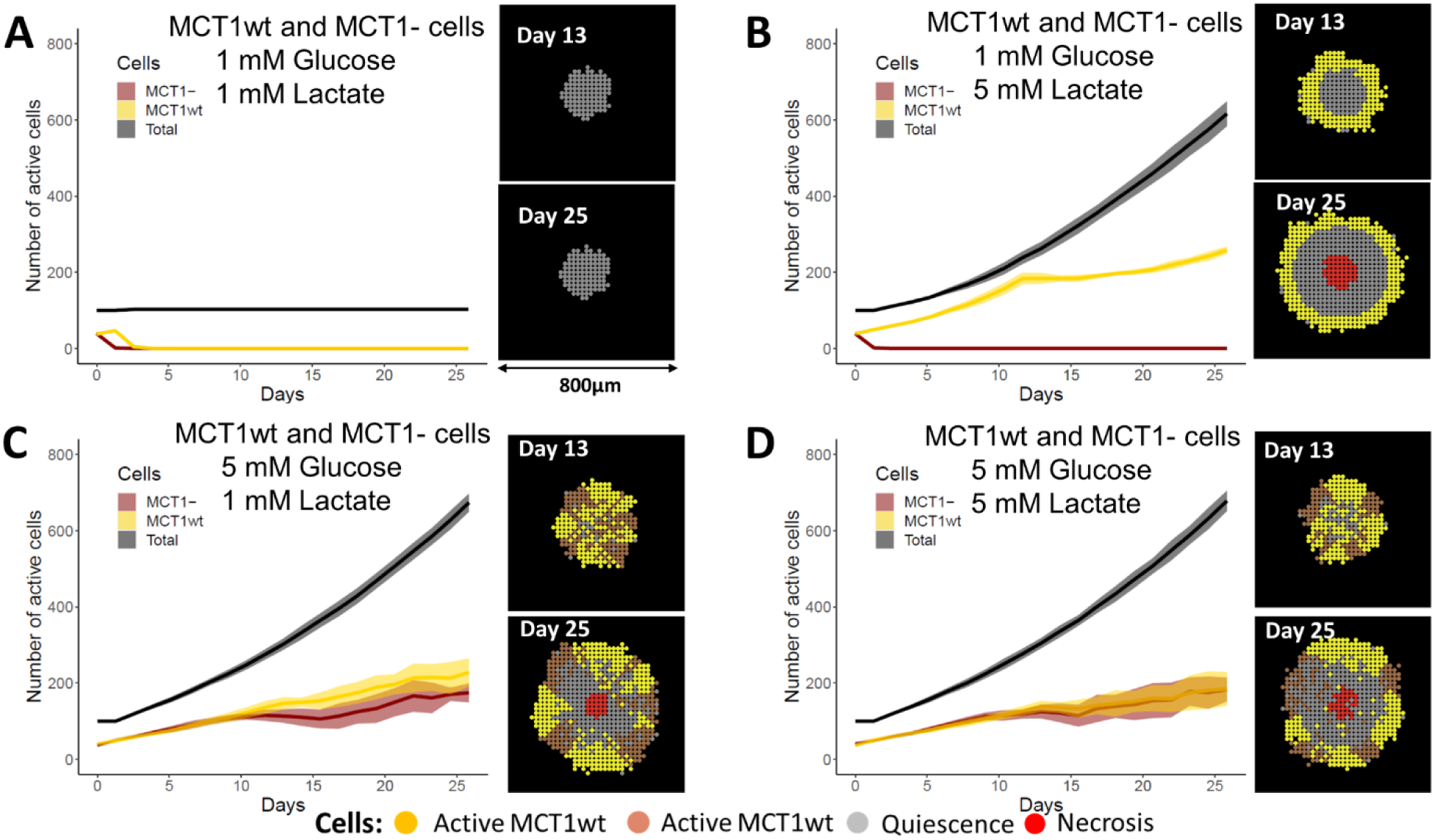
Competition between symbiotic and non-symbiotic tumour cells: Graphs depicting the number of active cells for two p53-cell populations with or without symbiotic capacity, with MCT1wt and MCT1-, respectively. The ratio of MCT1wt: MCT1-was 1:1 at the start of the simulations. Cells were allowed to compete at 6% of oxygen and varying levels of glucose and lactate maintained at the boundary of the computational domain. The number of total, active MCT1wt and active MCT1-cells are shown: **(A)**. 1 mM glucose and 1 mM lactate, **(B)**. 1mM glucose and 5 mM lactate, **(C)**. 5 mM glucose and 1 mM lactate, **(D)**. 5 mM glucose and 5 mM lactate. The results show that symbiotic cells would not get a significant competitive advantage over non-symbiotic cells if there is sufficient glucose in the tumour microenvironment.

The above results show that the nutrient levels in tumour cells not only induce a switch to different metabolic pathways for tumour survival, but also would apply a selection pressure on the spatial competition between symbiotic and non-symbiotic cells.

### 3.5. GLUT and MCT1 as actionable targets to disturb metabolic symbiosis

Inhibitors of metabolic pathways have been investigated as potential cancer therapeutics and various proteins involved in glucose metabolism such as GLUT, HK and LDH have been considered **[48, 49]**. Lactate transporter inhibitors block the Warburg and reverse Warburg effects and hence lactate-driven OXPHOS would be reduced **[50–53]**. However, blocking either glucose or lactate intake transporters would potentially force cancer cells to switch between glucose and lactate metabolic pathways and exhibit therapy resistance. We hypothesized that tumour cells would switch between glucose and lactate metabolic pathways when either pathway was blocked. Therefore, we simulated how the tumour would respond to GLUT1 and MCT1 inhibitors (denoted as GLUT1i and MCT1i) at varying concentration levels and investigated whether tumours would show therapy resistance against respective drugs. Both inhibitors were introduced at the beginning of simulations at a constant level at the boundary of the simulation domain. We assumed that these inhibitors would reach tumour cells by pure diffusion through the tumour microenvironment. It was assumed that the maximum inhibitory level of each drug is 85% and the drug would inhibit the respective membrane protein at a probability of 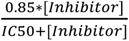, where [*Inhibitor*] and *IC50* are the inhibitor concentration near the cell and half-maximum inhibitory concentration of the inhibitor, respectively.

**Fig 9A** and **Figs S12A-C** show p53wt tumour growth at different levels of MCT1i and GLUT1i. We can see that the OXPHOS cell population dominates in the tumour at low concentration of both drugs. As the MCT1i drug concentration is increased, the glycolytic cell population dominates over OXPHOS. At GLUT1i concentration of 10×IC50, p53wt tumour growth is completely stopped because glycolysis is disturbed and hence apoptotic cells are not replaced by new cells. For p53-cells **(Fig 9B** and **Figs S12D-F)**, we can see that we need a higher concentration of drugs to reduce tumour growth and it shows that p53-tumour may exhibit more resistance against these inhibitors. These results show that p53-tumour cells would exhibit some therapy resistance for GLUT1 inhibitors by using lactate metabolism. However, metabolically active cells (i.e., green and blue coloured cells) would decrease as the concentrations of both drugs are increased. **Guan and Morris (51)** have shown that MCT1 inhibitors AZD3965 and CHC can block lactate uptake and hence tumour growth of 4T1 murine mammary carcinoma cell line. **Guan, Rodriguez-Cruz (50)** have also shown that AR-C155858 and AZD3965 would reversibly inhibit MCT1 of 4T1 cells. As reported in **[54]**, AR-C155858 is effective in reducing mammosphere formation of MCF7 and T47D cell lines. All these findings indicate that MCT1 would be a potential therapeutic target for breast cancer. However, our results suggest that blocking MCT1 would force tumour cells to switch to glycolysis and develop resistance against MCT1 inhibitors regardless of the p53 status. A recent clinical trial of the MCT1 inhibitor AZD3965 showed that some patients had slight increase in ^18^FDG uptake suggesting a lack of anti-tumour effect of the drug **[55]**. Yet, it should be noted that blocking MCT1 can disrupt metabolic symbiosis and therefore it would have unfavourable effect on tumour growth because glucose would be depleted faster when symbiosis is blocked as discussed earlier. Various GLUT inhibitors such as Glutor and Glupin seem to be effective in impairing glucose metabolism of tumour cells **[56, 57]**, but our results suggest that tumour cells may resist GLUT inhibitors if the microenvironment has enough lactate.

**Fig 9.**
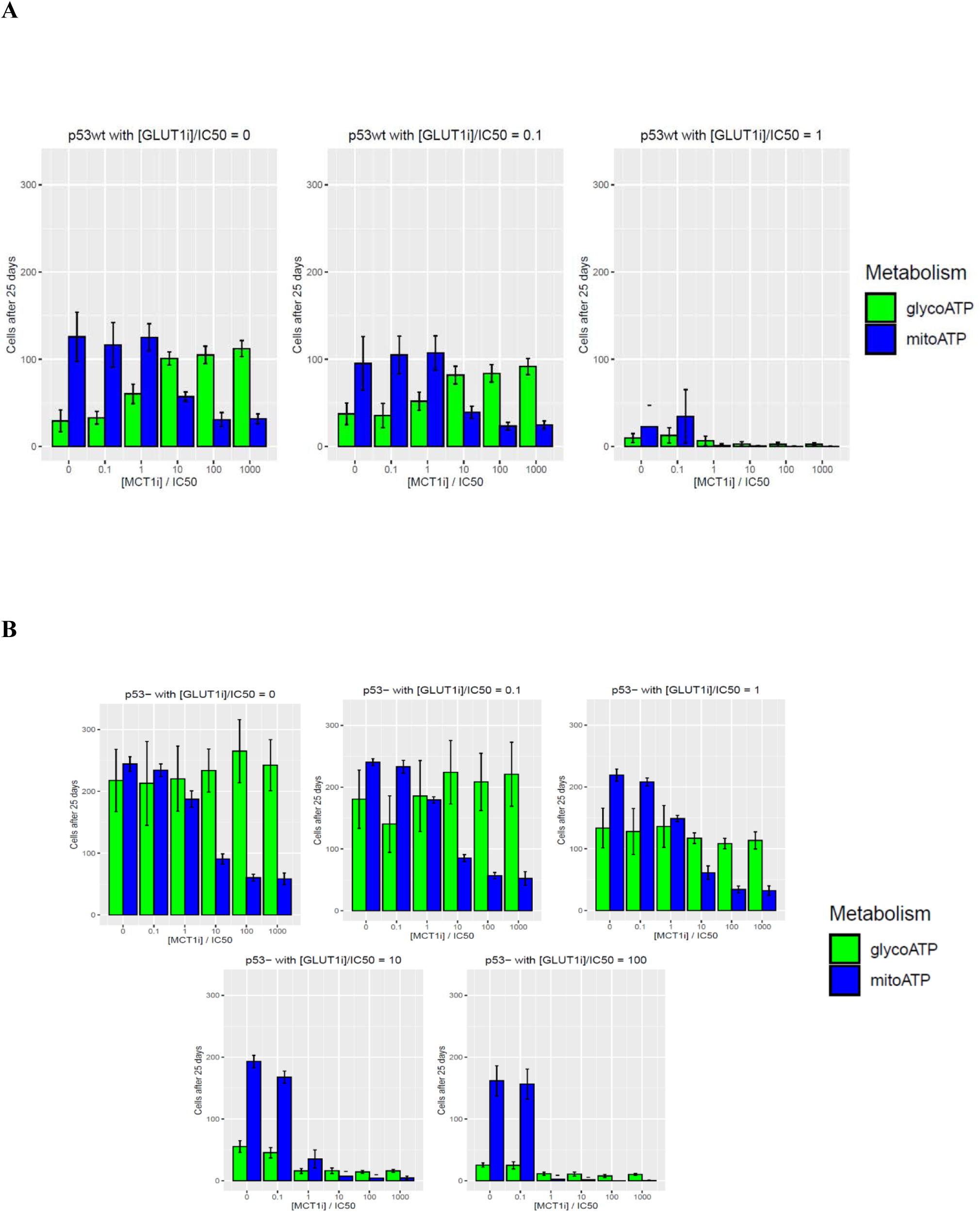
MCT1 and GLUT1 inhibition: Bar charts depicting how MCT1 and GLUT1 inhibitors (denoted as MCT1i and GLUT1i) interfere with glycolysis, OXPHOS, and tumour growth. The MCT1i concentration, [MCT1i], was varied up to 1000 times of its half-maximal inhibitory concentration (IC50) value while GLUT1i concentration, [GLUT1i], was varied up to 100 times of its IC50 value. Inhibitor concentration was maintained at the boundary of the computational domain throughout the simulation time. **(A)**. Number of glycolytic and mitochondrial ATP producing cells of p53wt tumour after 25 days of growth. **(B)**. Number of glycolytic and mitochondrial ATP producing cells of p53-tumour after 25 days of growth. The initial tumour consisted of 100 cells, with either with p53wt or p53-status.

Our results show that even though metabolic symbiosis can be disturbed and hence tumour growth can be reduced by targeting MCT1 and GLUT1 transport proteins in concordance with previous experimental studies, the tumour cells can also exhibit some resistance against those inhibitors by switching between glucose and lactate metabolic pathways. However, the model also predicts that a combination of MCT1 and GLUT1 inhibitors would be synergistic and work against drug resistance and the tumour growth can be completely stopped if both inhibitors are applied at an appropriate concentration. Though tumour growth can be completely stopped at high inhibitor concentrations, it is important to note that inhibitor level and schedule need to be carefully selected in clinical trials with consideration of potential normal tissue toxicity.

### 3.6. Metabolic symbiotic dynamics would depend on the characteristics of cells and microenvironment

As many of the assumptions were taken from existing literature, we performed a sensitivity analysis based on Latin Hypercube Sampling (LHS) and partial rank correlation coefficients **[38, 58]** to identify the parameters that more strongly affected our predictions. The model parameters related to diffusion and activation thresholds of substances (**Table. S3**) were varied within 20% from their baseline values. The total number of parameters was 20. To perform LHS, the range of each parameter (i.e., 80% - 120% of the baseline value) needs to be divided into an equal number of bins. The number of bins was chosen as 40 (for LHS, the number of bins needs to be more than 4×number of parameters/3) and then the LHS input parameter matrix was created according to the LHS sampling method. Then, our model was run for the parameter set in each bin and the model outputs were recorded against each parameter set. The values of each parameter and each output were ranked across bins from 1 to 40 and then the partial correlation coefficient was calculated between each parameter and each output.

**Fig S13** shows the statistically significant (p <0.05) partial correlation coefficients between the model parameters and four model outputs. We can see that the model outputs are correlated with some model parameters, but in a time dependent manner. The results show that the number of total cells, OXPHOS, glycolytic, and necrotic cells are not significantly sensitive to most of the parameters. This could be because the tumour size would mainly be determined by the phenotypes of boundary cells rather than inner cells which have greater contact inhibition due to denser packing. The total number of cells is negatively correlated with the glucose activation threshold (i.e., minimum glucose level needed to activate Glucose_supply node of the network. See S1 Text for more details about activation threshold) and lactate diffusion coefficient. This behaviour is intuitive because as the glucose activation threshold is increased, a fewer number of cells would have glycolysis and when the lactate diffusion coefficient is increased more lactate would diffuse out from the tumour and hence less lactate would be available for OXPHOS. The oxygen diffusion and half-saturation coefficients have positive correlation with OXPHOS at early and later stages of tumour growth, respectively. These two parameters would increase the oxygen level in the medium and hence this dependency could be expected. The oxygen consumption and activation threshold are strongly and negatively correlated with OXPHOS at later stages of tumour growth because increase of these parameters would cause lack of oxygen for respiration.

Lactate diffusion coefficient and activation threshold also have strong negative correlation with OXHPOS at early stage of the growth because increase of these parameters would weaken lactate intake through MCT1 transporters. The glucose activation threshold and lactate diffusion coefficient negatively correlate with glycolysis at later stages of tumour growth. The strong positive correlation between the lactate activation threshold and glycolysis at the early stage of the growth would be due to lactate intake being disturbed by increase in the threshold and hence more tumour cells tend to use glycolysis rather than lactate-fuelled OXPHOS. The oxygen consumption rate and glucose deficient necrosis threshold have some positive correlations with necrosis because both parameters would enhance nutrient depleted conditions in the medium.

**Fig S14** shows how symbiosis-induced tumour growth correlates with model parameters. The glucose activation threshold strongly and positively correlates with growth advantage gained by symbiosis. This is because an increased activation threshold could decrease glucose consumption and glycolysis and it would then force more tumour cells to maintain their growth through lactate metabolism. Therefore, as glucose activation threshold is increased, tumour cells which have symbiotic capacity would gain a growth advantage over non-symbiotic tumour cells.

Our results show that tumour has a symbiosis-induced growth advantage and domination of the mitochondrial ATP producing cell population over glycolytic cell population for the whole LHS parameter space (**Fig S15**). These results suggest that a 20% perturbation of the baseline parameters would not change qualitative behaviour of the model output and therefore the conclusion of our work would remain the same throughout the LHS parameter space. However, if changes were beyond these levels the specific conditions would need to be set, and the simulations reassessed.

## 4. CONCLUSIONS

Agent-based modelling is a powerful simulation technique adopted in many fields of sociology, engineering and biology to study complex emerging behaviours of heterogeneous systems. Here, we used and extended a multi-scale framework merging multi-agent modelling, diffusion-reaction and stoichiometric equations, and gene networks to understand how metabolic symbiosis between oxygenated and hypoxic tumour cells influence tumour growth. We show that:

i. Glucose level in the tumour microenvironment can be maintained by metabolic symbiosis and that this symbiosis helps hypoxic tumour cells survive through glycolysis, using this extracellular glucose. We found that the oxygen level of the medium would be decreased due to symbiosis and then more tumour cells would switch to hypoxic state. The nutrient conditions in the medium would vary over time as tumour evolves. It is well-known that hypoxic tumour cells can exhibit cancer therapy resistance including radiotherapy **[59]** and therefore, this metabolic symbiosis would indirectly increase therapy resistance. For instance, as it is clear in the literature **[60, 61]**, blocking lactate uptake would sensitize tumour xenografts to radiotherapy.
ii. Metabolic symbiosis would allow more glycolytic cells to switch to lactate-driven OXPHOS and hence more glucose would be available for the remaining glycolytic cells. Therefore, this metabolic cooperation would increase the active cell population in the tumour. The dependence on glucose may explain some of relationship of diabetes mellitus to cancer incidence and prognosis.
iii. As glucose and oxygen levels increase in the medium, the glycolytic and OXPHOS cell populations respectively increase. The outcome of the direct competition between symbiotic and non-symbiotic cells of the tumour would depend on the environmental nutrient levels. For instance, symbiotic cells would have a growth advantage in a glucose-limited lactate medium. The pathways in which alteration (i.e., under or over expressions) in the regulatory network would have beneficial or detrimental effects due to symbiosis on the tumour growth were identified.
iv. Some gene alterations (i.e., knockout or enrichment) can interact with metabolic symbiosis to decrease tumour growth. Furthermore, both p53wt and p53-cells show some resistance against MCT1 inhibitors. However, use of MCT1 inhibitors in combination with GLUT1 inhibitors may disrupt metabolic cooperation between oxygenated and hypoxic cell populations.

Our sensitivity analysis shows that the model outputs are sensitive to some model parameters only at different stages of tumour growth. However, the results show that the qualitative behaviour of the model would remain the same in whole parameter space which is within ±20% from the baseline value of each parameter.

The present model is a relatively simplified model of the metabolic symbiosis between hypoxic and oxygenated tumour cells. The model could be used to understand how metabolic changes induced by therapy in different genetic backgrounds interact. Future work should consider using robust cell regulatory networks including pathways such as lipid metabolism signalling pathway, adding further microenvironmental features (fibroblast, ECM, immune cells etc.), tumour heterogeneity and therapy modelling. Ultimately, any simulation result would need to be confirmed in validation in pre-clinical setting but the findings from this study could help in guiding the experimental design.

## SUPPORTING INFORMATION

**S1 Text. Methodology, Parameters and Supplementary Results:** Different spatial and temporal scales of the model and how these scales interact each other are described. More details about the cell regulatory network and model parameters are given. Supplementary results to those results presented in the manuscript are also included.

**S1 Chart. Breast Cancer Cell Lines Gene Expressions:** Breast cancer cell line gene expression data under normoxic and hypoxic conditions are given. The unit is log2 (FPKM+2). The data were used to produce Fig S4.

**S1 File. Cell Regulatory Network:** The source file to open the network with GINsim software (http://ginsim.org) and all the Boolean logical conditions and respective supporting evidence can be seen there.

**S2 File. Cell Regulatory Network:** A high resolution image of the network.

## Supporting information

S1-Text-Methodology_Parameters_and_Results

S1-Chart-Breast_Cell_Lines_Gene_Expressions

S1-File-Cell_Regulatory_Network

S2-File-Cell_Regulatory_Network

## ACKNOWLEDGEMENT

This work was funded by the European Research Council (ERC) under the European Union’s Horizon 2020 research and innovation programme (Project name microC, Grant agreement No. 772970). Harris AL was funded by Breast Cancer Research Foundation. The funders had no role in study design, data collection and analysis, decision to publish, or preparation of the manuscript. The computer simulations were run on High-Performance Computing (HPC) clusters at the Advanced Research Computing (ARC) facilities of the University of Oxford. All members and collaborators of the Buffa Lab of the University of Oxford are appreciated for their valuable comments and suggestions. We would also like to express our gratitude to Dr. Kenneth Kahn for his invaluable assistance in setting up the HPC system.

## AUTHOR CONTRIBUTIONS

**Conceptualization:** Buffa FM, Harris AL, Morten KJ

**Data Curation:** Jayathilake PG, Victori P, Pavillet CE, Voukantsis D, Miar A, Arora A, Buffa FM

**Formal Analysis:** Jayathilake PG, Miar A, Buffa FM

**Funding Acquisition:** Buffa FM, Harris AL

**Investigation:** Jayathilake, Miar A, Arora A

**Methodology:** Jayathilake PG, Victori P, Pavillet CE, Voukantsis D, Miar A, Buffa FM

**Project Administration:** Buffa FM

**Resources:** Voukantsis D, Miar A, Buffa FM

**Software:** Jayathilake PG, Victori P, Pavillet CE, Voukantsis D, Buffa FM

**Supervision:** Buffa FM, Harris AL

**Validation:** Jayathilake PG, Victori P

**Visualization:** Jayathilake PG

**Writing-Original Draft Preparation:** Jayathilake PG, Buffa FM

**Writing-Review & Editing:** Jayathilake PG, Victori P, Pavillet CE, Voukantsis D, Miar A, Arora A, Harris AL, Morten KJ, Buffa FM

