## Supplementary material for "Metabolic Symbiosis between Oxygenated and Hypoxic Tumour Cells: An Agent-based Modelling Study": S1-Text-Methodology_Parameters_and_Results

**S1 Text. Methodology, Parameters and Supplementary Results**

1. **Introduction**

The purpose of this supporting information is to provide more details about our multiscale agent-based mathematical model of tumour growth and supplementary results.

**2. Overview of multi-scale agent-based mathematical model**

Our model has three different spatial scales to describe different biological processes and cell-cell and cell-microenvironment interactions. The three scales are the sub-cellular, cellular and extra-cellular scales and these scales are inter-connected each other as shown in **Fig S1**. The sub-cellular scale describes cell-regulatory network and it decide cell phenotype based on signals it receives from the tumour microenvironment and sub-cellular molecular interactions. The cellular scale describes cell-cell and cell-micro environmental interactions and the extra-cellular scale describes spatial distribution of diffusible substances in the tumour microenvironment. Both sub-cellular and cellular scales are modelled as agent-based models while the extra-cellular scale is modelled as a continuum model. More details about each scale are given below.

2.1. **Model framework**
     2.1.1. **Sub-cellular scale: Boolean network**

The cell regulatory network is modelled at the sub-cellular scale by using an agent-based model. This is a Boolean network which contains a MAPK (Mitogen-Activated Protein Kinases) network together with glucose and lactate metabolic pathways as shown in **Fig S2**. This is a mathematical representation of signalling and metabolic pathways involved in cellular responses to stimuli such as growth factors (e.g., Epidermal growth factor, Hepatocyte growth factor) and nutrients (e.g., Glucose, Lactate). The each node represents molecules (e.g., Proteins, Genes), cellular processes (e.g., Electron Transport Chain, Tricarboxylic Acid Cycle) and cell phenotypes (e.g. Proliferation, Apoptosis). In Fig S2, the green and red lines describe positive and negative interactions between nodes, respectively. Each node has its own specific Boolean logical condition and if the condition is true the node is considered as active (i.e., 1) and otherwise inactive (i.e., 0). The vast majority of the network was taken from literature and logical conditions associated with those nodes can be found in [1, 2]. The newly added nodes to the network and associated logical conditions are explained in **Table S1**. To read more about the network, our Boolean network contained in the source file, **S1 File,** can be opened in GINsim software (<http://ginsim.org>) and all the Boolean logical conditions and respective supporting evidences can be seen there. We have converted this network source file to an html file using Oxygen XML Editor to use with our multi-scale model.

This network is encapsulated inside each tumour cell and the network can obtain stimuli such as growth factors and nutrients from the extra-cellular environment through input nodes (i.e., rose coloured nodes of Fig S2). An environmental stimulus can activate the respective input node if local stimulus concentration is above a certain threshold. For example, the input node, Glucose_supply, is activated if the local glucose level is above a certain threshold. Similarly, the input node, Oxygen_supply, is activated if the local oxygen level is above a certain threshold and so on (**Tables S1** and **S2**). The input nodes could be cell membrane receptors such as Epidermal Growth Factor Receptor – EGFR, membrane transport proteins such as Glucose Transporter – GLUT1 and stresses such as DNA damage. Upon receiving stimuli, the network can decide which metabolic pathway to use and that could be glycolysis, glucose-driven oxidative phosphorylation (OXPHOS) or lactate-driven OXPHOS. The network can also decide the cellular phenotype (i.e., blue coloured node of Fig S2) which could be proliferation, growth arrest, apoptosis or necrosis.

2.1.2. **Cellular scale: Cellular automaton method**

At cellular scale, individual cell behaviour is modelled using the Cellular Automaton (CA) method which is a lattice-based agent-based modelling approach. Each cell is represented by a spherical particle and the particle can move on a Cartesian grid according to CA rules [2]. This model describes cell-cell and cell-microenvironmental interactions.

A proliferate cell which has completed the cell division time can only divide if there is an empty space around the cell, otherwise it may remain in contact inhibition or quiescence state for a certain period of time before checking environmental conditions again to find its new phenotype. A cell turns to apoptosis would be removed from the simulation immediately. Growth arrest cells would consume nutrients at half of the normal rate. The necrotic cells would not consume or produce any substance other than exist in in the tumour. Cells can interact with the tumour microenvironment though consumption and production of diffusible substances such as nutrients and growth factors.

2.1.3. **Extra-cellular scale: Tumour microenvironment**

At this scale, following diffusion reaction equation describes the distribution of diffusible substances in the tumour microenvironment.

$\frac{\partial C_{S}}{\partial t}=D_{S}\nabla^{2}C_{S}+R_{S}\rho_{cell,S}$	S1

where *C*, *D* and *R* are the substance concentration, its diffusion coefficient and its rate of consumption or production, respectively. $\rho_{cell,S}$represents the density of cells which consume or produce substance *S*. For example, the growth factor TGFA is produced by only hypoxic tumour cells as modelled in the cell regulatory network and therefore only the cells with active TGFA are included when calculating respective cell density. The diffusible substances include oxygen, glucose, lactate and growth factors. The growth factors are TGFA, HGF and FGF for activation of EGFR, cMET and FGFR, respectively. More details about the substance production and consumption rates, $R_{S}$ , have been discussed in the main text.

The diffusion-reaction equation is solved on a Cartesian grid by using Forward Time Centered Space (FTCS) finite difference method.

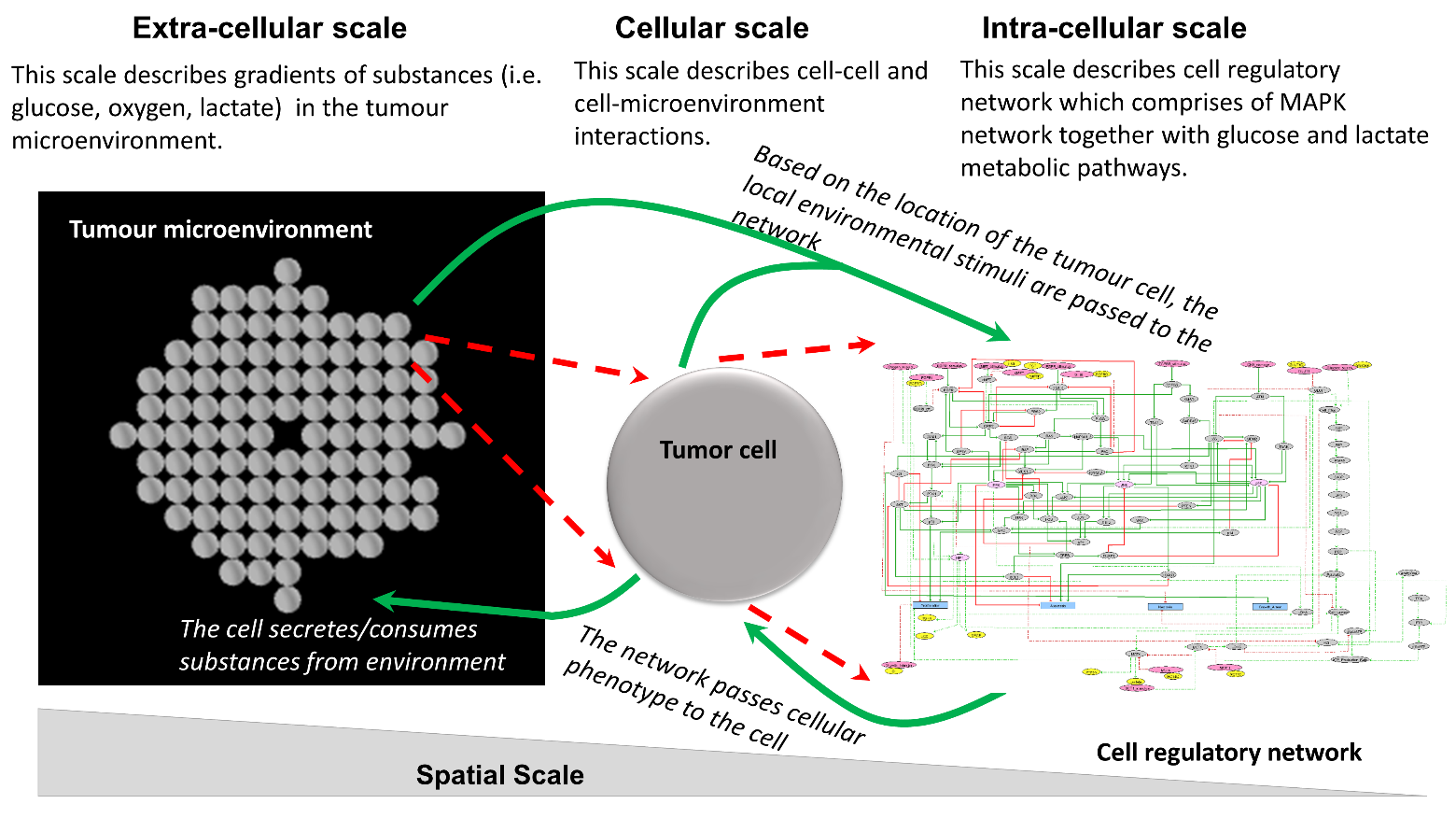

**Fig S1**. Multi-scale modelling framework. The extra-cellular scale, an equation-based model, describes distributions of diffusible substances in the tumour microenvironment. The cellular scale, an agent-based model which is a cellular automaton model, describes cell-cell and cell-microenvironmental interactions. The sub-cellular scale, an agent-based model which is a Boolean network, describes subcellular molecular interactions. The three scales are coupled each other and information are shared between them.

2.2. **Coupling between scales**

As shown in **Fig S1**, three different spatial scales of this tumour model are described by three sub-models using an equation-based (i.e., diffusion-reaction equation) model and two agent-based models (i.e., Cellular Automaton model and Boolean network). Time scales of different processes have also been separated as shown in **Fig S3**. At each *T_Network_*, one randomly selected node of the regulatory network will be updated based on existing Boolean conditions of neighbouring nodes (i.e., 0 or 1) and its own logical condition. The cell phenotype is updated at an interval of *T_Phenotype_* (red ticks) based on the outputs from the regulatory network for given stimuli. The distribution of diffusible substances in the tumour microenvironment are updated at an interval of *T_Diffusion_* (green ticks) by solving the diffusion-reaction equation for steady state at each green tick. A tumour cell would not divide until it completes its cell division time of *T_Division_*.

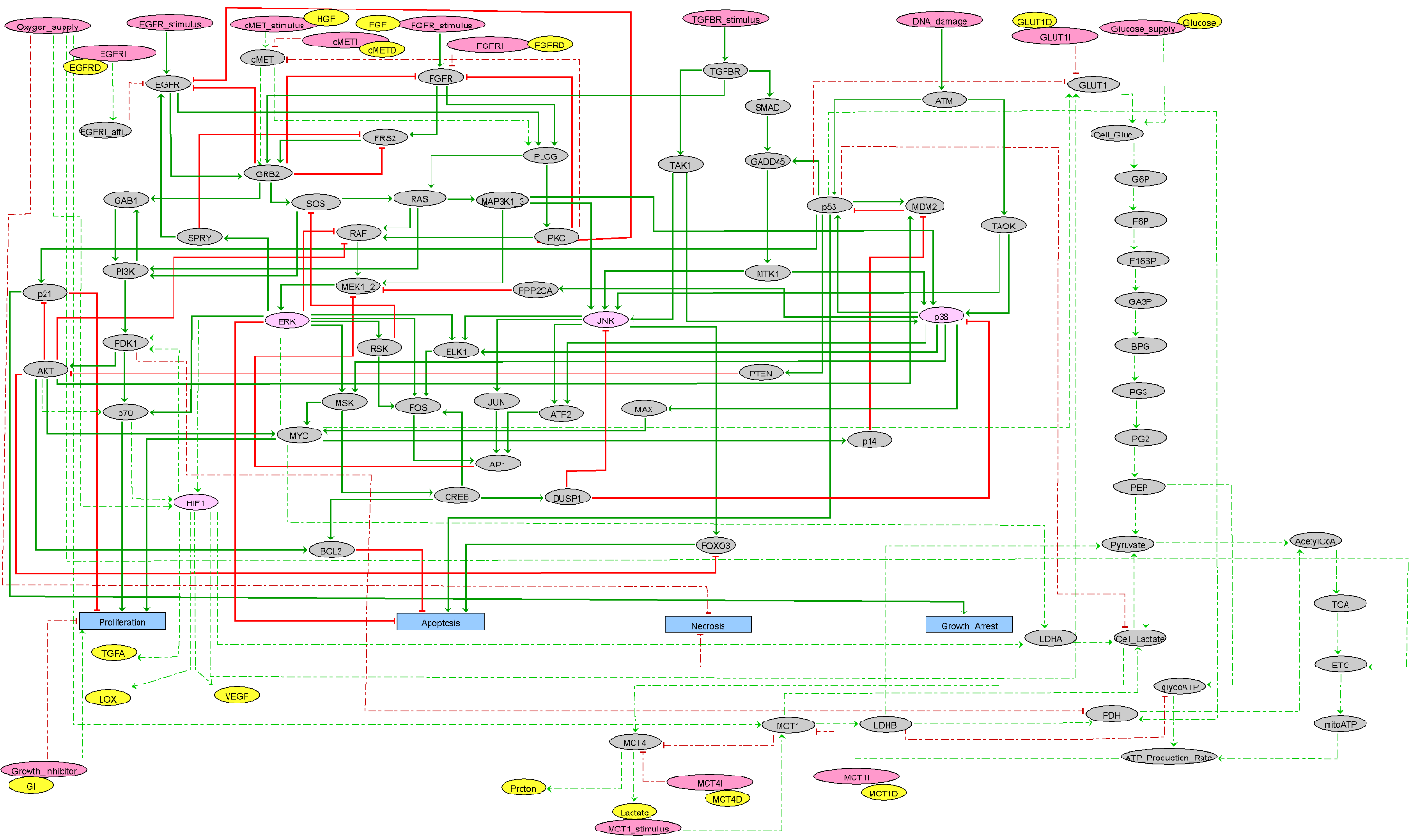

**Fig S2**. The cell regulatory network of our model (microC model). The modified MAPK network decides the cell phenotype based on inputs obtained from the microenvironment. Oxygen_supply, Glucose_supply, EGFR_stimulus, cMET_stimulus, FGFR_stimulus, TGFBR_stimulus, DNA_damage and Growth_inhibitor are inputs to the network. The inhibitor nodes (EGFRI, cMETI, FGFRI, GLUT1I, MCT1I, and MCT4I) are also inputs. The inhibitor nodes are activated by respective drugs (e.g. GLUT1I is activated by GLUT1D, MCT1I is activated by MCT1D and so on). The Boolean network is updated asynchronously and cell phenotypical outputs are calculated. The outputs are Proliferation, Apoptosis, Necrosis and Growth_Arrest. The yellow colored nodes are diffusible substances. The newly added interactions to the original MAPK network taken from the literature [2] are shown in dotted lines. The solid and dotted green lines are positive interactions and solid and dotted red lines are negative interactions. More details about Boolean logical conditions at each node are given in [1, 2] and Tables S1-2. A high resolution image of this network is available in S1 and S2 Files.
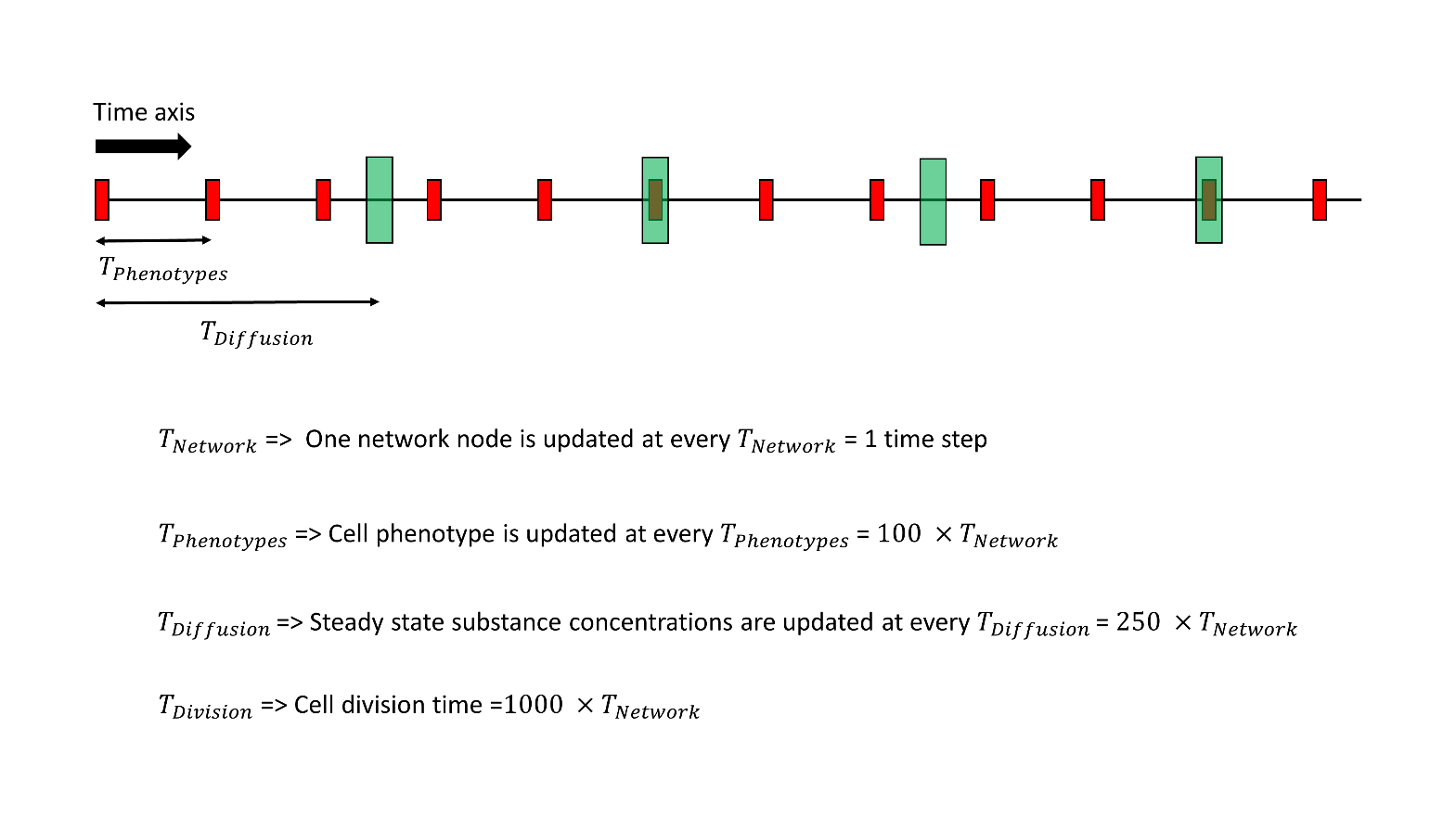

**Fig S3**. Different time scales are used for different proceses. The smallest time step is defined as the time for updating one node of the regulatory network (*T_Network_*). Cell phenotype and diffusion fields are updated at red (*T_Phynotypes_*) and green (*T_Difusion_*) ticks, respectively. A cell which is older than the cell division time (*T_Division_*) can divide if its phenotype is Proliferation.

2.3. **Stochasticity of the model**

Stochasticity appears in the model in following processes.

i. Cell regulatory network update: The regulatory network is initialized randomly, which means each node is initialized with 0 or 1 randomly. Then at each time step, a randomly selected node of the regulatory network will be updated based on its Boolean logical condition.
ii. Initial cell distribution: Different types of cancer cells such as different mutants as set by the user are placed in the tumour microenvironment randomly to make the initial tumour spheroid.
iii. Cell division: When a cell is divided, the new cell is located next to the mother cell in a randomly selected direction.
     2.4. **Software and programming language**

The model has been originally implemented on Netlogo as reported in [2]. The present symbiosis model was implemented on the same modelling platform. The Netlogo model and its dependent files used for this study have been stored in GitHub at <https://github.com/CBigOxf/jayathilake2022> . The Netlogo model file can be run with dependent files given in our repository. More instructions about how to choose parameters are given at the top of the NetLogo code. The model output data were analysed and visualized using R.

3. **Model parameters, experimental methods, and clinical data**
   3.1. **Model parameters**

The model parameters are shown in **Table S3**.

3.2. **Normoxia/hypoxia experiments for breast cancer cell lines**

Hypoxic and normoxic environments were created at 2% and 20% of oxygen, respectively (**Fig S4**).

3.3. ***cBioPortal***

We checked the mRNA expression levels of genes involved in our network model using the breast cancer samples of TCGA PanCancer Atlas (The Cancer Genome Atlas, <https://www.cancer.gov/tcga/>) using the cBioPortal online tool (https://www.cbioportal.org). The mRNA expression levels are given with relative to normal samples (**Fig S5**).

3.4. ***sigQC***

The R package sigQC allows us to check multiple gene signature sets across multiple gene expression datasets ([https://CRAN.R-project.org/package=sigQC](https://cran.r-project.org/package=sigQC)). We checked our signature genes in TCGA and CCLE breast cancer (Cancer Cell Line Encyclopedia, <https://sites.broadinstitute.org/ccle/>) datasets (**Fig S6**).

**Table S1.** Metabolic molecules and regulatory nodes implemented in regulatory network. In the Boolean expressions, &, |, and ! represent AND, OR, and NOT logical conditions, respectively. Only the newly added logical conditions are given here, and the rest of the conditions have been reported in [1]. **S1 File** can be opened in GINsim software (<http://ginsim.org>) and all the Boolean logical conditions and respective supporting evidences can be seen there.

| **Molecule (node)** | **Logical condition** | **Reference** |
| --- | --- | --- |
| GLUT1: Glucose Transporter 1 | (HIF1 \| !p53 \| MYC) & !GLUT1I | [3-7] |
| G6P: Glucose 6-Phosphate | Cell_Glucose & Glucose_supply | - |
| F6P: Fructose 6-Phosphate | G6P | [8] |
| F16BP: Fructose 1,6-Bisphosphate | F6P | [8] |
| GA3P: Glyceraldehyde 3-Phosphate | F16BP | [8] |
| BPG: 1,3-Bisphosphoglycerate | GA3P | [8] |
| PG3: 3-Phosphoglycerate | BPG | [8] |
| PG2: 2-Phosphoglycerate | PG3 | [8] |
| PEP: Phosphoenolpyruvate | PG2 | [8] |
| AcetylCoA: Acetyl Coenzyme A | Pyruvate & PDH | [8] |
| TCA: Tricarboxylic Acid Cycle | AcetylCoA | [9] |
| ETC: Electron Transport Chain | TCA & Oxygen_supply | [9] |
| mitoATP: mitochondrial ATP | ETC | [9] |
| LDHA: Lactate Dehydrogenase A | HIF1 & MYC | [6, 7] |
| LDHB: Lactate Dehydrogenase B | MCT1 | [10] |
| PDH: Pyruvate Dehydrogenas | !PDK1 \| p53 \| LDHB | [11-14] |
| glycoATP: glycolytic ATP | PEP & !LDHB | [10] |
| MCT1: Monocarboxylate transporter 1 | Oxygen_supply & MCT1_stimulus & !MCT1I | [10] |
| MCT4: Monocarboxylate transporter 4 | Cell_Lactate & !MCT1 | [10] |
| GLUT1I: GLUT1 Inhibitor | GLUT1 Drug > threshold | - |
| MCT1I: MCT1 Inhibitor | MCT1 Drug > threshold | - |
| cMET | cMET_stimulus & !cMETI | - |
| GRB2 | EGFR \| cMET \| FRS2 \| TGFBR | KEGG networks |
| PDK | PI3K \| HIF1 \| MYC | [6, 7] |

**Table S2.** Input nodes and corresponding stimulus of the regulatory network. If the stimulus concentration around a tumour cell is beyond a certain threshold the respective input node is activated for that particular cell. These threshold values are given in Table S3.

| **Input node** | **Stimulus** | **Used in the present study** |
| --- | --- | --- |
| Oxygen_supply | Oxygen | Yes |
| Glucose_supply | Glucose | Yes |
| EGFR_stimulus | TGFA | Yes |
| cMET_stimulus | HGF | Yes |
| FGFR_stimulus | FGF | No |
| TGFBR_stimulus | TGFB | No |
| MCT1_stimulus | Lactate | Yes |
| DNA_damage | Factors causing DNA damage | No |
| EGFRI | EGFRD (EGFR Drug) | No |
| cMETI | cMETD (cMET Drug) | No |
| FGFRI | FGFRD (FGFR Drug) | No |
| GLUT1I | GLUT1D (GLUT1 Drug) | Yes |
| MCT1I | MCT1D (MCT1 Drug) | Yes |
| MCT4I | MCT4D (MCT4 Drug) | No |
| Growth_inhibitor | GI (Growth Inhibiting factors) | No |

**Table S3.** Model parameters. If there is no reference for a parameter, the value is chosen such that the model work smoothly and the value is also close to its typical order of magnitude.

| **Parameter** | | **Value** | **Reference** | **Remarks** |
| --- | --- | --- | --- | --- |
| Oxygen | Diffusion coefficient | 1.0 × 10^-9^ m^2^/s | [15] |  |
|  | Consumption rate | 3.00 × 10^-17^ mol/s/cell | [16] |  |
|  | Half-saturation coefficient | 0.005 mM | [17] |  |
|  | Boundary level | 0.07 mM | [18] |  |
|  | Necrosis level | 0.011 mM (1% P_O2_) | [19] | Pathological hypoxia |
|  | Activation threshold | 0.022 mM (2% P_O2_) | [19] | Physiological hypoxia |
| Glucose | Diffusion coefficient | 6.7 × 10^-11^ m^2^/s | [20] |  |
|  | Half-saturation coefficient | 0.04 mM | [17] |  |
|  | Boundary level | 5.0 mM | [18] |  |
|  | Necrosis level | 3.9 mM | Assumed | A wide range of values seen in the literature. 0.06 mM [21], 0.15 µM [22]. 2.27 mM [23], 8mM [20]. |
|  | Activation threshold | 4.0 mM | [24] | [24] has used 5 mM, but we reduce it to 4 mM such that we see tumour growth at 5mM which is commonly seen in literature. |
| Lactate | Diffusion coefficient | 6.7 × 10^-11^ m^2^/s | [21] |  |
|  | Half-saturation coefficient | 0.04 mM | Assumed | Assuming same as for glucose |
|  | Boundary level | 1 mM | Assumed | Low lactate level far from tumour |
|  | Activation threshold | 1.5 mM | [25] |  |
| TGFA  (*Growth factor of EGFR*) | Diffusion coefficient | 5.18 × 10^-11^ m^2^/s | [20, 26] |  |
|  | Consumption rate | 2.00 × 10^-17^ m^3^/s/cell | Assumed | Chosen such that reasonable TGFa amount is maintained in the medium. |
|  | Production rate | 2.00 × 10^-20^ mol/s/cell | Assumed | Chosen such that reasonable TGFa amount is produced which is in nM scale. 4.00 × 10^-25^ [20] |
|  | Activation threshold | 1.0 nM | Assumed | Typical TGFa concentration of tissues is in the order of nM [27] |
|  | Boundary level | 0.0 | [2] |  |
| HGF  (*Growth factor of*  *cMET*) | Diffusion coefficient | 8.50 × 10^-11^ m^2^/s | [28] |  |
|  | Consumption rate | 2.00 × 10^-18^ m^3^/s/cell | Assumed | Chosen such that we can see HGF-induced cMET activation in tumour. |
|  | Production rate | 0.0 |  | HGF is externally provided. |
|  | Activation threshold | 1 nM | Assumed | Typical HGF concentration is in order of nM [29] |
|  | Boundary level | 2 nM | Assumed | Chosen such that we can see HGF-induced cMET activation in tumour. |
| H+ | Diffusion coefficient | 1.0 × 10^-9^ m^2^/s | [17] |  |
|  | Boundary level | 40 nM | Assumed | Acidity decreases as moving away from tumour. |
| MCT1i  GLUT1i  (*MCT1 and GLUT1 inhibitors*) | Diffusion coefficient | 2.2 × 10^-10^ m^2^/s | Assumed | Assuming similar to 2.5 × 10^-10^ of Gefitinib/EMD [28] |
|  | Consumption rate | 4.00 × 10^-17^ m^3^/s/cell | Assumed | Assuming similar to TGFA |
|  | Activation threshold | IC50 (17 nM for MCT1i and 4nM for GLUT1i) |  | Typical values [30]  MCT1i=AZD3965 drug  GLUT1i=Glupin drug |
|  | Boundary level | Variable (See the text) |  |  |
| Spatial & temporal parameters | Computational domain | 0.8 × 0.8 mm^2^ | - | A reasonable size for computation |
|  | Cell diameter | 20 µm | [31] |  |
|  | Cell division time | 1000 steps = 1.3 days | [32] |  |
|  | Diffusion time | 250 steps | - |  |
|  | Cell phenotype time | 100 steps | [2] |  |

4. **Supplementary results**

**Figs S4-S6** and **Figs S7-S15** show our experimental/TCGA/CCLE data and simulation data, respectively.

**A**

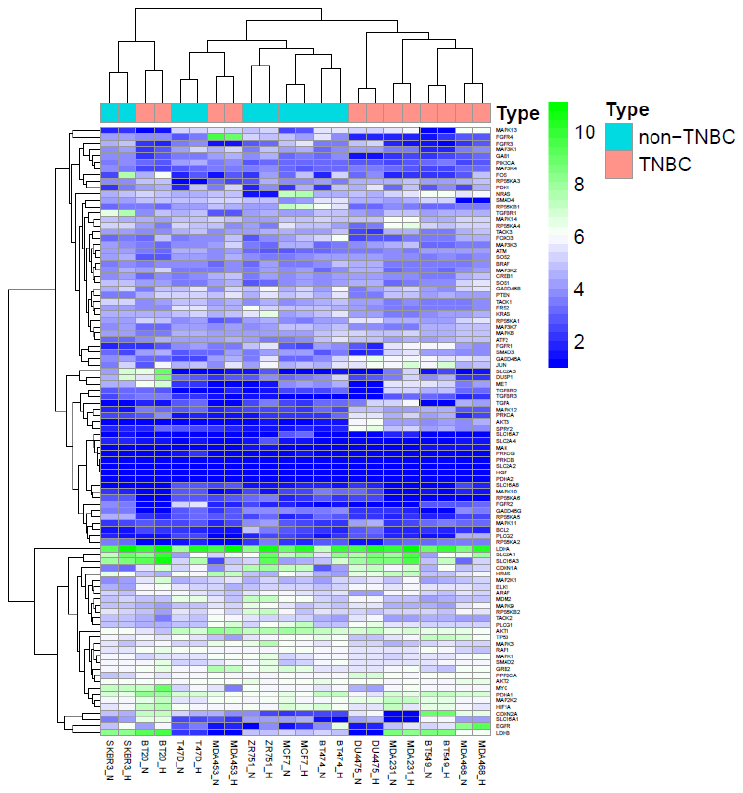

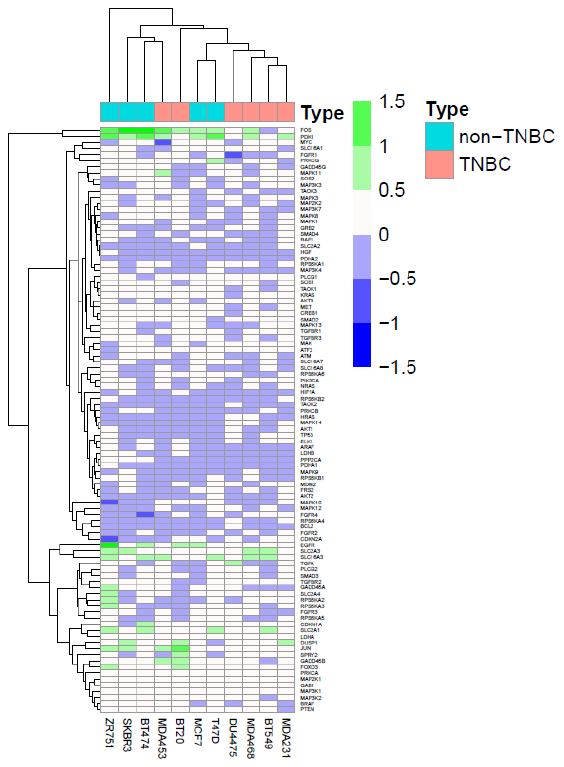

**B**

**Fig S4**. **(A)**. Expression of our network genes of some Triple Negative Breast Cancer (TNBC) and non-TNBC cell lines under normoxia (N) and hypoxia (H). The unit is log2 (FPKM+2). **(B)**. The gene expressions (EXP) at hypoxia is normalized by respective expressions at normoxia (log2 (EXP at hypoxia/ EXP at normoxia)). The data used to produce these Figs are given in S1 Chart.

**A**
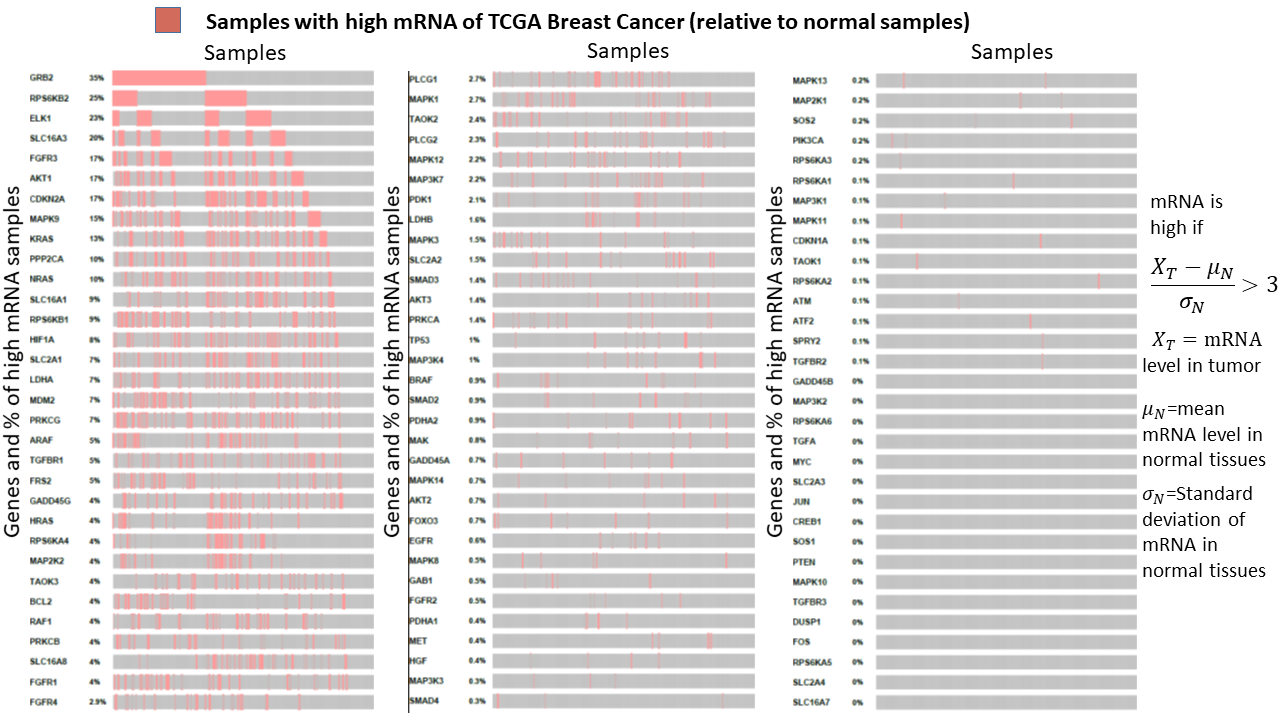

**B**
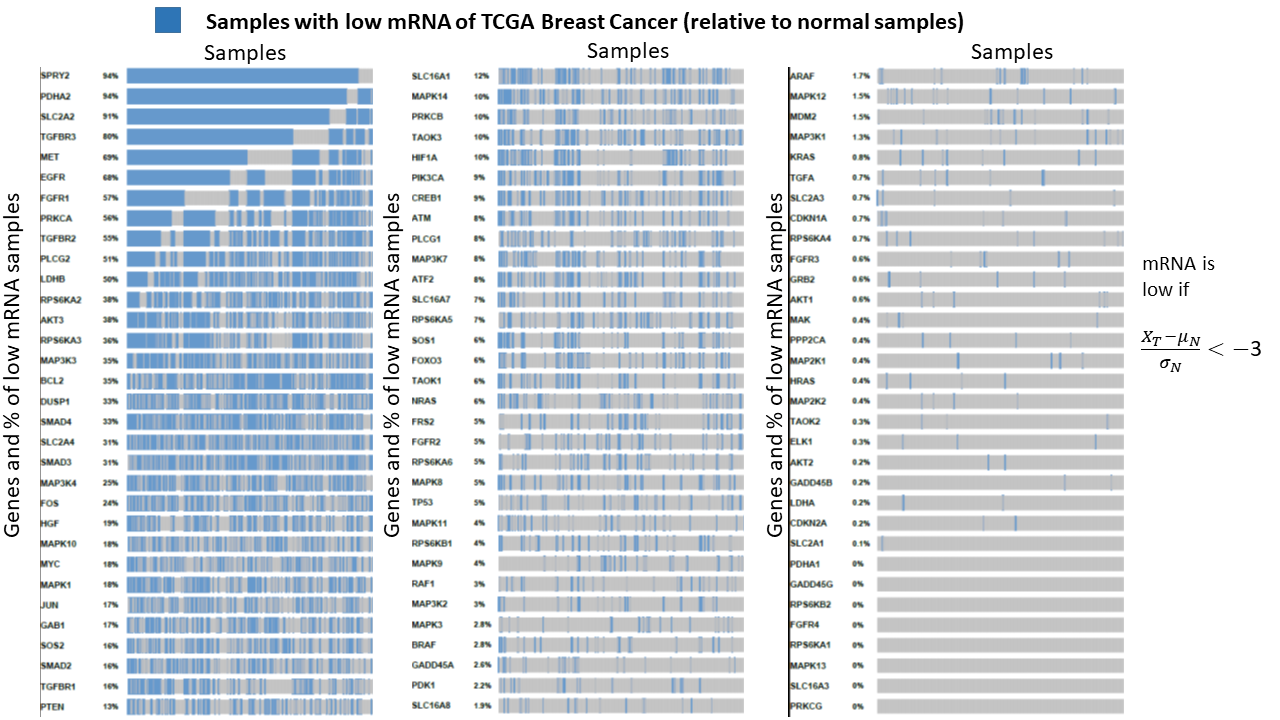

**Fig S5.** **(A)**. TCGA breast cancer samples with high mRNA expression of network genes relative to normal samples. Genes are ordered from high to low of percentage of high mRNA samples. **(B)**. TCGA breast cancer samples with low mRNA expression of network genes relative to normal samples. Genes are ordered from high to low of percentage of low mRNA samples. The horizontal axis shows samples with high (red) or low (blue) expressions of respective genes. We considered the gene as over-expressed when the standard deviation was above +3 with respect to normal samples and under-expressed as when it was below -3.

**A**

Correlation coefficient for TCGA breast cancer mRNA data

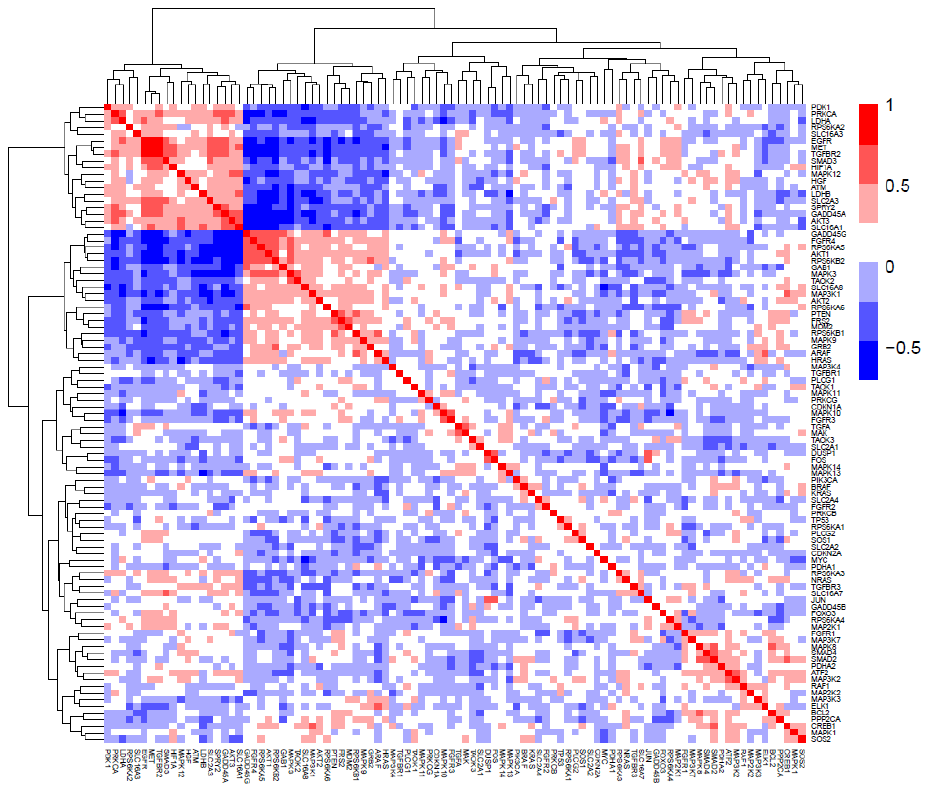

**B**

Correlation coefficient for CCLE breast cancer mRNA data

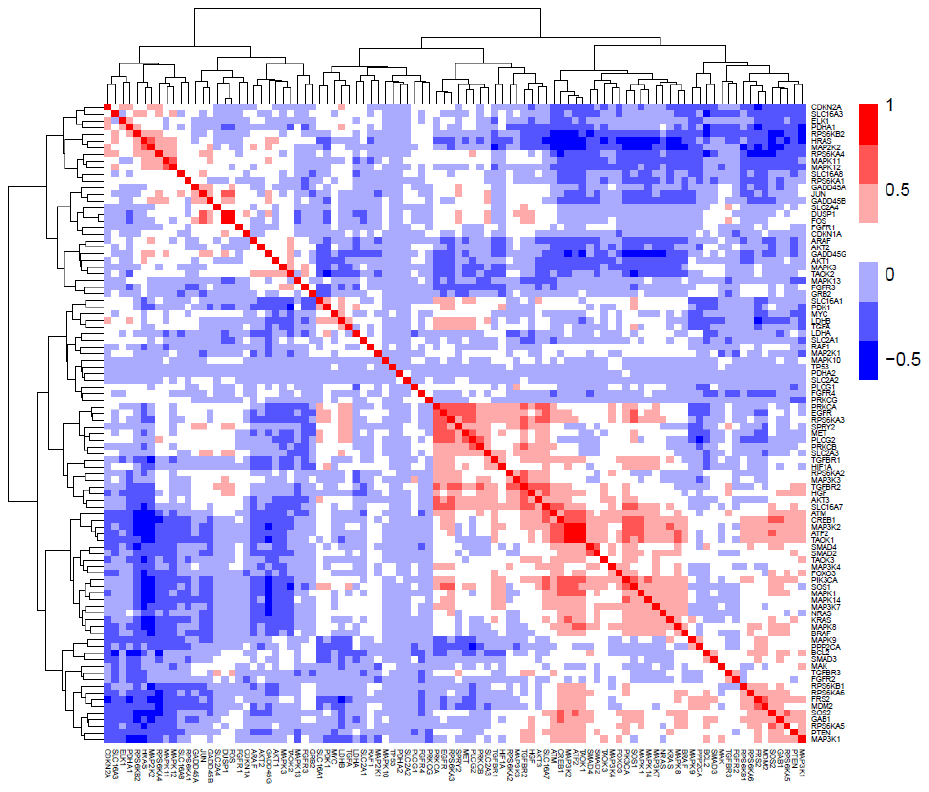

**C**

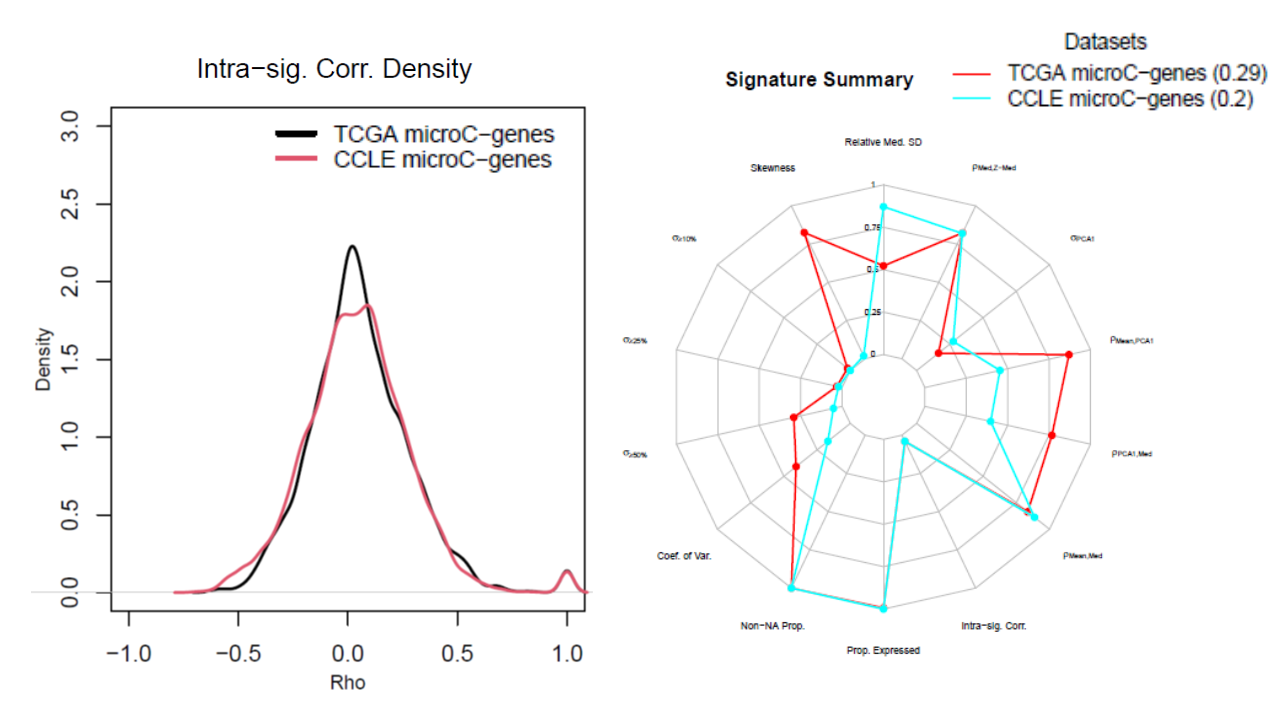

**Fig S6.** sigQC metrics were calculated for our network genes in TCGA and CCLE breast cancer RNA Seq. datasets: **(A).** Correlation coefficients between genes are shown for TCGA data. **(B).** Correlation coefficients between genes are shown for CCLE data. **(C).** Distribution of correlation coefficients and comparison of sigQC metrics calculated for genes using TCGA and CCLE datasets. The distribution of the metrics are fairly similar between two datasets.

**A**
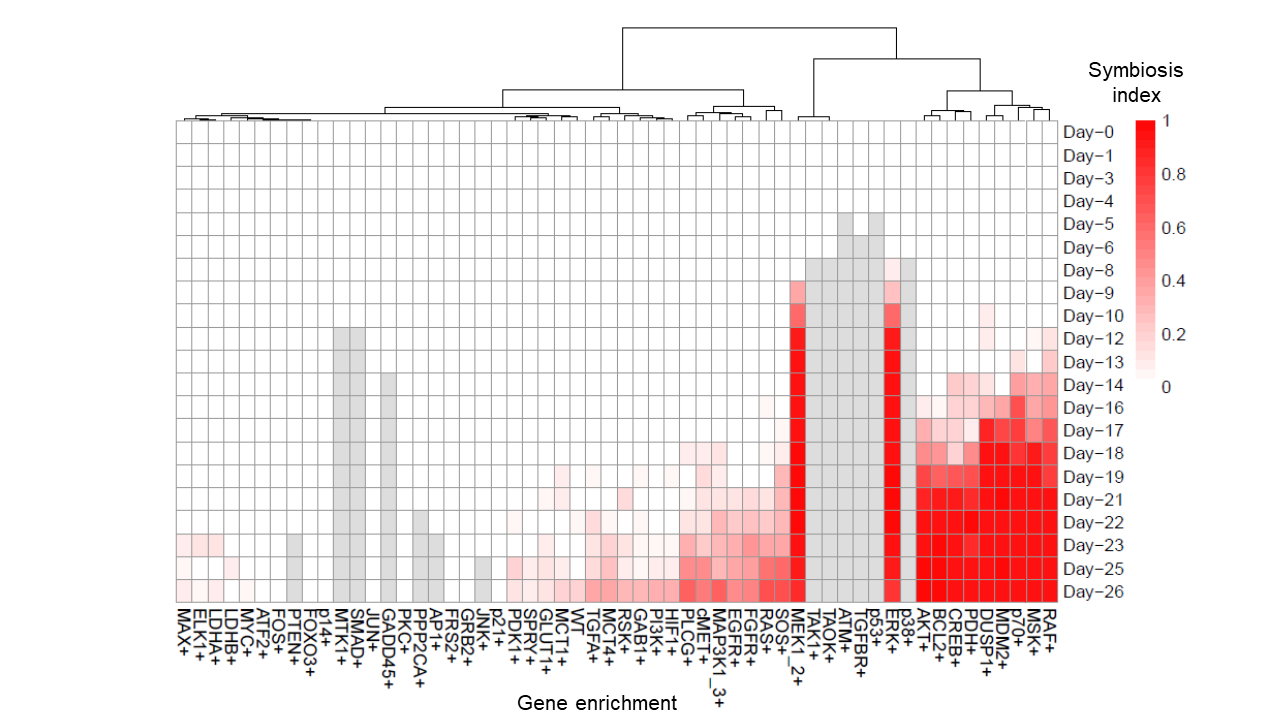

**B**

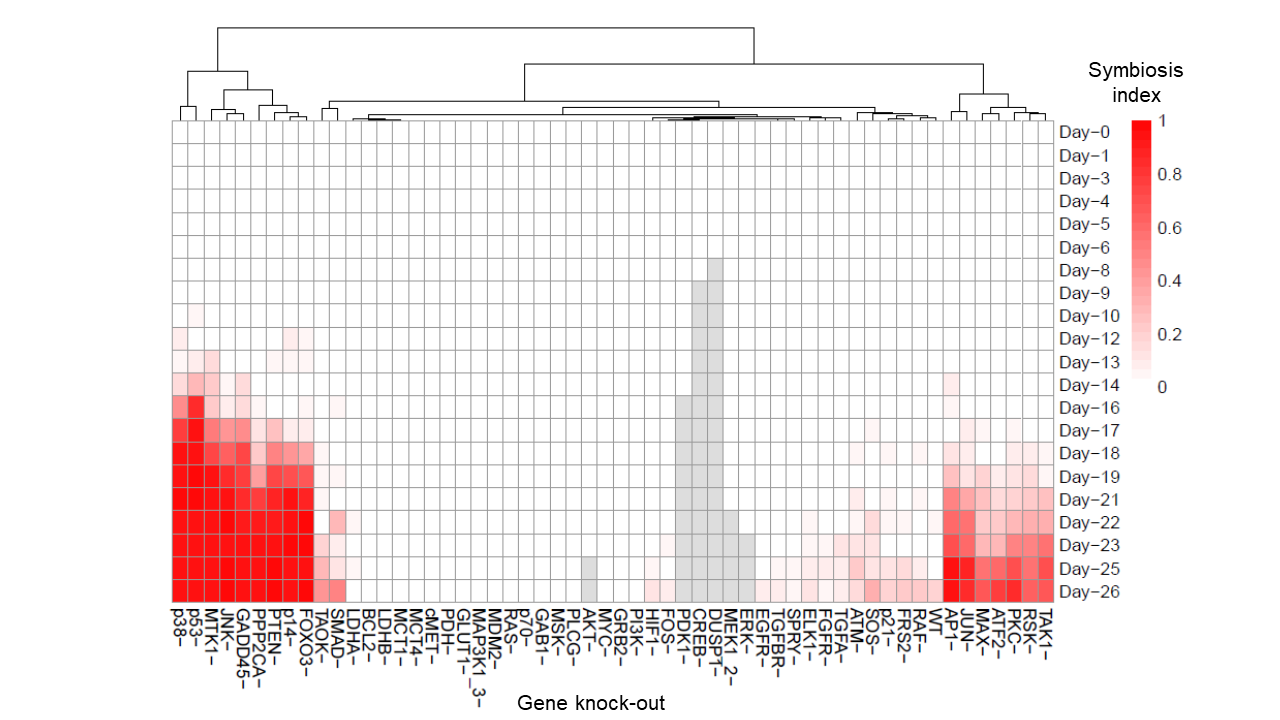

**C**

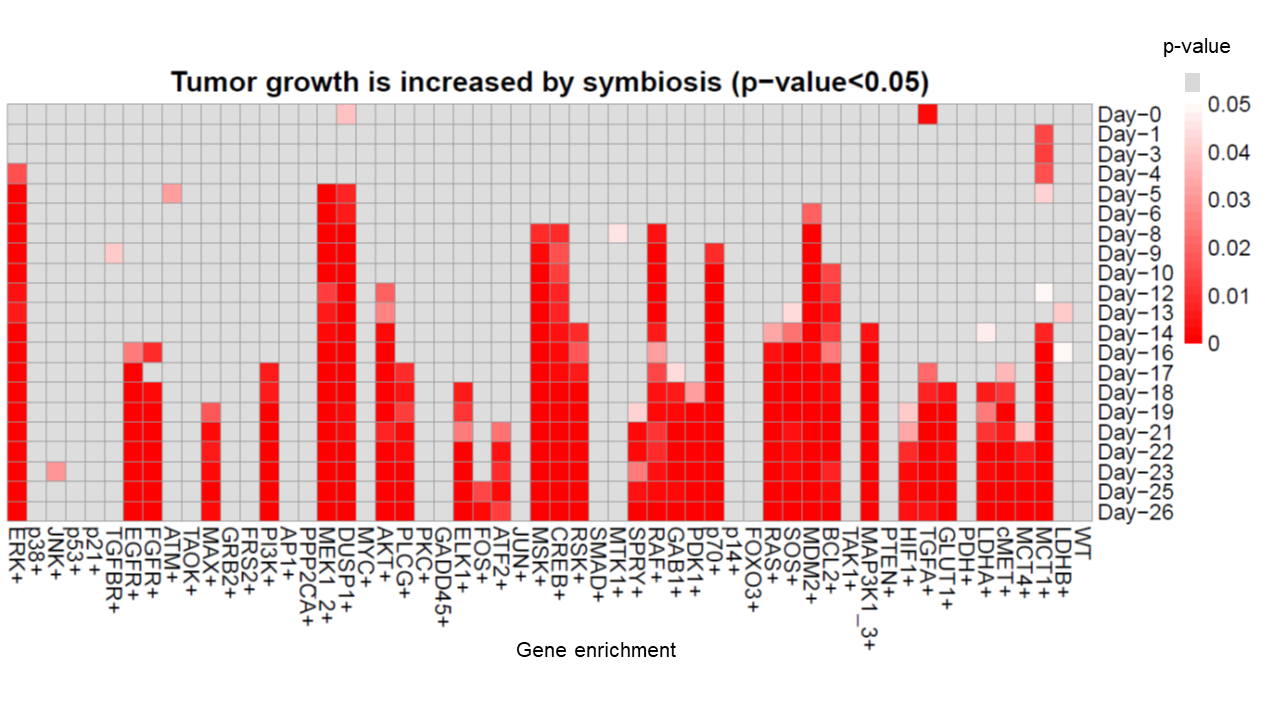

**D**

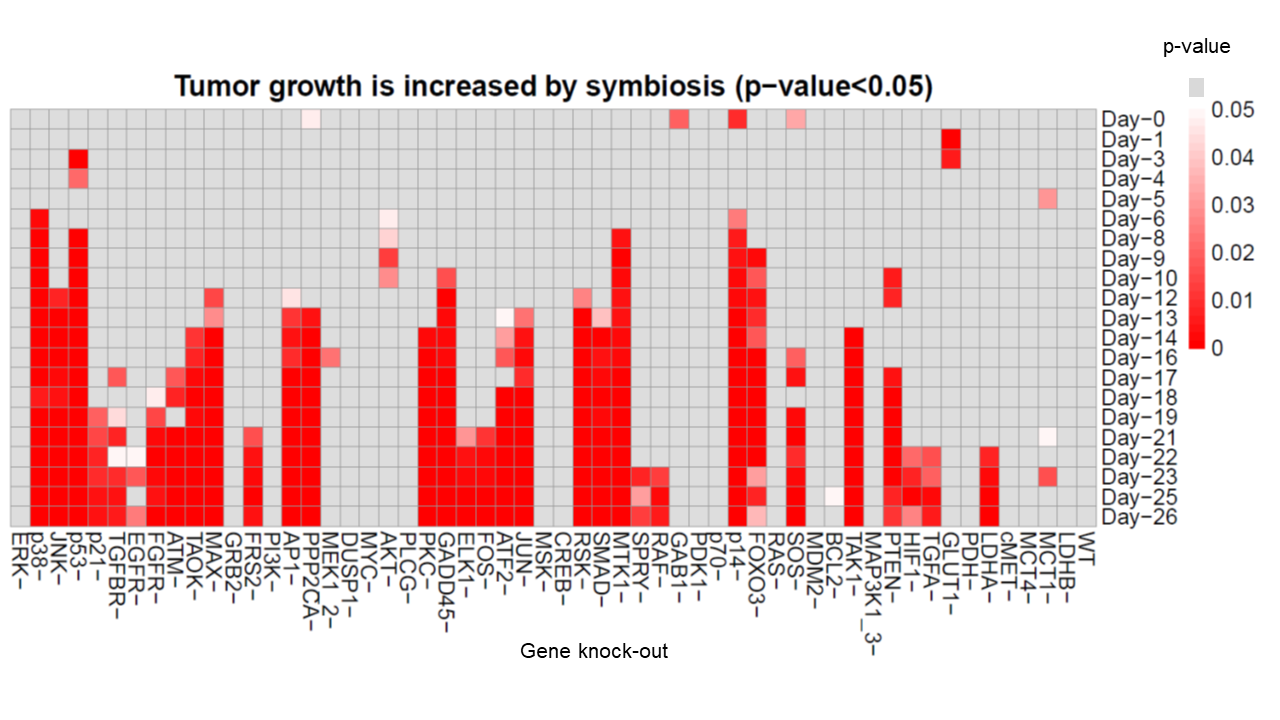

**E**

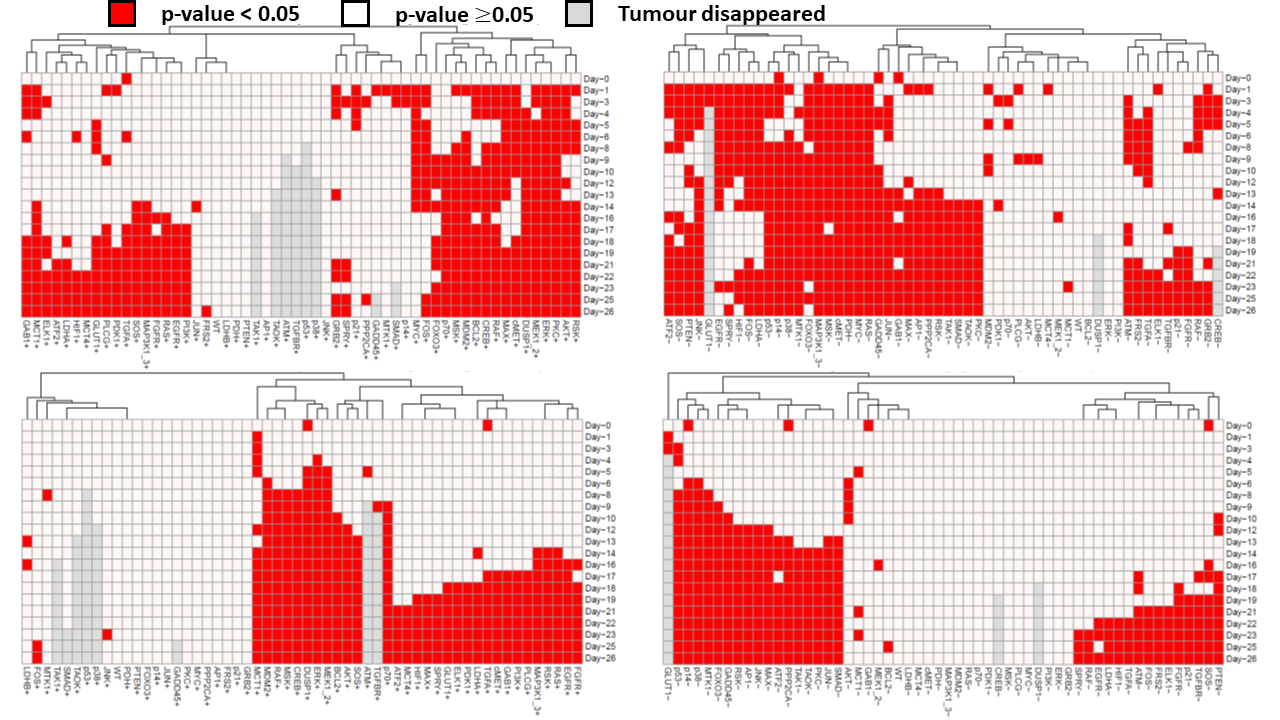

**Fig S7.** Metabolic symbiotic simulations with network gene alterations (Enriched (+) and Knockout (-) status) and wild type (WT) status: **(A)**. Temporal variation of symbiosis index at each gene enrichment status. **(B)**. Temporal variation of symbiosis index at each gene knockout status. **(C)**. Whether symbiosis-induced growth is greater than non-symbiotic tumour growth with each gene enrichment is shown. **(D)**. Whether symbiosis-induced growth is greater than non-symbiotic tumour growth with each gene knockout status is shown. The results show that clusters of gene alterations can be identified, which enhance symbiosis while some other gene alterations reduce symbiosis (A, B). Colors indicate symbiosis index (A, B) and p values (C, D). p values from 0 to 0.05 are shown in red to white color scale and p values $\geq$ 0.05 are shown in grey color. **(E)**. Genes are clustered based on p-value is less than 0.05 (red) or not (white) of the results shown in Fig 4C (top-left), Fig 4D (top-right), Fig S4C (bottom-left), and Fig S4D (bottom-right).

**A**

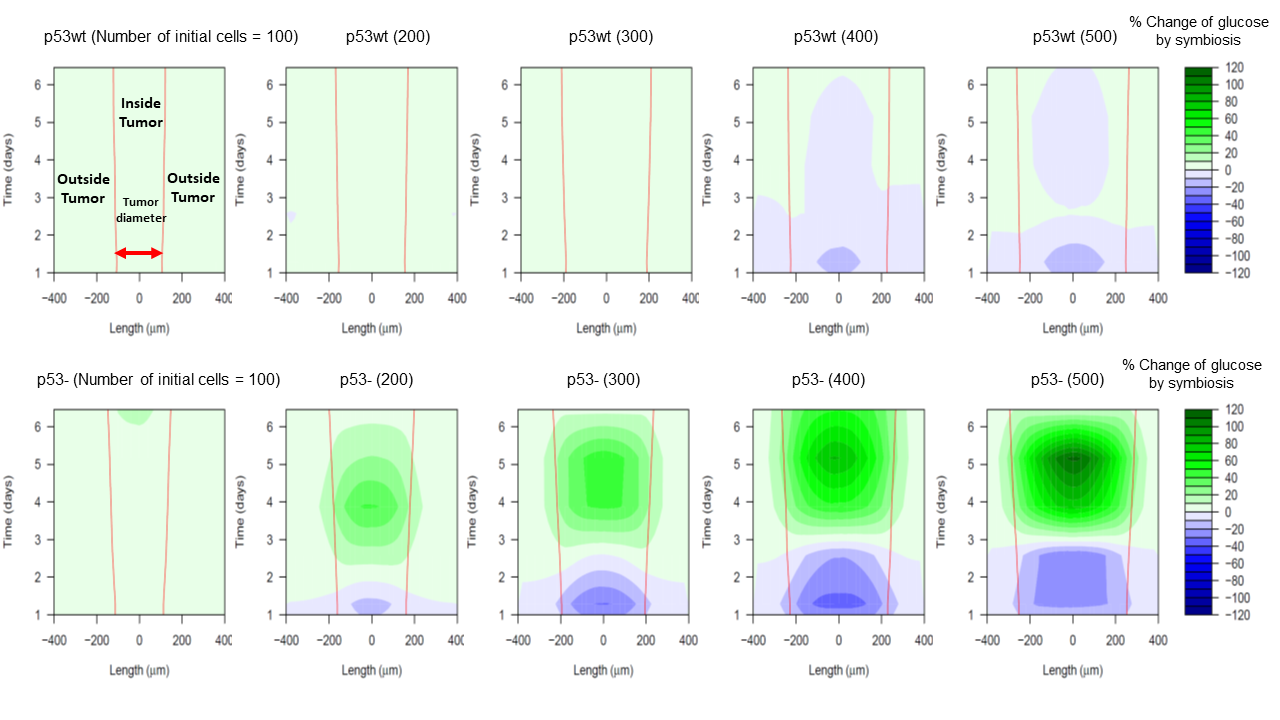

**B**

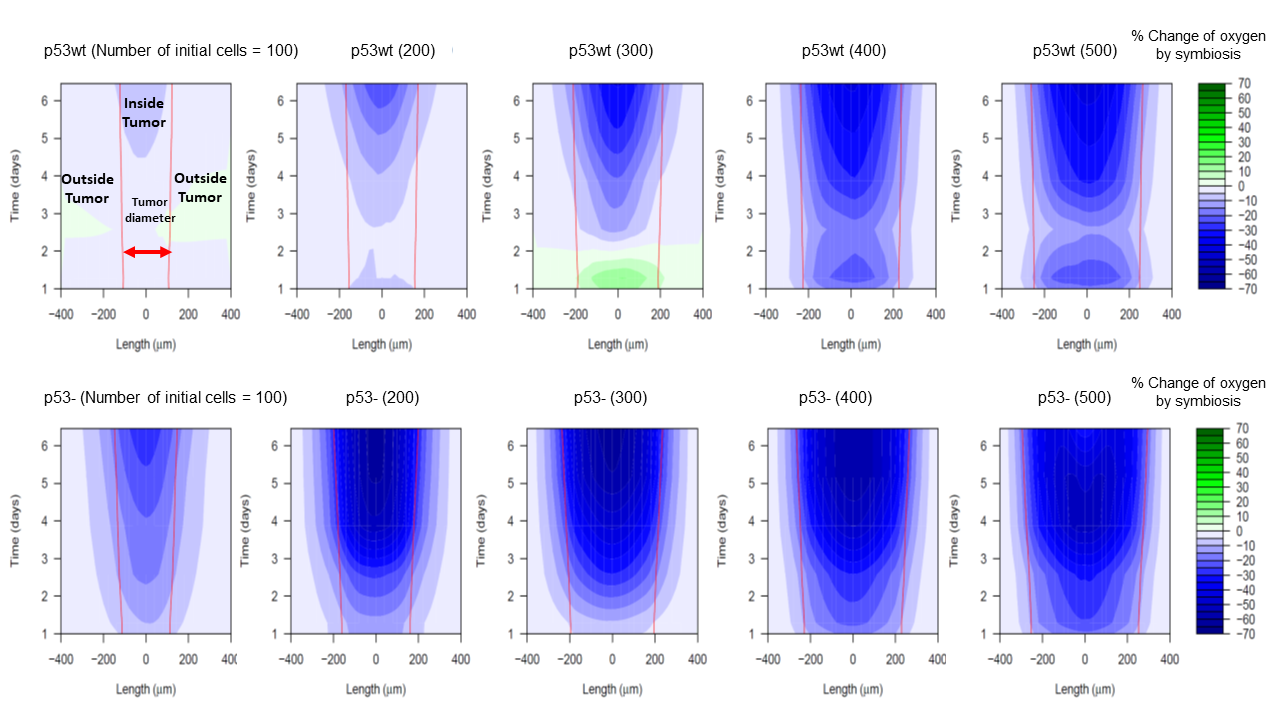

**Fig S8.** Symbiosis-induced change of glucose and oxygen of the microenvironment over time is shown here (here, the Length is the cross section through the center of the tumour). The heat maps show the variation of the percentage change of oxygen and glucose due to metabolic symbiosis under p53wt and p53- status, and at different initial tumour sizes: **(A)**. The symbiosis would increase the glucose level in the medium. **(B)**. The symbiosis would decrease the oxygen level in the medium.

**A**

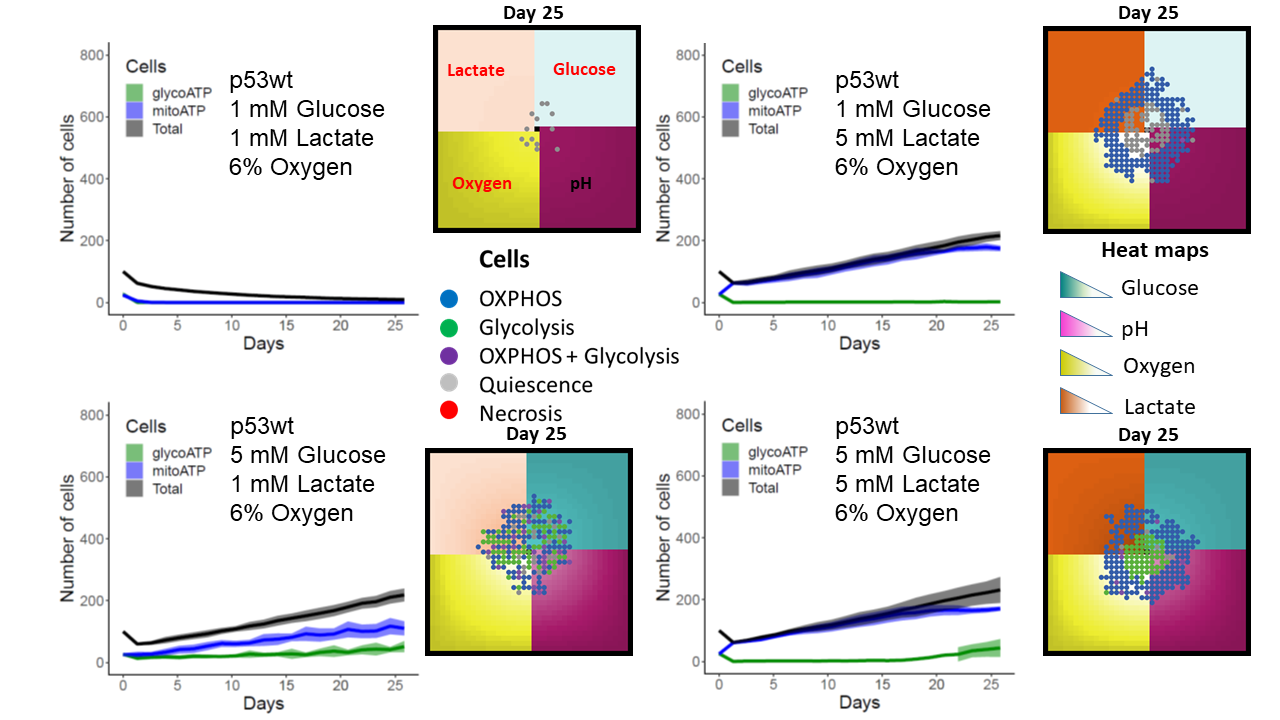

**B**

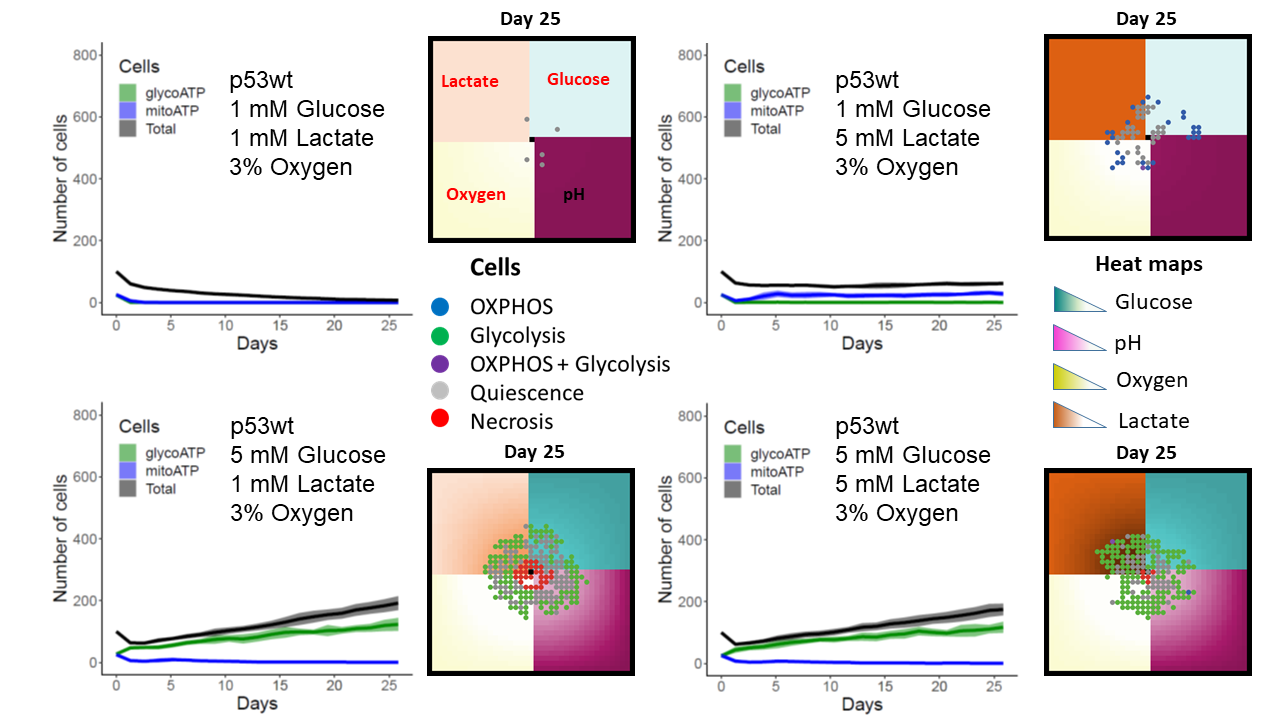

**C**

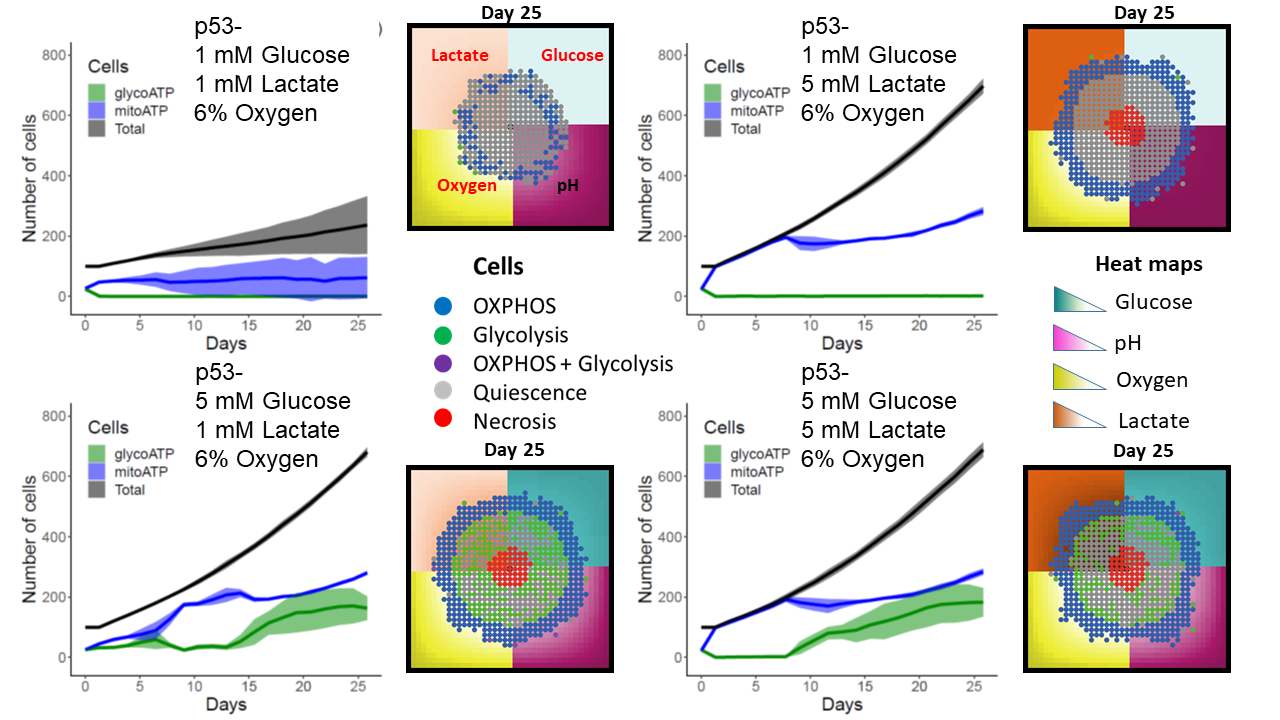

**D**

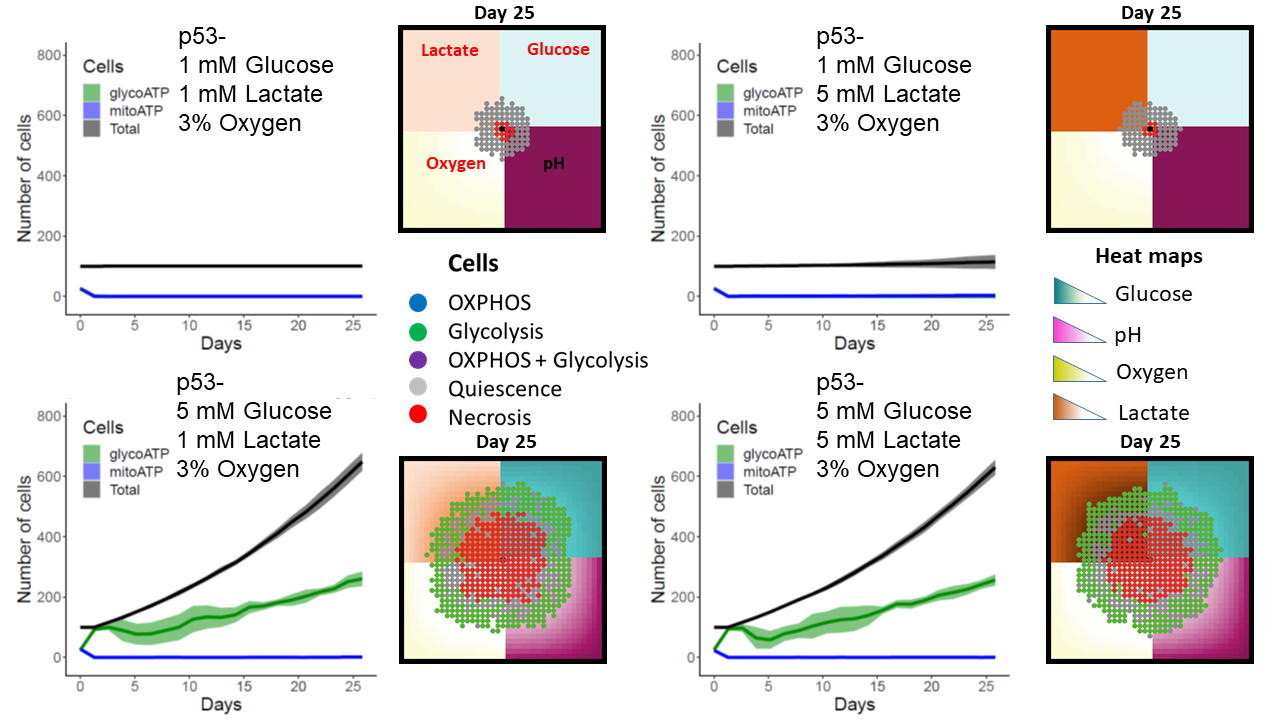

**Fig S9.** Tumour growth under different environmental conditions (different levels of glucose, lactate and oxygen) are shown for p53wt and p53- cells: **(A)**. p53wt cells with 6% oxygen level. **(B)**. p53wt cells with 3% oxygen level. **(C)**. p53- cells with 6% oxygen level. **(D)**. p53- cells with 3% oxygen level. The results show that both glucose and lactate metabolism can fuel tumour growth when tumour is well-oxygenated.

**A**

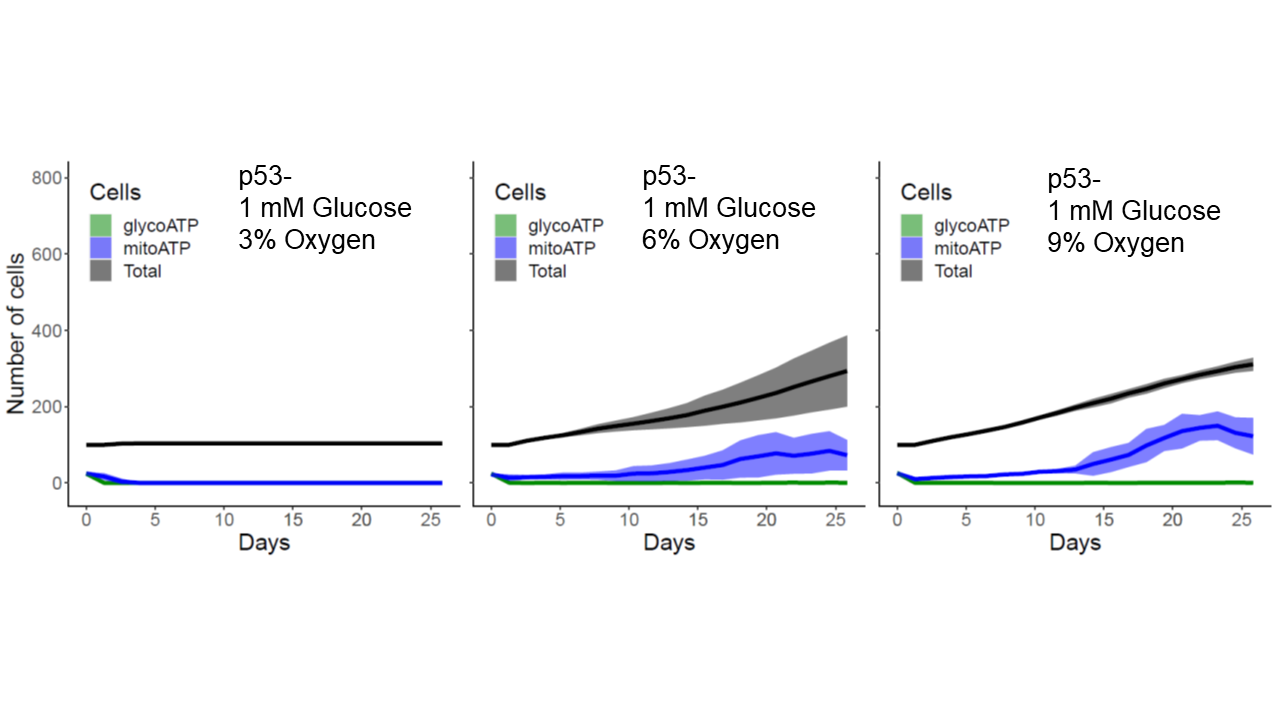

**B**

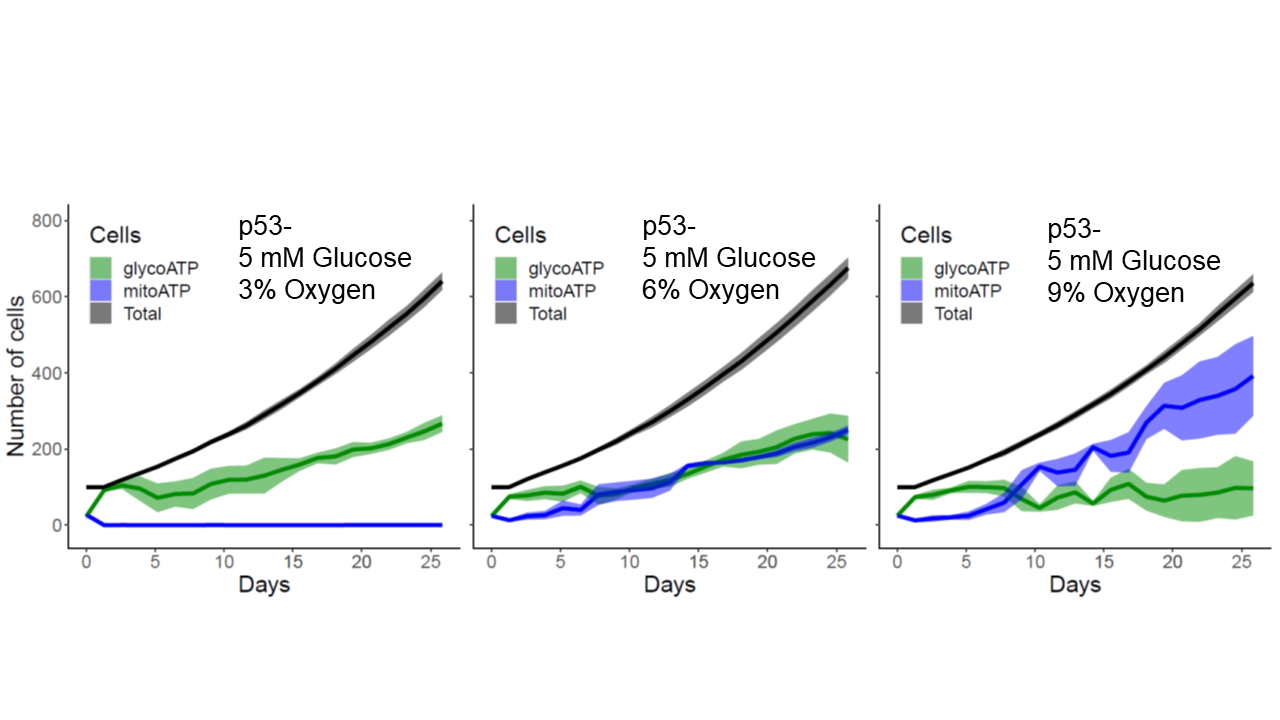

**C**

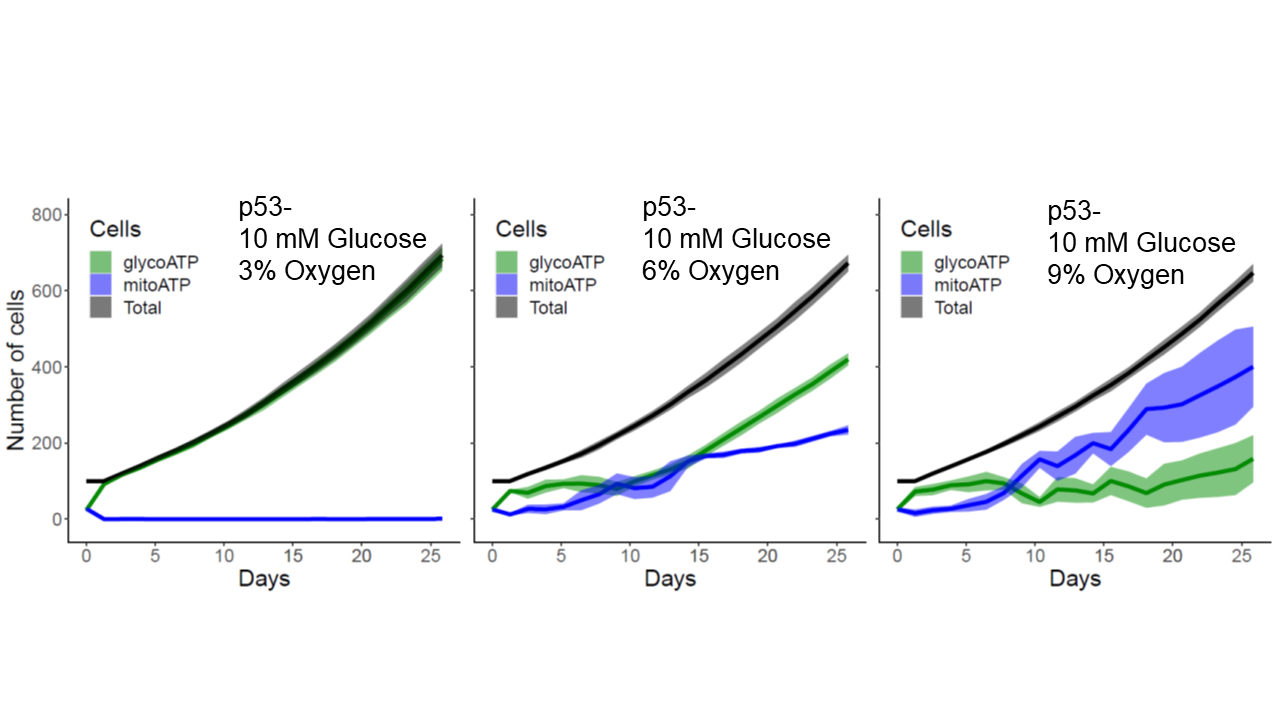

**Fig S10.** Tumour growth curves are shown for different combinations of oxygen and glucose. As the oxygen level is increased, more tumour cells switch to OXPHOS because they have both glucose and lactate as the energy source for mitochondrial ATP production. The oxygen level at the boundary of the simulation domain (square box) was maintained at 3%, 6%, and 9% O2 while the glucose level was at 1 mM (A), 5 mM (B) and 10 mM (C). Note that oxygen and glucose levels in the tumour were much lower than these boundary values.

**A**

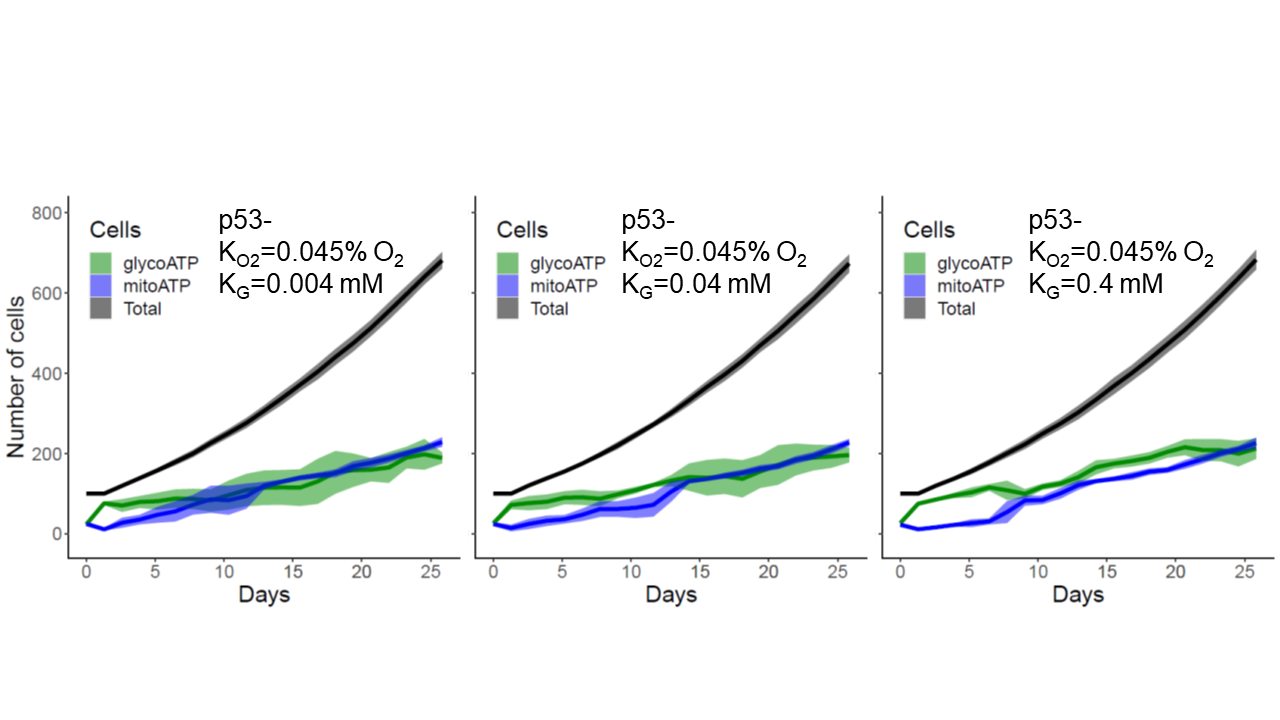

**B**

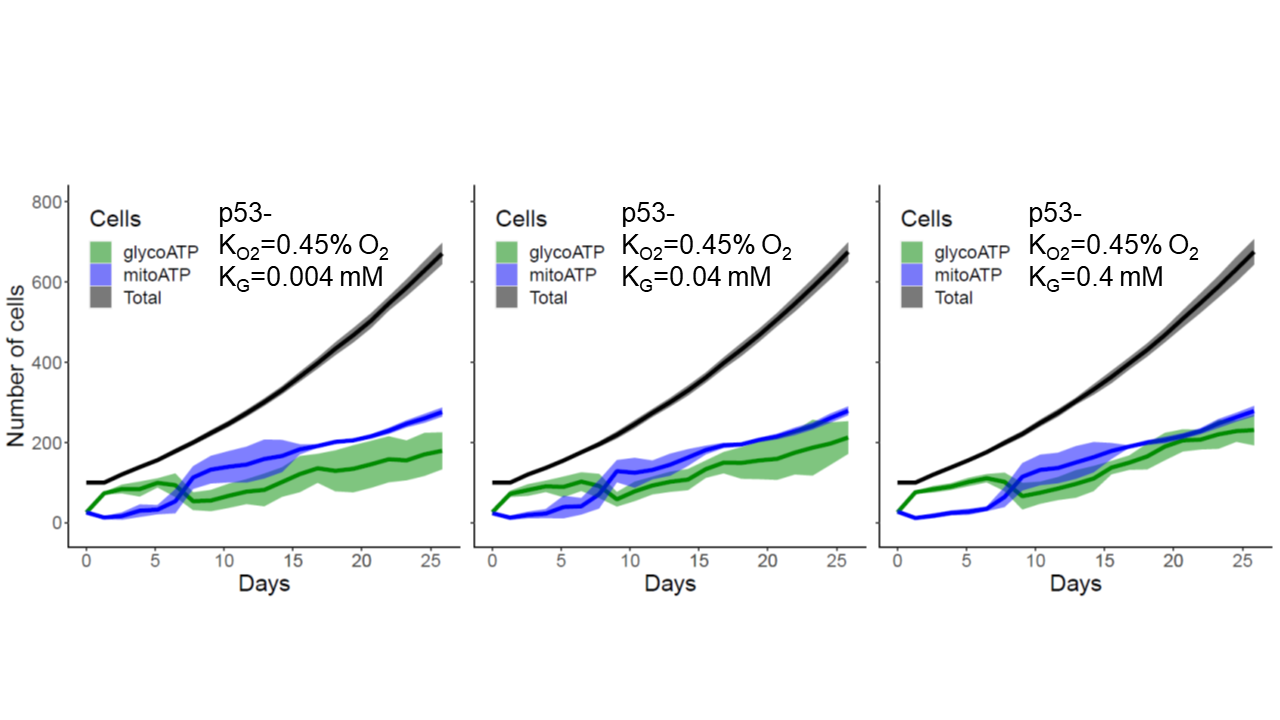

**C**

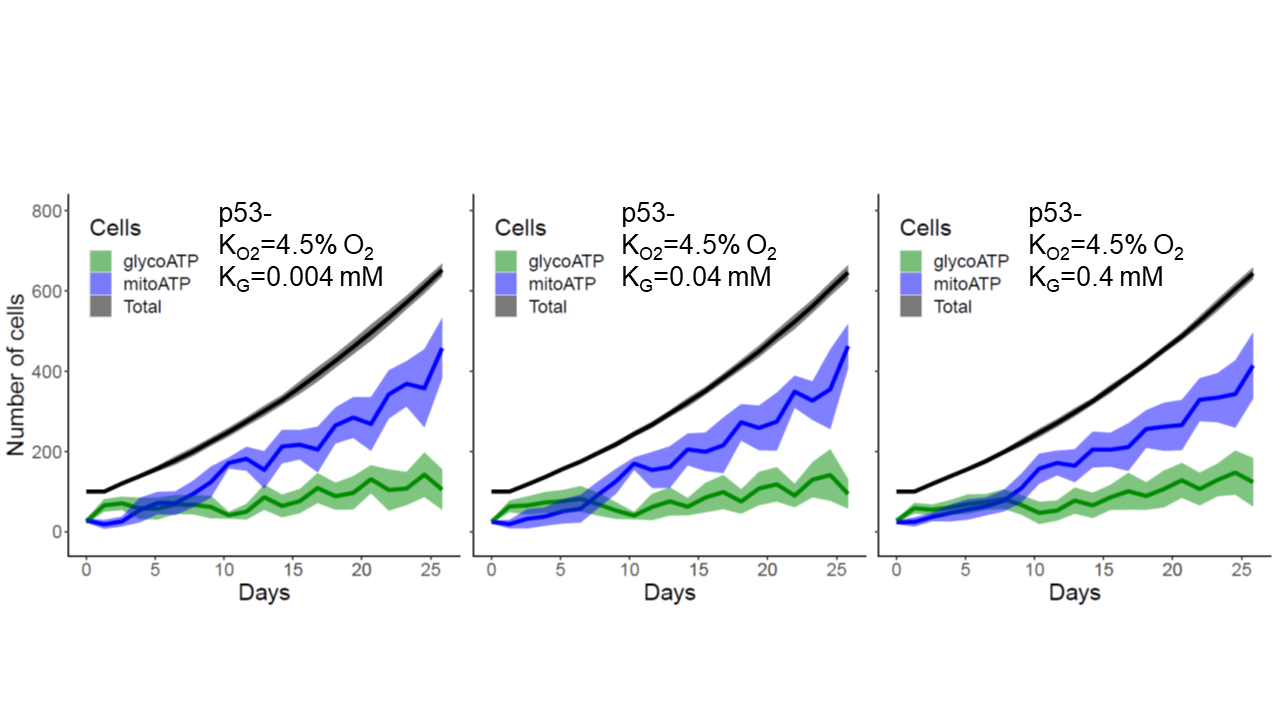

**Fig S11.** The effect of half-saturation coefficients of oxygen and glucose on the metabolic pathways is shown. The half-saturation coefficient of oxygen was kept at 0.045% oxygen (A), 0.45% oxygen (B) and 4.5% oxygen (C) while the half saturation coefficient of glucose was varied to 0.004 mM, 0.04 mM and 0.4 mM. The pathways seem more sensitive to the variation of the half-saturation coefficient of oxygen.

**A**

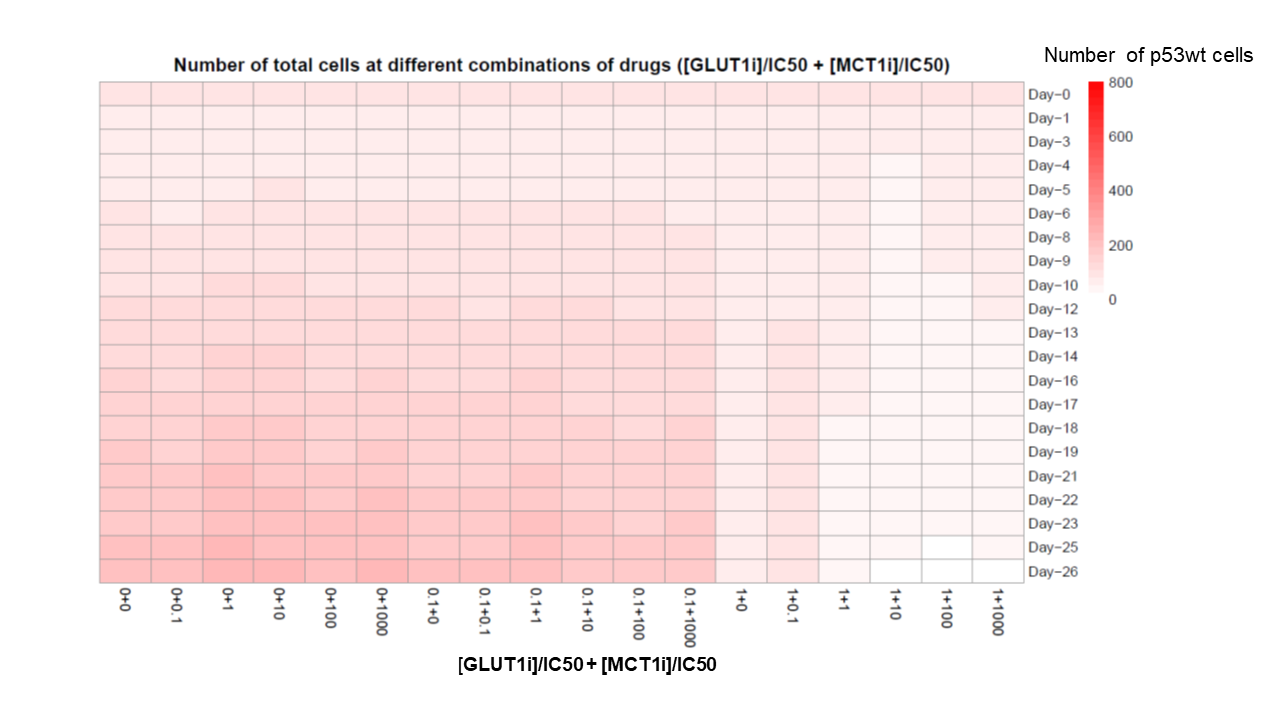

**B**

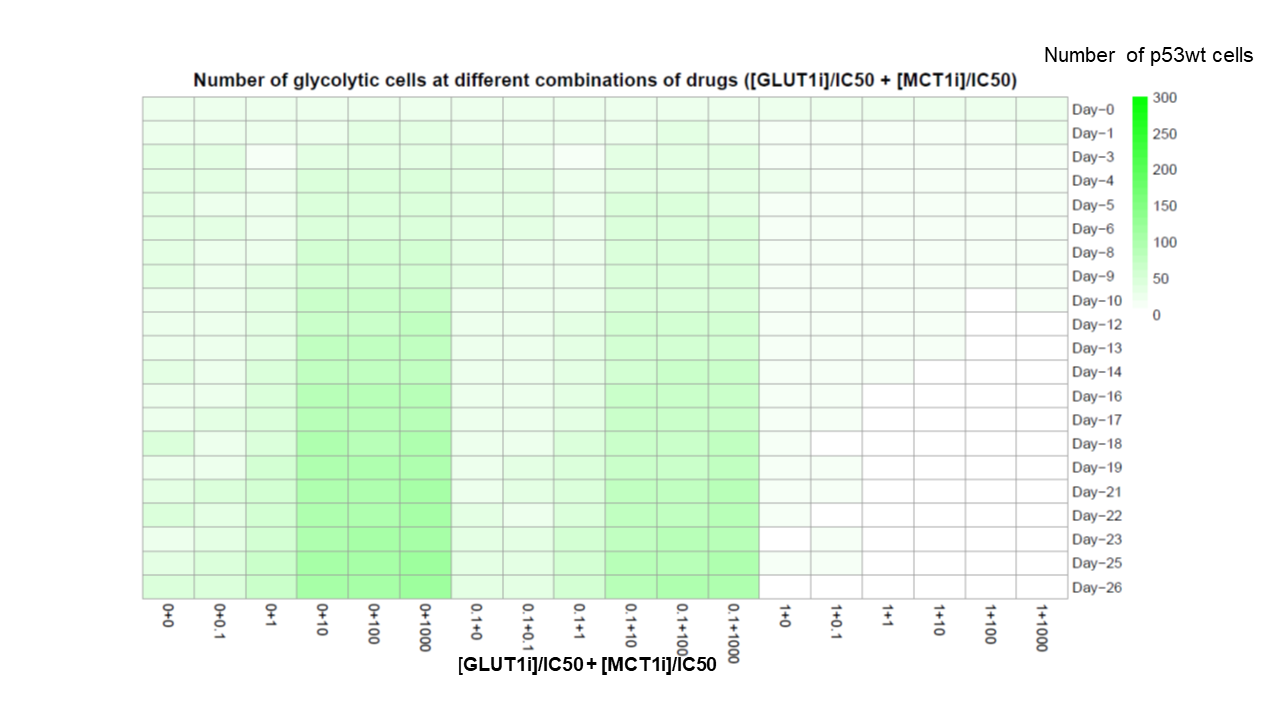

**C**

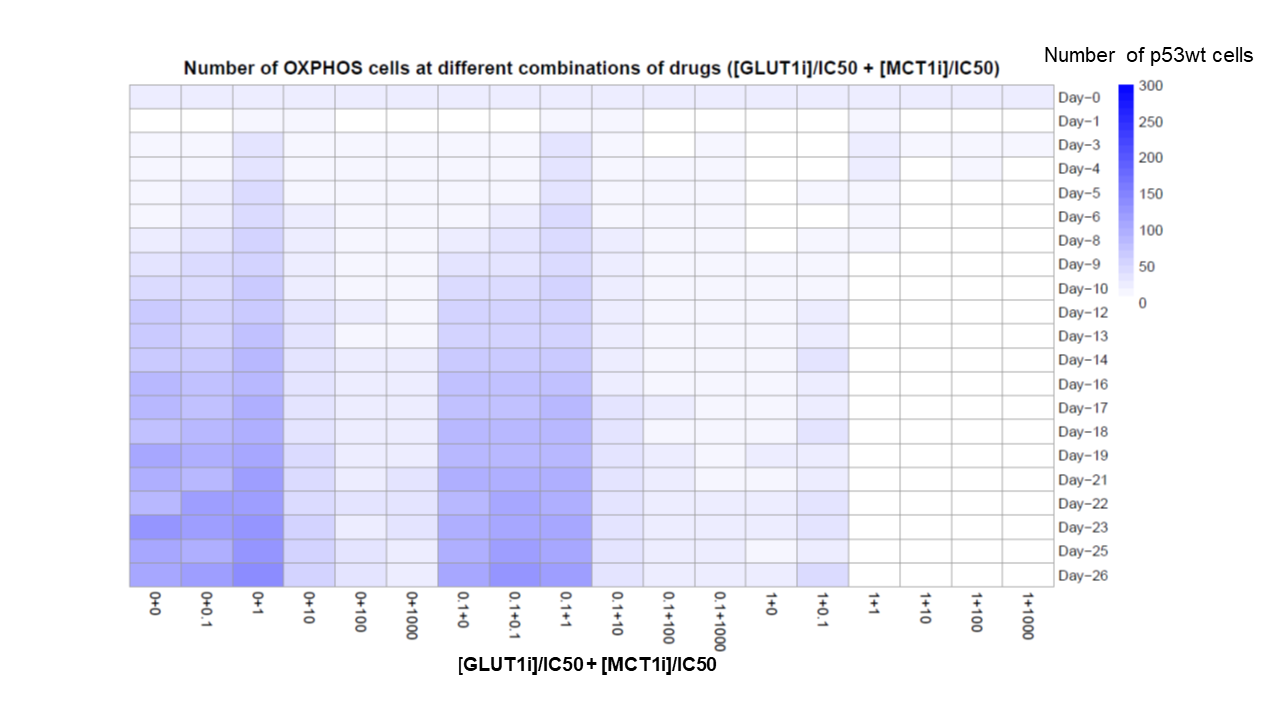

**D**

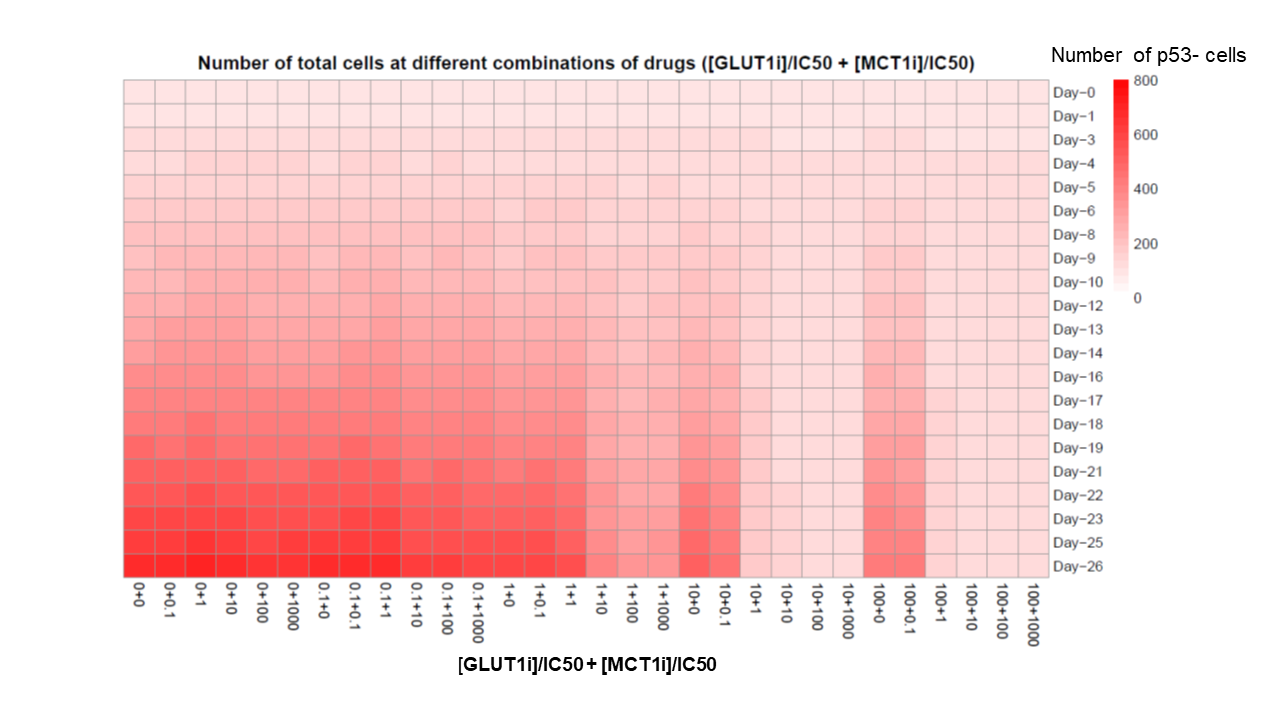

**E**

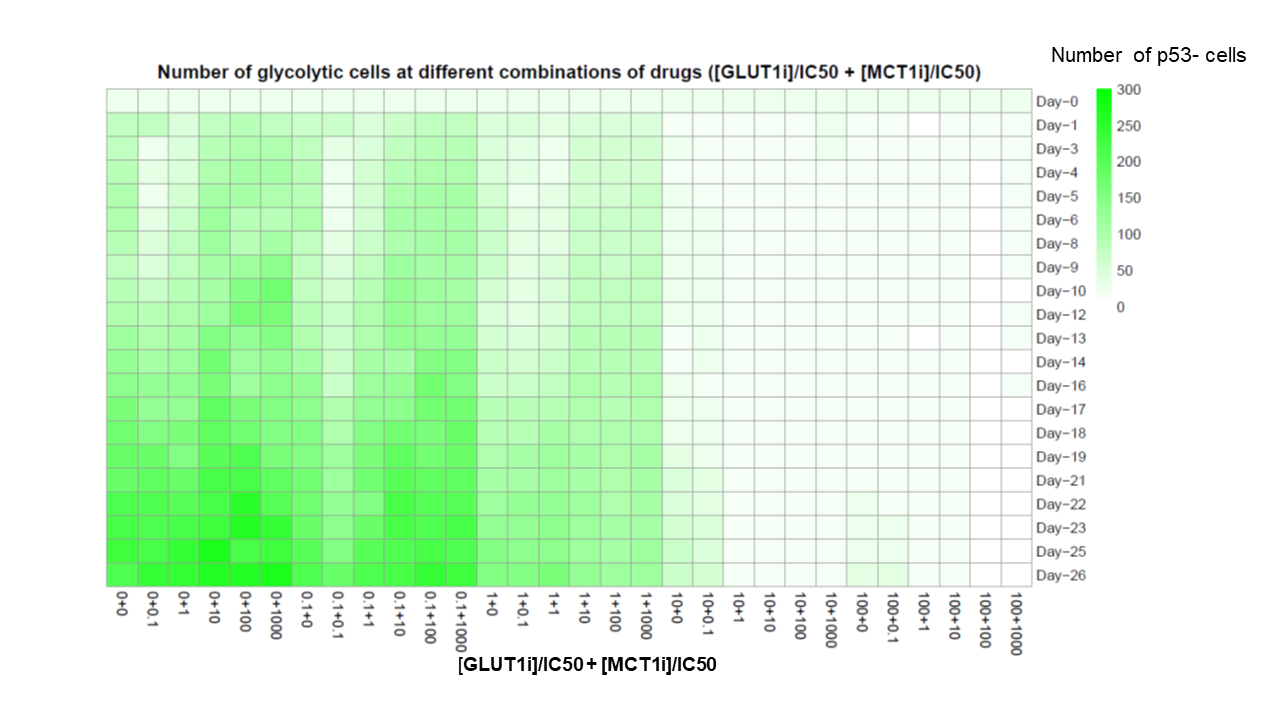

**F**

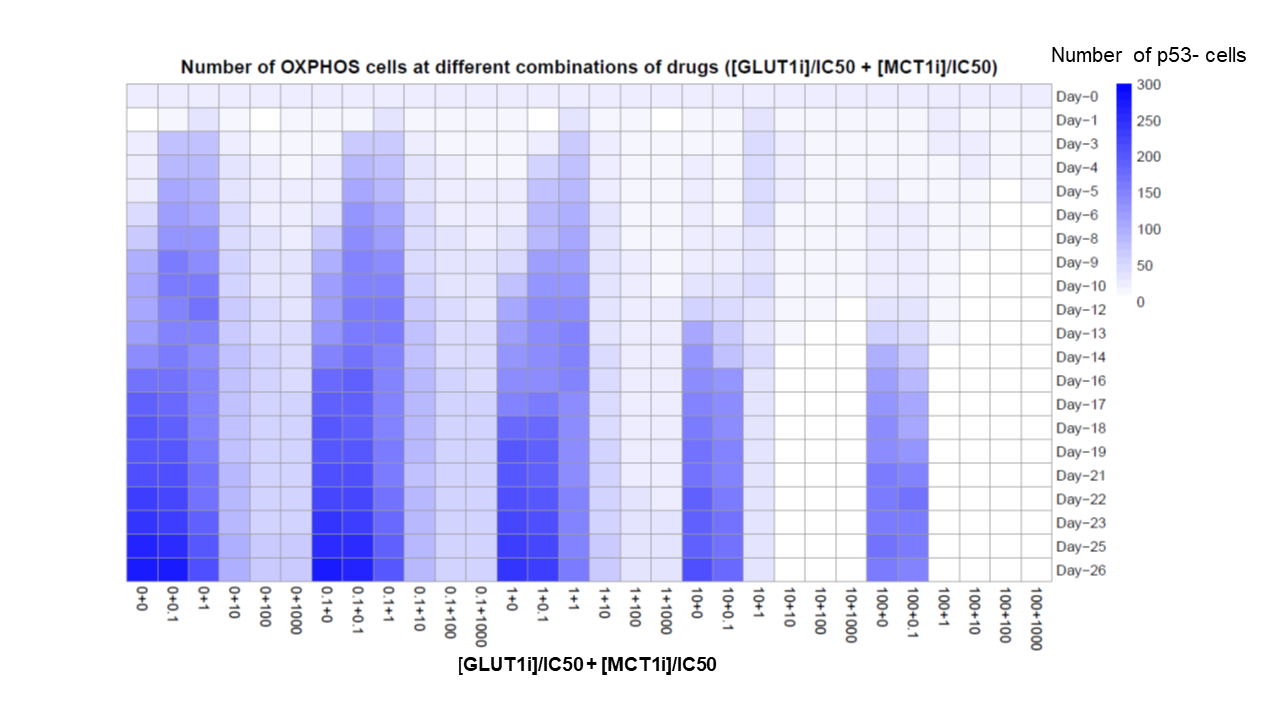

**Fig S12.** Tumour growth over time at different combinations of GLUT1 and MCT1 inhibitors. [GLUT1i]/IC50 and [MCT1i]/IC50 were varied from 0 to 100 and 0 to 1000, respectively: **(A, B, C)**. p53wt tumour cells. Note that the tumour is completely disappeared at [GLUT1]/IC50 = 10. **(D, E, F)**. p53- tumour cells. Temporal variations of total cells (A, D), glycolytic cells (B, E), and OXPHOS cells (C, F) are shown.

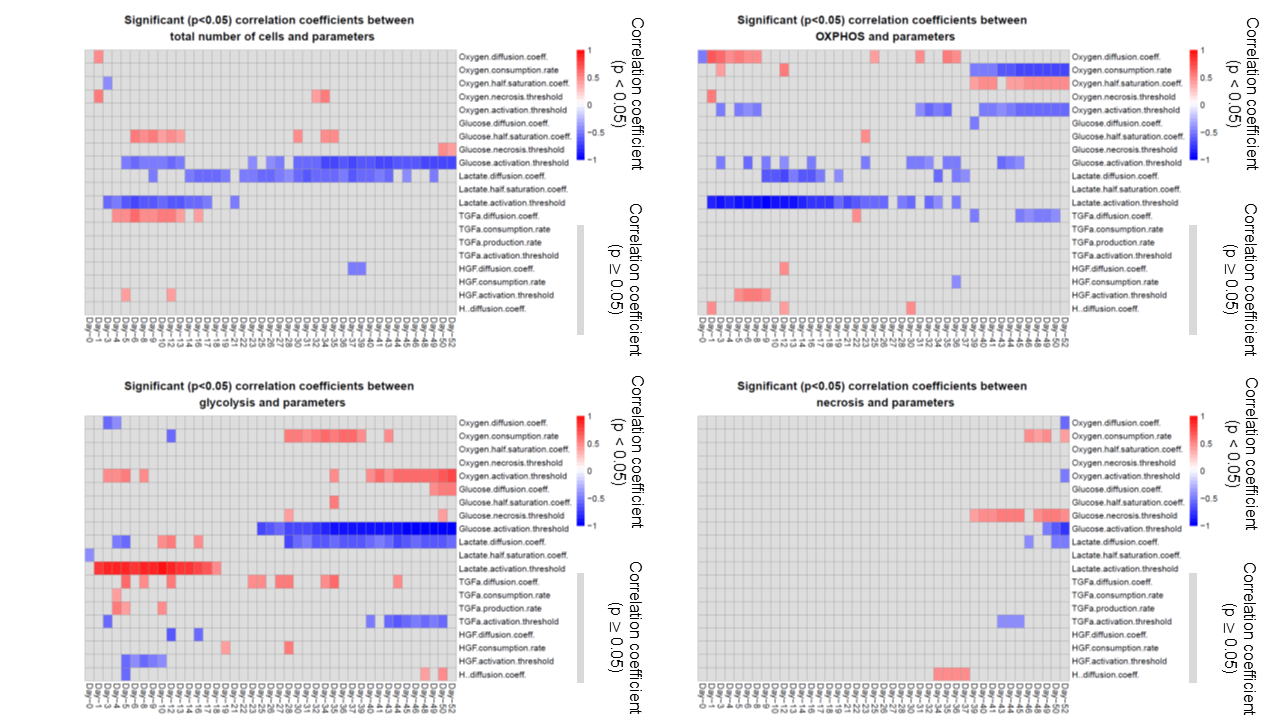

**Fig S13.** Partial correlation coefficients between model parameters and outputs over time. The model outputs are total number of cells, OXPHOS cell, glycolytic cell, and necrotic cell populations. The outputs have significant correlations with some parameters in a time dependent manner.

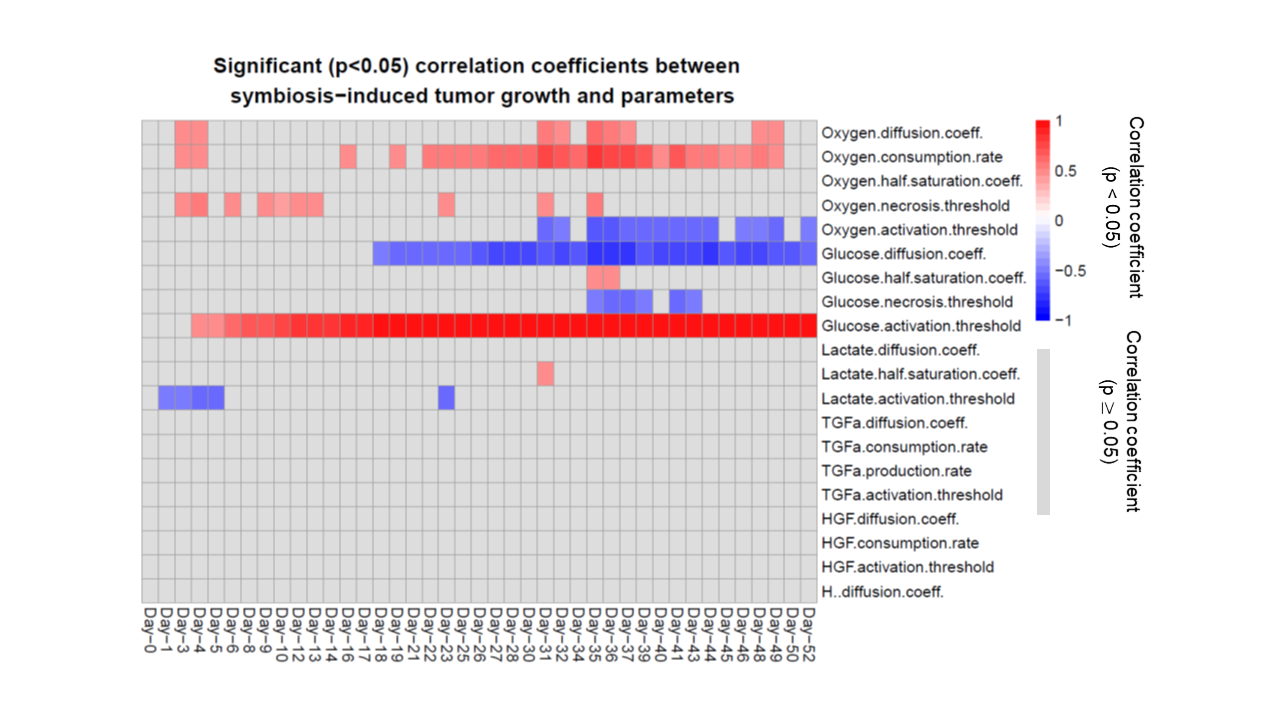

**Fig S14.** Partial correlation coefficients between symbiosis-induced growth increment of tumour and model parameters over time. Glucose activation threshold and oxygen consumption rate have strong positive correlations with metabolic symbiosis while glucose diffusion coefficient and oxygen activation threshold have negative correlations with symbiosis.

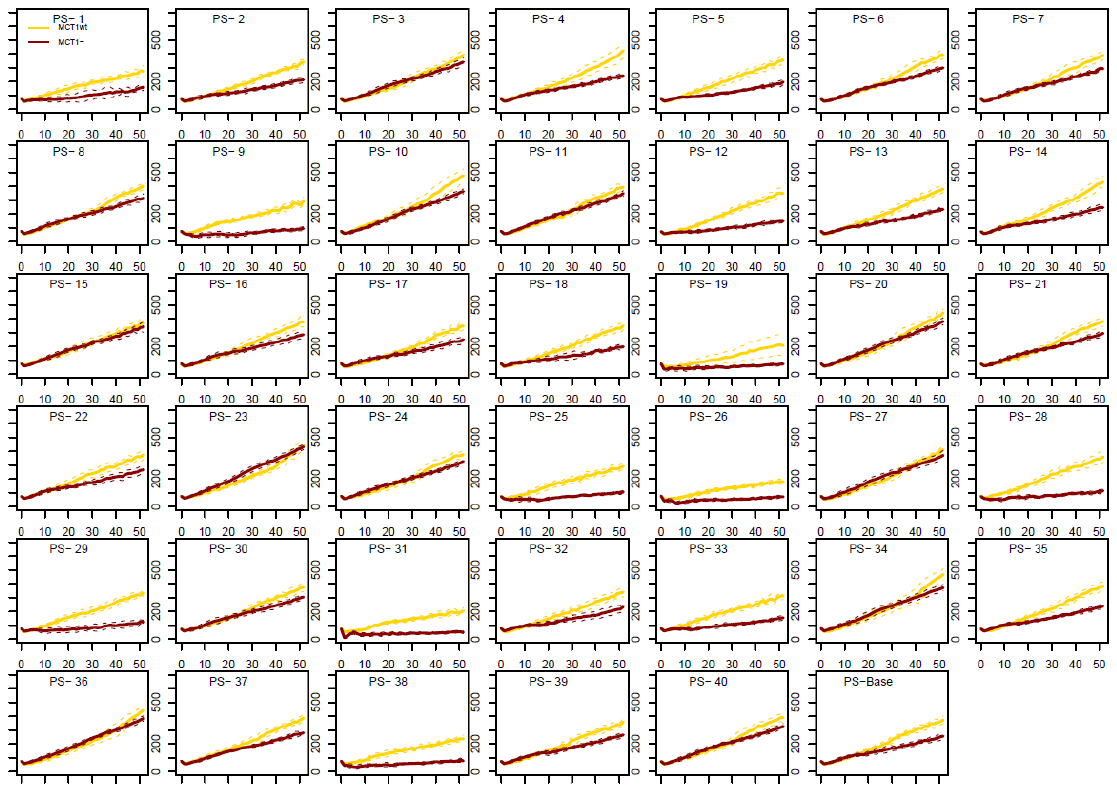

**A**

**B**

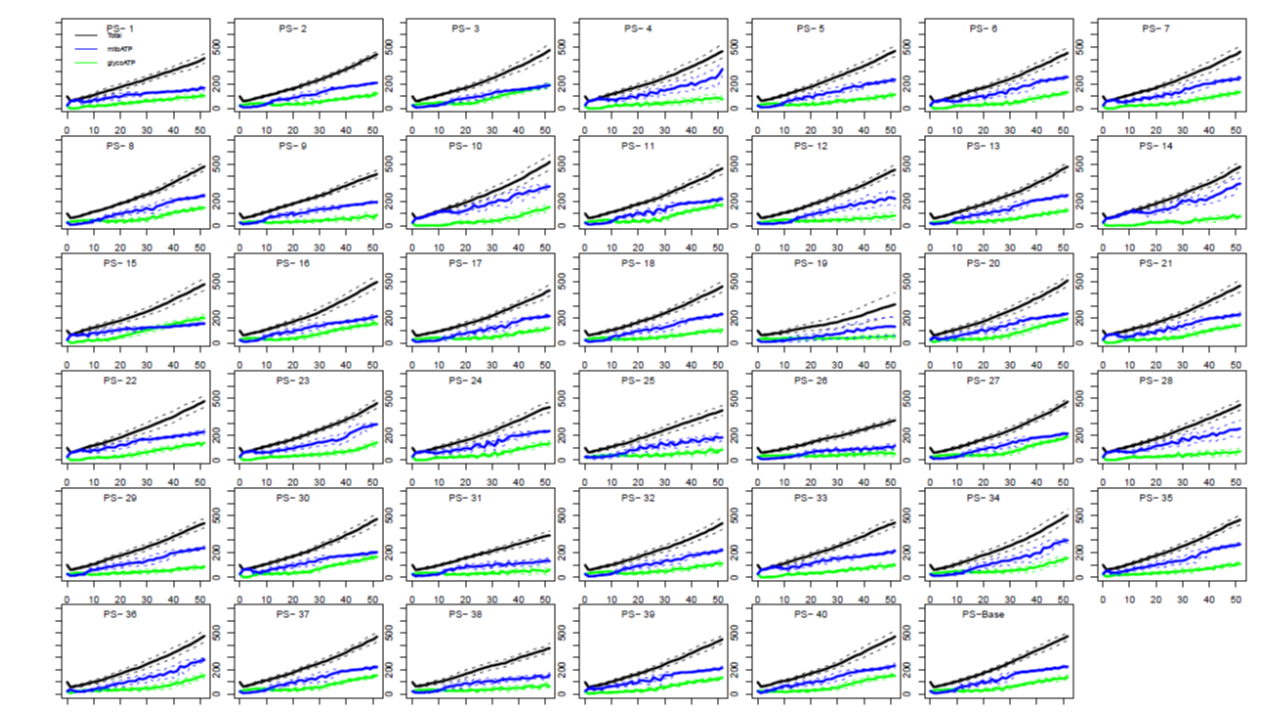

**Fig S15.** **(A)**. Number of active tumour cells with symbiosis (MCT1wt) and without symbiosis (MCT1-) for each perturbed parameter set (PS1-40) and the baseline parameter set (PS-Base). **(B)**. Number of total cells, and mitochondrial and glycolytic ATP producing cells obtained at each parameter set.
