## Supplementary figures and images for "Metabolic Symbiosis between Oxygenated and Hypoxic Tumour Cells: An Agent-based Modelling Study"

### S2-File-Cell_Regulatory_Network.png

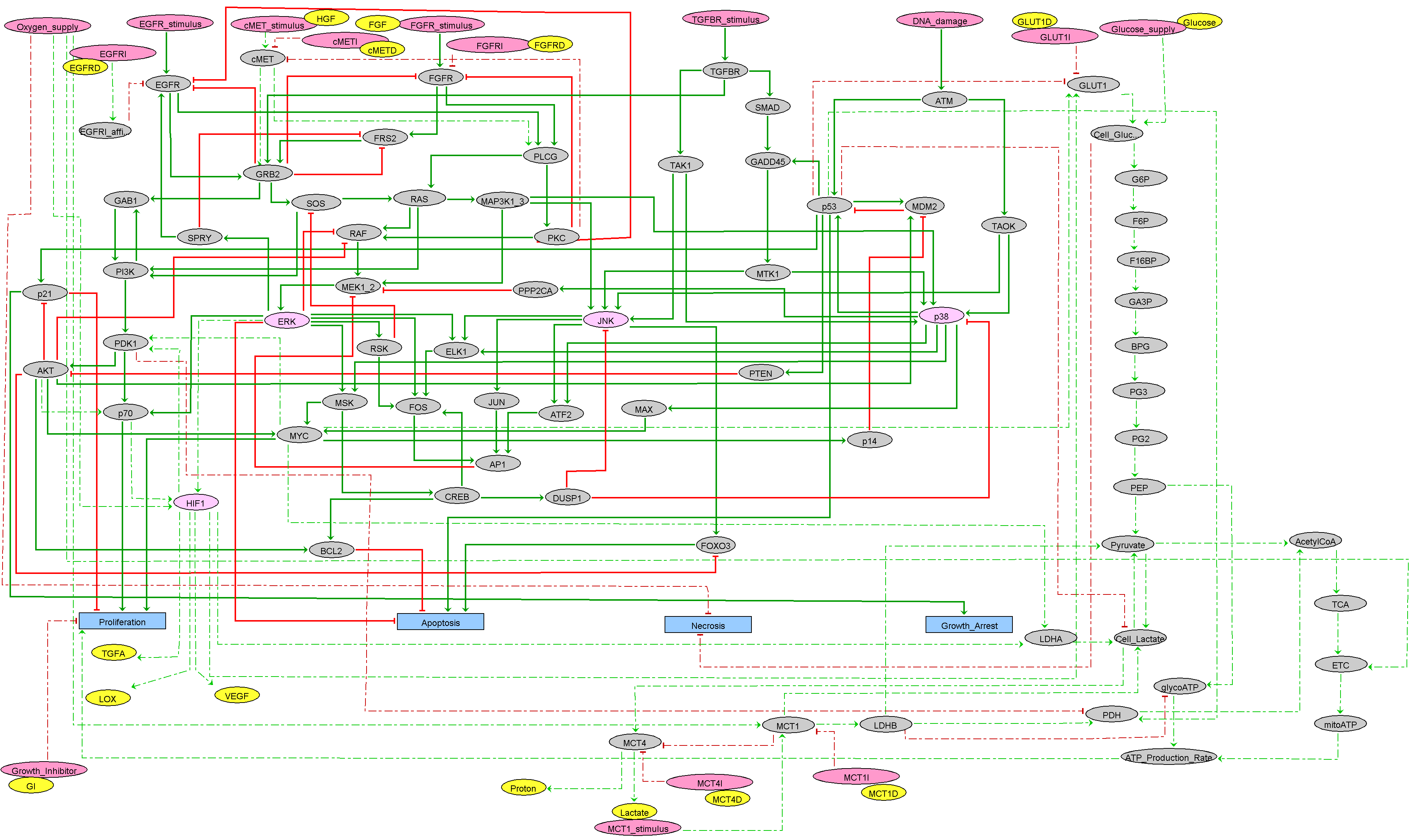
